# Re-evaluating management of established pests including the European wasp, *Vespula germanica* using biocontrol agents

**DOI:** 10.1101/2022.11.22.517291

**Authors:** Oscar Cacho, Susie Hester, Peter Tait, Raelene Kwong, Greg Lefoe, Paul Rutherford, Darren Kriticos

## Abstract

Established pests such as the European wasp (*Vespula germanica*) are often overlooked as candidates for management programmes (eradication and/or containment) because the use of traditional surveillance and control techniques over very large areas becomes uneconomic. Use of biological control agents that persist in the environment is usually the only economically feasible option, however the processes around approvals for release of biocontrol agents can take significant amounts of time and resources, especially if screening and testing of potential agents is required.

This project investigates whether the European wasp could be a candidate for a renewed management programme in south-eastern Australia given the availability of a biocontrol agent following successful screening and testing of an agent, *Sphecophaga vesparum vesparum*, in the 1980s. Whether a biological control programme is worthwhile pursuing depends on the size of the benefits to industry, community and the environment from a reduction in European wasp abundance. This project explores the benefits and costs of European wasp management using a biocontrol agent, and importantly, includes valuation of the social and environmental impacts of the pest.

## 1. Executive Summary

### 1.1 Key findings

Plausible and biologically meaningful estimates of parameter values for the population dynamics of the European wasp and the biocontrol agent, based on NZ studies and available Australian data, allowed the exploration of scenarios in which biological control would succeed in supressing the European wasp. Given the parameter values used in the modelling, the following conclusions may be drawn:

1. **Impacts of European wasps are significant** If European wasps continue to spread across Australia without a formal management programme, total damage over a time period of 50 years could be in the order of $2.66 billion in present value terms. More than half of this is due to the damage that wasps cause to the use of public places for recreational and sporting activities.
2. **Non-market impacts of European wasp outweigh market impacts** Without a formal management programme, the impacts on biodiversity, use of public places for recreation and human health were estimated to be more than one-and-a-half times the market impacts over a 50-year period.
3. **Benefits of biological control outweigh the costs** Four plausible biological control scenarios, based on different values of agent growth, mortality and effectiveness, were chosen for analysis. In all cases the introduction of the control agent reduced damages. The reduction in damages ranged from $14.1m to $95m. Benefit-cost ratios ranged from 2.7 and 12.5 for the scenarios analysed.

### 1.2 Recommendations

While the economics of controlling European wasp using biological control looks promising because of the sizable benefits of control, we cannot recommend proceeding with *S. v. vesparum* as the biocontrol agent based on current knowledge about the performance of this agent. Rather we recommend the department:

1. **Develop closer contact with NZ experts.** New Zealand has recently imported *S. v. vesparum* collected from the United Kingdom, thought to be the likely origin of New Zealand’s European wasps, and thus more likely to increase the parasitoid’s effectiveness in controlling the wasp. The Australian biocontrol effort could leverage off this research. We therefore recommend that closer contact with the New Zealand biological control team be established in order to benefit from the NZ research and improve our understanding of likely agent performance in Australia.
2. **Undertake case studies** Case studies should be undertaken to gain additional insights into efficient biocontrol release strategies and selection of locations for release. A first step would be to analyse the ACT’s eWasp dataset. This dataset is a resource that contains detailed information about nest locations over time, method of detection and stinging events.
3. **Investigate other biological control agents** Based on research currently underway in New Zealand, there appear to be several biological control agents that might provide additional control of the European wasp:

- the mite Pneumolaelaps niutirani
- A mermithid nematode in the genus *Steinernema*.
- *Volucella* hoverflies.

## 2. Introduction

Many social insects – wasps, termites, bees and ants – are highly successful invaders, becoming major pests when they establish outside their native ranges (Moller, 1996). Of the invasive wasps, two *Vespula* species are notable for their ecological, economic and human impacts: the European wasp^1^, (*Vespula germanica*); and the common wasp (*Vespula vulgaris*). The European wasp is the most widespread of the two species (Lester and Beggs, 2019). It is native to Europe, Northern Africa, and temperate Asia, and introduced into North America, Chile, Argentina, Iceland, Ascension Island, South Africa, Australia and New Zealand. It can tolerate or adapt to a wide range of habitats and climates (de Villiers et al., 2017) and has significant negative impacts on communities, industry and the environment in regions where it has been introduced.

Both invasive *Vespula* species are established in Australia and New Zealand. In New Zealand, the total quantifiable annual impact of *Vespula* wasps on primary industries, human health, traffic accidents^2^, and local governments was estimated at NZD133 million, including an option value for apiculture development of NZD58 million (MacIntyre and Hellstrom, 2015). The European wasp is the most widespread of the two species in Australia. The damage caused by European wasp in the south-eastern part of Australia, where the wasp established almost 60 years ago has not been calculated, although is likely to be substantial given the lack of any sustained and widespread control strategies during this time.

Established pests such as the European wasp are often overlooked as candidates for management programmes (eradication and/or containment) because the use of traditional control techniques over very large areas becomes uneconomic. Use of biocontrol agents in this context, where the goal is permanent establishment of the agent and control of the pest (rather than eradication) is usually the only economically feasible option, however the processes around approvals for screening, testing and release of biocontrol agents can take significant amounts of time and resources, especially if screening and testing of potential agents is required.

The European wasp could be a candidate for a renewed management programme in south-eastern Australia given the availability of a biocontrol agent, released in parts of Victoria during 1991 and 1992, but this would depend on the benefits to industry, community and the environment from a reduction in European wasp abundance. This project will explore the benefits and costs of European wasp management using a biocontrol agent, and importantly, will include valuation of the social and environmental impacts of the pest. The common wasp will not be considered in this analysis due to its restricted range in Australia, although it may have significant impacts in cooler regions such as Tasmania (B. Brown, personal communication, December 8, 2020).

### 2.1 Objectives

The main objective of this project is to understand the costs and benefits of European wasp management using classical biological control, and as a result, whether additional funds should be made available to control this widespread pest through such a programme.

The model developed in this study will be used to answer the following questions:

1. What are the costs and benefits of European wasp control, and how do these vary with pest density and spread?
2. How much will it cost to reduce the population of European wasps to a particular density, with a particular likelihood of success?
3. What are the time frames within which the European wasp could be reduced to a particular density for a given budget?
4. How effective would the biological control agent need to be in order to reduce European wasp to a particular density over a particular period of time?
5. How should populations of biological control agents be managed in order to maintain low population densities of European wasps?
6. What magnitude are the measurable non-market impacts, and does their inclusion change the business case?

### 2.2 Methodology

This project explored the benefits and costs of European wasp management using a decision analysis model, where ‘management’ comprised *classical* biological control – the introduction of a specialised natural enemy, from the region of origin of an invasive pest (Avtzis et al., 2020). The biocontrol agent of interest in the current context is *Spechophaga vesparum vesparum,* a parasitoid wasp which attacks European wasp nests and feeds on the developing larvae and pupae. The use of *inundative* biological control, where the goal of agent release is to overwhelm the pest rather than to permanently establish the agent, is not considered in this study.

While benefit-cost analysis (BCA) is the standard method of evaluating the cost-effectiveness of response options in the management of pest incursions (van Wilgen et al., 2004; Paine et al., 2015), decision analysis, based on economic principles and with grounding in ecology and other sciences, has made significant contributions to the management of invasive species in recent years (Epanchin-Niell and Hastings, 2010). Adopting a modelling approach will allow more flexibility in the analysis of management scenarios compared to BCA – the latter typically only considers the ‘do nothing’ in addition to one or two management scenarios.

## 3. The European wasp in Australia

### 3.1 Biology

The European wasp is a ‘social’ insect, and possesses characteristics of eusociality that allow it to succeed as an invasive species (Moller, 1996; Beggs et al., 2011): there is an overlap of generations; reproduction is restricted to a few individuals; and there is cooperative brood care (Wilson and Hölldobler, 2005). European wasp colonies are highly eusocial, consisting of reproductive females (gynes and queens), reproductive males (drones) and sterile females (workers).

The normal colony lifecycle is annual, with nests being founded by a single reproductive queen in the spring. The reproductive cycle progresses as follows (Figure 1): the queen emerges in spring to forage and prepare new nests of about 20 cells in size; workers emerge 4-6 weeks after egg-lay and assume foraging, nest protection and nest building duties, thus freeing up the queen for egg laying; the nest grows over summer and the new queens are produced in autumn; these queens mate with drones in autumn then fly off to hibernate in sheltered areas during winter (Widmer et al., 1995).

**Figure 1.**
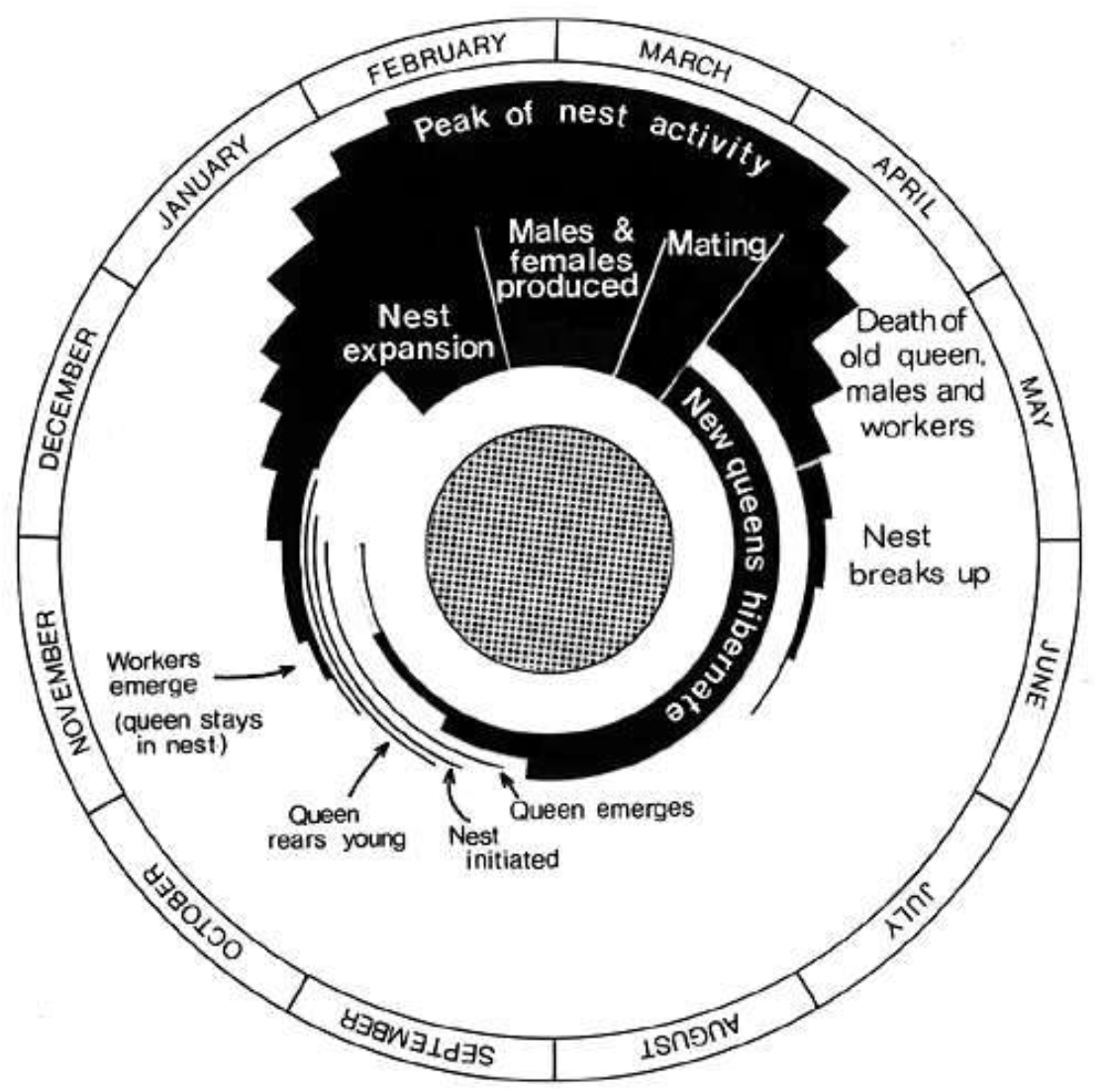
European wasp lifecycle. Source: Landcare Research New Zealand (2012).

Nests consist of a series of hexagonal cells used for rearing young and are arranged in a roughly circular pattern, with layers of cells forming ‘combs’ (Kasper, 2004). Nests are constructed from chewed wood fibre (pulp), and are usually located underground, although may be found in hollow trees and man-made structures. Nest sizes, and thus wasp numbers, increase rapidly through summer and autumn. Analysis of nests in South Australia over three seasons shows wasp numbers growing from 2000 in December to more than 17,000 per nest during May (Table 1) (Kasper, 2004).

**Table 1.**
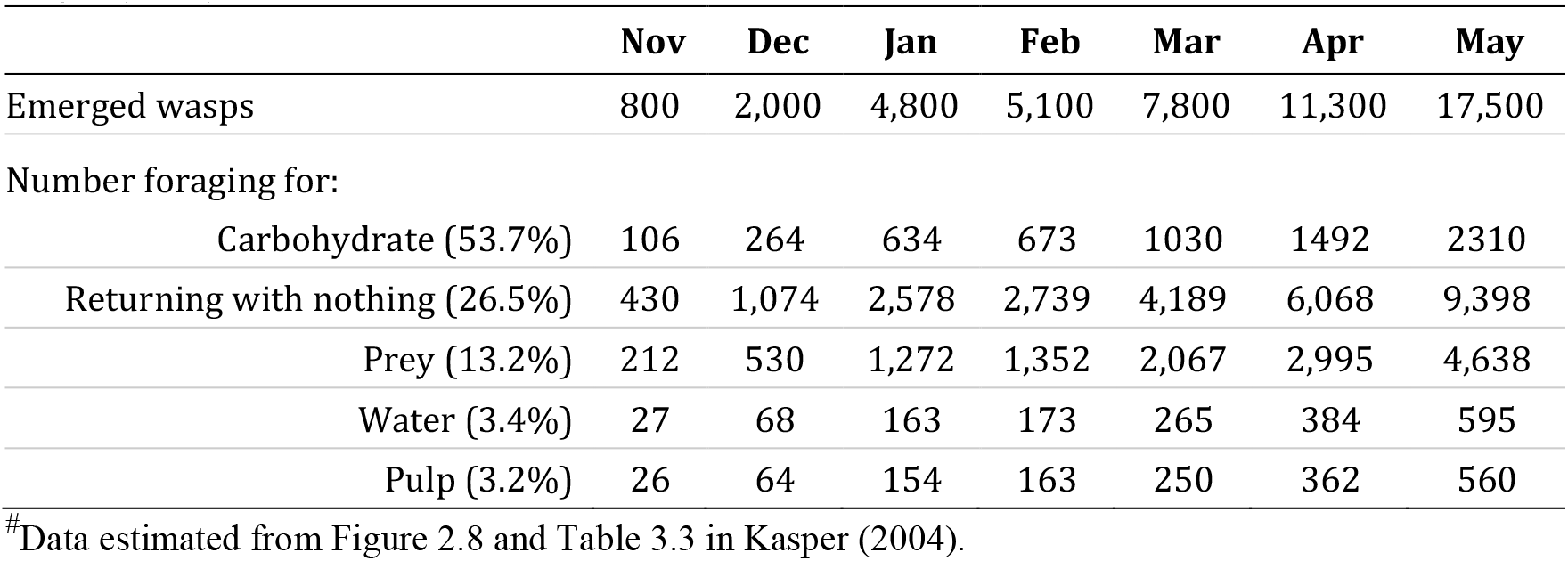
Numbers of wasps emerged from annual *V. germanica* nests and foraging behaviour. Source: <u>Kasper (2004).</u>

In their native range, European wasp colonies naturally die off in winter. This is not the case in locations experiencing mild winters, including parts of Australia, where nest construction can continue throughout the year, and over-wintered nests are common (Spradbery and Maywald, 1992; Widmer et al., 1995; Kasper, 2004). Overwintering is a modification of the usual annual lifecycle, where polygyny (multi-queening) occurs ‒ more than one, and often many hundreds, of productive queens share the same nest. As a result, overwintered nests can reach very large sizes in summer, producing thousands of individuals.

#### 3.1.1 Foraging behaviour

Foraging distance from nests is not well documented, but is reportedly up to 500 m (Goodman, 2014). European wasps forage for animal prey (protein) to feed developing larvae, carbohydrates (concentrated sugar) to satisfy their own energy requirements and those of their workers, wood pulp for nest building, and water for thermoregulation inside the nest (Harris, 1991; Richter, 2000). Sampling of large numbers of foraging wasps in South Australia over several months revealed most wasps forage for carbohydrates (54%), followed by wasps returning with nothing (26.5%) and wasps returning with prey (13.2%) (Table 1) (Kasper, 2004).

Wasps collect carbohydrates opportunistically from a range of sources, including nectar from flowers, plant sap and sweet liquids from fruit (Evans and Eberhard, 1970 cited in Richter, 2000). Foraging for prey is also opportunistic and from a wide range of insects including bees (Harris, 1991). European wasps can thus be problematic for beekeepers, particularly in autumn when their need for sugar reaches a peak. In addition to robbing hives of honey, European wasps remove dead or dying bees found in front of the hive entrance, prey on live bees and enter hives to remove brood (Goodman, 2014).

### 3.2 History of spread

While the first detection of individual European wasps in Australia was in a timber consignment in Sydney in 1954 (Chadwick and Nikitin, 1969), the first nests were discovered in 1959 in Hobart, and subsequently in Western Australia (1977), Victoria (1977), New South Wales (1978), and South Australia (1978) (Spradbery and Maywald, 1992). The pest is now well-established in Tasmania, Victoria, South Australia, New South Wales, and the Australian Capital Territory. In those jurisdictions, eradication programmes were typically initiated upon first discovery of the pest, but subsequently abandoned as the pest became widespread (Crosland, 1991). The public is now referred to private pest controllers or may attempt nest destruction themselves, to remove nests on private land. In Western Australia the European wasp is the subject of an ongoing eradication campaign. The state-government funded campaign commenced in the 1980s and involves surveillance and the destruction of all nests found within the state.

### 3.3 Impacts

European wasps have significant negative impacts on horticulture, apiculture, tourism and outdoor social activities, animal health and biodiversity in regions where they have been introduced. This analysis considers the damage caused by the wasp to human activity, the environment and the economy via six specific types of damage: i) use of public places for sporting and recreational use (*public areas damage*); ii) impacts on households (*household damage*); iii) impacts on the natural environment (*nature conservation damage*); iv) impacts on beekeeping (*honey production damage*); v) impacts on pollination (*pollination damage*); and vi) impacts on ripened soft fruit (*fruit damage*).

#### 3.3.1 Impacts on human activity

##### Public area damage

European wasps are a major nuisance because of their synanthropic behaviour ‒ they aggressively forage for human food (sugar and protein), disrupting outdoor dining and recreational activities, sporting activities and use of public places in general.

##### Household damage

European wasps are more aggressive than bees and will attack when their nests are disturbed and they are capable of inflicting multiple painful stings on humans and pets. Unlike bees, European wasps do not die after stinging. Wasp stings may cause allergic reactions, and a sustained attack from a large swarm can result in life-threatening envenomation (McGaln et al., 2000). While relatively rare, deaths have occurred as a result of European wasp stings, and there are many records of stings requiring medical attention and even hospitalisation (Widmer et al., 1995; Levick et al., 1997). People have to avoid outdoor areas where wasps might live.

#### 3.3.2 Environmental Impacts

##### Nature conservation damage

European wasps have a broad, omnivorous and opportunistic diet that includes honeydew, nectar, insect prey, vertebrates and carrion (Lester and Beggs, 2019; Spencer et al., 2020). As a result, they may have disruptive impacts on a range of ecosystem process, including reducing the numbers of some arthropods (Sackmann et al., 2000; Kasper, 2004). In New Zealand the pest is known to reduce faunal diversity as a result of direct competition for food, particularly honeydew (Elliott et al., 2010), predation on other insects (Harris, 1991) and even nestling birds (Moller, 1990). There may also be adverse impacts on local flora due to wasp predation on insects responsible for pollination and other forms of nutrient transfer (Fordham, 1991). There is little information about environmental impacts of European wasps in Australia, although local reductions in arthropods have been reported in Tasmania (Bashford, 2001; Potter-Craven et al., 2018).

European wasps may have positive impacts. For example, *Vespula* wasp species are effective pollinators and their foraging pressure may replace that of native species that have been lost from the system (see studies cited in Lester and Beggs 2018). Since none of these impacts have been measured under Australian conditions we simply note that they may exist, and do not explore them further in the analysis.

#### 3.3.3 Impacts on primary industries

European wasps cause significant losses to beekeeping, and to many horticultural industries through their impact on pollination, honey production and damage to ripened soft fruits, all of which are considered in this study. Other impacts on workers and livestock through stinging are not considered.

##### Honey production

European wasps cause significant losses to apiculture by attacking bees and bee hives ‒ the wasps kill honey bees and their larvae for protein, rob hives of honey and spread bee diseases (Clapperton et al., 1989; Widmer et al., 1995). While strong bee colonies are able to repel attacks, significant losses may still occur from sustained attack (Goodman, 2014). Defending hives against wasps reduces bee foraging time.

Bee keepers must devote resources to managing hives in order to prevent destruction and raiding by European wasps. In 2019, beekeepers in New Zealand lost between 0.6% and 1.6% of their bee colonies to European wasps (Stahlmann-Brown et al., 2020) and in a bad wasp year they could expect to lose up to 5% of their hives (Spradbery 1986 cited in Widmer et al. 1995). It is thought similar damage could be expected in Australia (Crosland, 1991).

Losses to beekeeping in NZ from the direct effect of wasps on hives was estimated at $8.8m per annum (MacIntyre and Hellstrom, 2015). This value captures wasp control management costs; the cost of hives lost to wasps; and production losses from bees focusing on defence rather than food collection. Cook (2019) estimated the damage to apiculture in Western Australia, where European wasps are the subject of an eradication programme, would reach more than $1.1 million per year if wasps were left unmanaged. Managed hives in Western Australia typically produce at least 1,600 tonnes of honey per year worth around $5 million.

##### Pollination damage

European wasps affect pollinators through competition for resources and predation. In depleting honey-bee colonies, wasps impact on pollinator-reliant crops. Crops experiencing reduced yield because of reduced pollination, and their level of pollinator reliance are given in Table 2.

**Table 2.** Reliance of various crops on pollination by bees

| Crop | Pollinator reliance (%) |
| --- | --- |
| cotton | 20<sup>a</sup> |
| canola | 16<sup>c</sup> |
| sunflower | 30<sup>a</sup> |
| peanut | 10<sup>a</sup> |
| pulse_oth | 10<sup>a</sup> |
| citrus_oth | 20<sup>a</sup> |
| mandarin | 30<sup>a</sup> |
| orange | 30<sup>a</sup> |
| avocado | 100<sup>a</sup> |
| mango | 90<sup>a</sup> |
| orchard_oth | 20<sup>a</sup> |
| apple | 90<sup>a</sup> |
| pear | 50<sup>a</sup> |
| apricot | 70<sup>a</sup> |
| cherry | 90<sup>a</sup> |
| nectarine | 60<sup>a</sup> |
| peach | 60<sup>a</sup> |
| stone_oth | 70<sup>a</sup> |
| fruit_other | 10<sup>a e</sup> |
| almond | 100<sup>a</sup> |
| macadamia | 90<sup>a</sup> |
| beans | 10<sup>a</sup> |
| lettuce_tot | 10<sup>a</sup> |
| melon | 100<sup>a</sup> |
| onion | 100 |
| pumpkin | 90<sup>a</sup> |

<sup>a</sup>Gill; <sup>b</sup>McGregor (1976) cited in Cunningham et al. (2002); <sup>c</sup>Manning and Boland (2000);
<sup>d</sup>Delaplane and Mayer (2000) as cited in Cunningham et al. (2000); <sup>e</sup>de Oliveira (1997) as cited in Cunningham et al. (2000).

##### Fruit damage

Social wasps opportunistically exploit any available source of concentrated sugar, including the sweet liquids from fruits, and use these as an energy source for adult wasps and developing young (Evans and Eberhard, 1970 cited in Richter, 2000). European wasps are known to cause yield losses by hollowing out fruit (Goodall & Smith as cited in Cook, 2019) and damage wine grapes by introducing diseases (Lester and Beggs, 2019). In Australia, wine grapes and strawberries have reportedly been damaged by European wasps, with yield losses of 10-25% being reported (Table 3).

**Table 3.** Reported damage by wasps to fruit crops

| Crop | Location | Yield loss (%) | Source; Notes |
| --- | --- | --- | --- |
| Grape | Tasmania; | 25 | Bashford (2001); several vineyards, 1999/2000 |
| Grape | Victoria | 10-15 | Thomas (1993) cited in Bashford (2001). |
| Strawberry | Tasmania | 20 | Bashford (2001); one grower affected 2000 |

### 3.4 Biological control using Spechophaga vesparum vesparum

In 1980, the European wasp parasitoid *Spechophaga vesparum vesparum* was collected from Europe and imported into New Zealand for testing and subsequent mass rearing (Donovan and Read, 1987). Field releases of the parasitoid, as yellow overwintering cocoons, began in 1987. Cocoons were placed in release boxes, specially designed to give maximum protection from rodents, insects and shelter from the weather. Release boxes were initially stocked with around 100 cocoons and placed at 39 sites across New Zealand (Read et al., 1990; Moller et al., 1991). Boxes were replenished over subsequent seasons and by 1990 more than 108,000 yellow parasitoid cocoons had been released (Read et al., 1990). Subsequent monitoring at release sites indicated the parasitoid was having difficulty becoming established (Moller et al., 1991) although establishment was confirmed at two sites (Beggs et al., 2008). The poor performance of *S. v. vesparum* in New Zealand is thought to be caused by a ‘genetic bottleneck’ because all releases were essentially derived from a single female parasitoid [Ward, 2014; Beggs et al. 2008]. Thus, chances of successful control could be improved by sourcing different genetic strains of parasitoids from different populations in Europe.

In conjunction with the New Zealand biological control programme, the parasitoid was imported into Australia, tested for host-specificity (Field and Darby, 1991) and mass reared. Between 1989 and 1993 over 120,000 cocoons were released across parts of south-eastern Australia (Lefoe et al., 2001), however monitoring of release sites was not well-funded and as a result no evidence that the parasitoid established at any of the release sites was collected (R. Kwong, personal communication, July 3, 2020). Although more testing is required to draw a final conclusion, it is reasonable to conclude that poor performance of the agent in New Zealand is mirrored in Australia, and that sourcing different genetic strains of parasitoids would see an improvement in agent performance in Australia.

#### 3.4.1 Sphecophaga vesparum vesparum lifecycle

While the process of predation by *S. v. vesparum* on European wasp is relatively straightforward – it attacks European wasp nests and feeds on the developing larvae and pupae – its lifecycle is complex (Beggs et al., 1996; Harris and Rose, 1999). During spring, winged male and female *S. v. vesparum* emerge from cocoons in the remains of old wasp nests, one to four seasons after their cocoons were formed. Females enter new wasp nests and lay eggs onto larvae or pupae, with the parasitoid larvae feeding on the host and subsequently forming one of three types of cocoons: yellow, weak-walled yellow and white. Yellow cocoons are thick-walled that will remain dormant in the nest, producing winged adults up to four years later. Weak-walled yellow cocoons produce winged adults capable of flight within two weeks. Weak-walled white cocoons produce short-winged females within two weeks and these continue to lay eggs within the parental nest and multiple generations follow during the same season (Harris and Rose, 1999).

## 4. Modelling the spread and management of European wasp

The conceptual model in Figure 2 represents the decision analysis model that will be used to analyse the costs and benefits of European wasp control in south-eastern Australia. Western Australia is excluded as the state still has an eradication program. Central to the model is the population dynamics of the wasp and the biocontrol agent, *S. v. vesparum*. Together, these ultimately determine the likelihood that management of the pest – in this case through biocontrol release and maintenance – will succeed. The population dynamics model is the focus of the current chapter.

**Figure 2.**
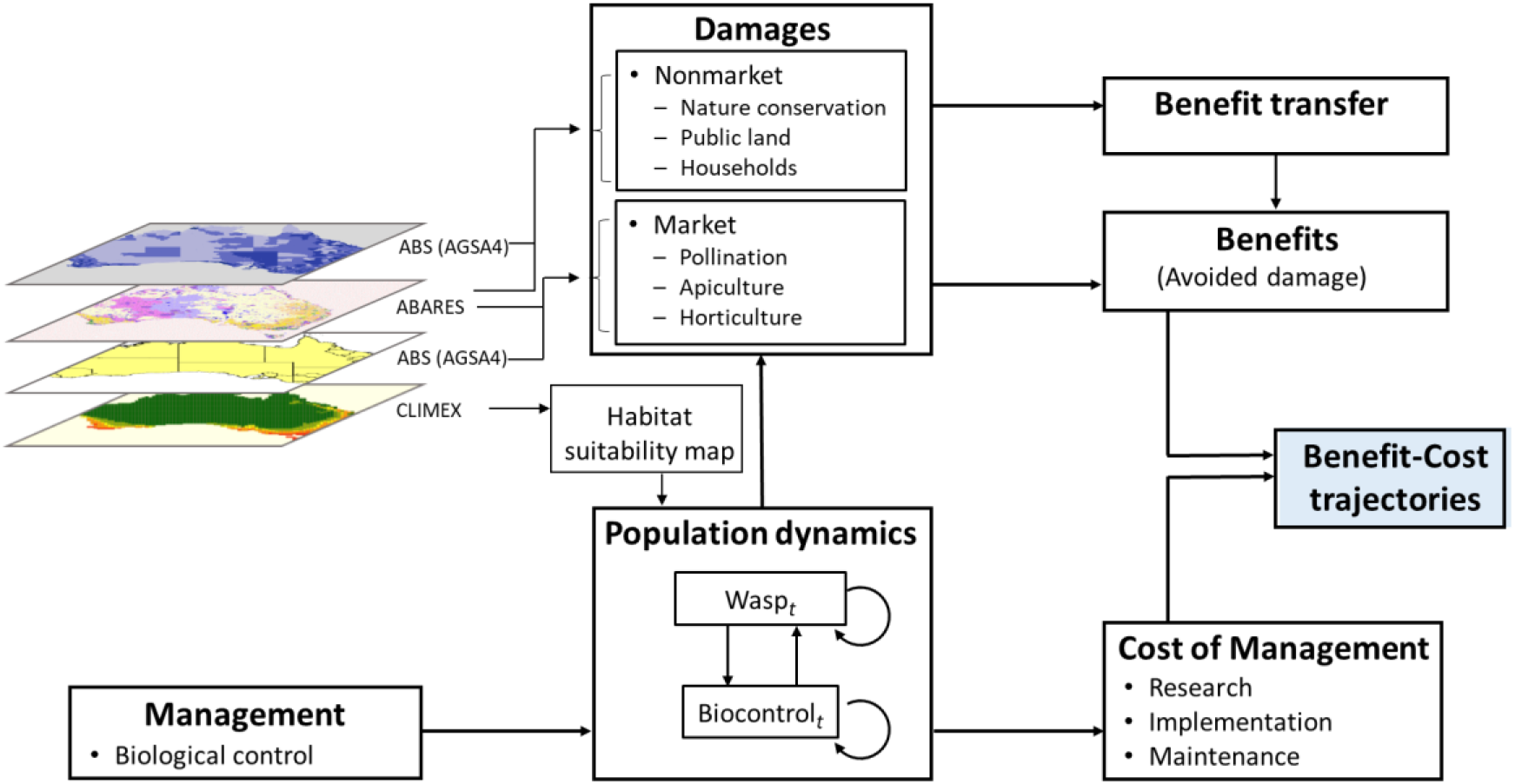
Conceptual model

On the output side of the model (top of Figure 2), the wasp impacts on pollination, honey production and horticulture (market damages); nature conservation, use of public places for recreation and sporting activities, and households (non-market damages). The total amount of damage occurring depends on variables that are related to location, such as land use, population density, presence of nature reserves, and habitat suitability (see Chapter 5). Damages are converted to dollar values through market and non-market valuation. Calculation of market damages is described in Section 5.3. Benefit-transfer was used to calculate non-market impacts (see 5.4 and Appendix A).

The situation where no management occurs shows the worst-case-scenario for wasp impacts over time. This is the baseline scenario and shows the value of damages against which all other management outcomes are compared. Four feasible biological control scenarios were tested, and these are described in Section 6.1 Controlling the spread and density of wasps through a biological control programme is aimed at reducing damages. The benefits of biological control are therefore the damages that are avoided because the pest is being managed. To determine these, the net benefits of the biological control programme are compared to the impacts from not having a biological control programme. These are described in Chapter 6 and Appendix C.

### 4.1 Population dynamics

Studies from Australia (Kasper, 2004) and New Zealand (Barlow et al., 2002) have modelled the population dynamics of European wasp using a Ricker equation (Ricker, 1975), incorporating weather effects and density dependence. We modify the model of Barlow et al. (1996) to represent the spread of wasps and parasitoids and their interaction. The model consists of Ricker growth equations and Cauchy dispersal equations for both the wasp (*W*) and the biocontrol agent (*B*). The interaction between the two species is introduced through the growth parameters as explained below.

#### 4.1.1 Growth

Based on the life cycles presented in Chapter 3, the wasp population is expressed as number of nests per ha in spring (*W_s_*) and autumn (*W_a_*), whereas the biocontrol population is expressed as number of adults per ha in spring (*B_s_*) and autumn (*B_a_*). The value of *B_a_* is a proxy for the number of cocoons that overwinter in wasp nests.

The growth model is:

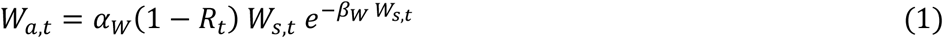

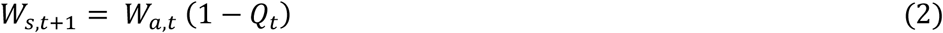

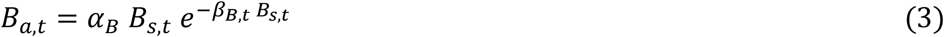

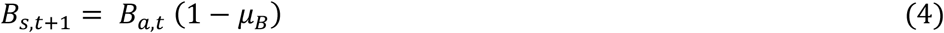

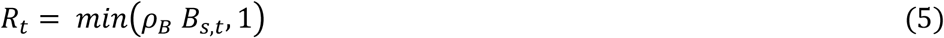

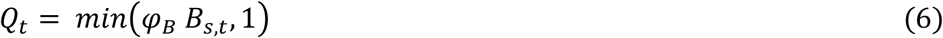

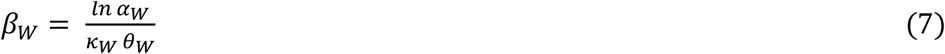

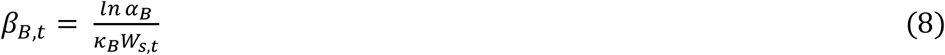

Variables are defined in Table 4 and parameter values are presented in Table 5. The time step (*t*) is one year, with each year starting in spring, when wasp queens emerge to start building up new nests.

**Table 4.** Variables of population growth model. In the numerical model all variables are column vectors with 522 rows representing cells in the Climex map that have w asp habitat suitability > 0.

| Variable | Description |
| --- | --- |
| $W_{s,t}$ | Number of wasp nests per ha in spring of year $t$ |
| $W_{a,t}$ | Number of wasp nests per ha in autumn of year $t$ |
| $B_{s,t}$ | Number of adult parasitoids (biocontrol agent) per ha in spring of year $t$ |
| $B_{a,t}$ | Number of adult parasitoids (biocontrol agent) per ha in autumn of year $t$ |
| $R_t$ | Reduction in wasp growth rate caused by biocontrol agent in year $t$ |
| $Q_t$ | Winter mortality rate of wasp nests caused by biocontrol agent in year $t$ |
| $K_t$ | Number of wasp nests destroyed by humans per ha in year $t$ |

**Table 5.** Population model parameters

| Parameter | Value | Description | Source |
| --- | --- | --- | --- |
| $\alpha_W$ | 3.237<br>(3.202 - 3.325) | Wasp growth parameter in Ricker equation | (a) |
| $\beta_W$ | eq (7) | Wasp growth parameter in Ricker equation | (c) |
| $\kappa_W$ | 0.912<br>(0.891 - 0.967) | Carrying capacity of wasps (nests/ha) when $\theta = 1$ | (a) |
| $\theta_W$ | 0 - 1 | Wasp habitat suitability of site | (d) |
| $\gamma_W$ | 4.925<br>(4.903 - 4.969) | Dispersal parameter for wasps | (a) |
| $\delta_W$ | 0.341<br>(0.039 - 0.597) | Wasp detection probability parameter | (a) |
| $\alpha_B$ | 2.0 | Parasitoid growth parameter in Ricker equation | (b) |
| $\beta_B$ | eq (8) | Parasitoid growth parameter in Ricker equation | (c) |
| $\kappa_B$ | 235 | Carrying capacity of parasitoid (cocoons per wasp nest) | (b) |
| $\mu_B$ | 0.85 | Parasitoid winter mortality | (b) |
| $\rho_B$ | 0.004 | Reduction in wasp growth rate per adult parasitoid | (b) (e) |
| $\varphi_B$ | 0.005 | Winter mortality of wasp nests per adult parasitoid | (b) (e) |
| $\gamma_B$ | 3.0 | Dispersal parameter for parasitoid | (e) |

(a) GA calibration (see section 4.3); (b) Barlow et al. (1996); (c) algebraic estimation; (d) based on Climex map ( $\theta = 1$ when eco-climatic index is at its maximum); (e) plausible, subject to sensitivity analysis.

Equations (5) and (6) represent reductions in, respectively, the spring growth rate and winter survival rate of wasp nests caused by the biocontrol agent. Equation (7) relates wasp growth rate to the maximum carrying capacity (*κ*_*W*_) and the habitat suitability of the site (*θ*_*W*_). Finally, equation (8) relates the biocontrol growth rate to the maximum numbers of cocoons per wasp nest (*κ*_*B*_) and the number of wasp nests per ha in spring (*W_s,t_*), which together represent the carrying capacity of the site.

We assume that all nests detected by humans are destroyed, at a cost of $250 per nest (J. Bariesheff, personal communication, June 29, 2020). The probability that a nest will be detected is given by:

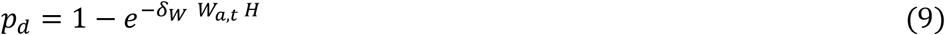

where H is the number of households per ha and *δ_W_* is a detectability parameter related to the probability of encounter between humans and wasps given their respective local density. The number of nests destroyed is:

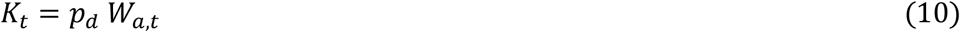

and after destruction the number of nests is updated to:

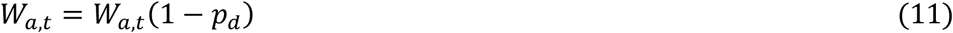

All the variables above and in Table 4 are expressed as column vectors of dimension 522 × 1, where the rows are cells in the Climex map that have wasp habitat suitability > 0. A *site* refers to one of these rows and it has a direct correspondence to a cell in the Climex map in Figure 11 (Chapter 5). The results of solving the model (1)-(8) for a planning horizon of *T* years are presented as matrices of dimensions 522 × *T*, where rows are Climex cells and columns are time periods. There is one such result matrix for each vector. Such that:

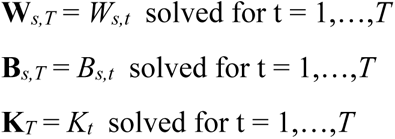

This matrix representation of results is useful for two reasons: (1) it allows analysis of patterns in space as well as in time, and (2) it fits directly with the impact model, where commodities and land uses are presented as vectors of 522 rows corresponding to Climex cells.

#### 4.1.2 Dispersal

Each spring, queens emerge from hibernation and start building new nests and multiplying their colonies. Dispersal to new sites can occur early in the season as queens find nesting sites, and later in the season as new queens establish new nests. For simplicity we assume that dispersal occurs in spring. The probability that a queen emerging in site *i* will move to site *j* and establish a nest there is given by:

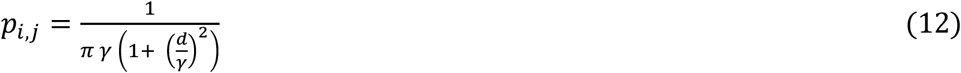

Where *γ* is a dispersal parameter and *d* is the distance between points *i* and *j*. In the numerical model distance is expressed as a matrix **D** of dimensions 522 ×522, representing the distance between each Climex cell and every other cell on the map in km. This allows dispersal probability to be estimated for all sites at once, yielding the matrix **P** of the same dimensions as **D**. The expected number of nests after dispersal for the given state *N_t_*, where *N_t_* represents the number of wasp nests or parasitoids at any time (i.e. *W_s,t_* and *B_s,t_*), is calculated as:

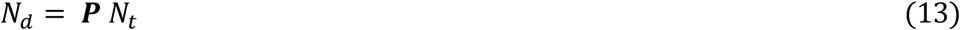

*N_d_* is a 522 ×1 vector of expected numbers after dispersal, however, our purpose is to represent stochastic dispersal so this equation is not used in the model. The actual dispersal for each run of the model is selected by sampling from a uniform distribution using the Matlab (Mathworks, 2020) random number generator for each column in **P** given *N_t_*.

### 4.2 Biocontrol Analysis

The success of the biocontrol program hinges on two factors: (1) the feasibility that the parasitoid will establish and spread; and (2) the effectiveness of the parasitoid in suppressing growth and spread of wasp nests. Factor (1) is related to three biocontrol parameters: growth rate (*α_*B*_*), winter mortality (*μ*_*B*_) and spread rate (*γ_*B*_*) of the parasitoid. Factor (2) is related to two biocontrol parameters: the reduction in wasp growth rate (*ρ*_*B*_) and the winter mortality of wasp nests (*φ*_*B*_) caused per adult parasitoid. In this section we analyse how these parameters affect the feasibility of using the parasitoid *S. v. vesparum* as a biocontrol agent for *V. germanica*.

#### 4.2.1 Growth and spread rates

The relative rate of increase for a population, R = *N_t_*_+1_/*N_t_*, must be > 1 in order for the population to grow. Barlow et al. (1996) present evidence of low *R* values for *S. v. vesparum* in New Zealand and they explain possible causes based on the underlying parameters in their model. **The bottom line is that if *R* < 1 for the biocontrol**, there is no need to undertake further evaluation of its potential benefits, as **the organism will be unable to establish a viable population**. In this case the main question is whether there are mechanisms for increasing the value of *R*, and at what cost.

Base parameter values for the biocontrol were presented in Table 5. Figure 3 (a) shows that, for the base value of *α_B_* = 2, winter mortality must be < 0.5 in order for the parasitoid population to grow, given the presence of wasp nests. This is lower than the mortality of 0.85 estimated by Barlow et al. (1996) and so an important question is whether this rate may be lower in Australia given the dryer conditions compared to NZ.

**Figure 3.**
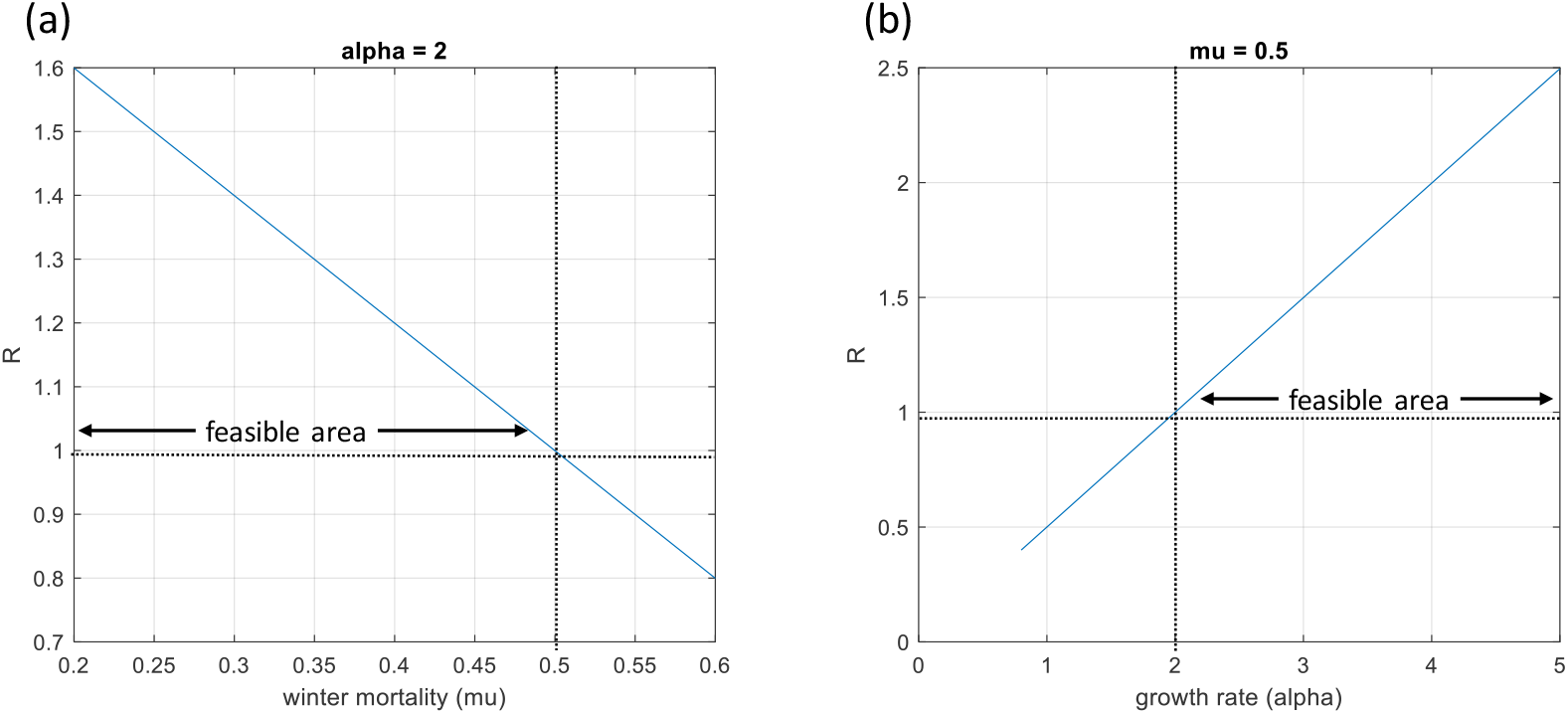
Effect of mortality and growth parameters on feasibility of establishment of the biocontrol agent. The feasible area indicates parameter values that result in R > 1.

The rate of spread of the biocontrol agent influences the probability that the parasitoid will establish across the landscape where wasp nests are present. The rate of spread is determined by *γ*_*B*_ (Figure 4). Barlow et al. (1998) estimated a velocity of spread of 1.15 to 1.6 km per year.

**Figure 4.**
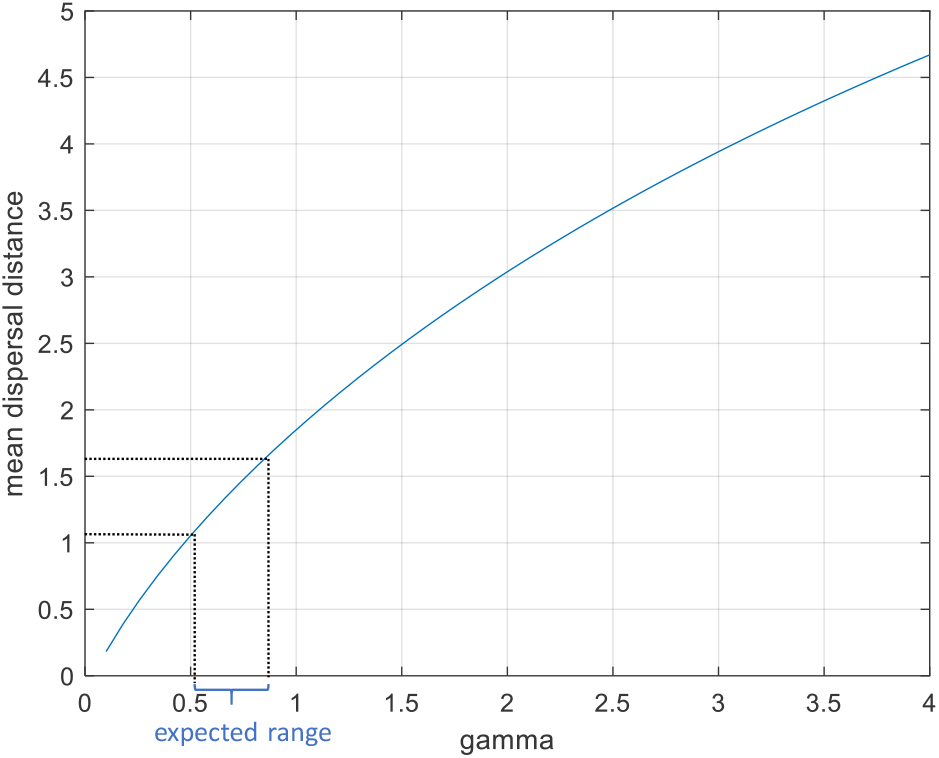
Expected range of *γ_B_* values based on spread velocity estimated for *S. vesparum* in New Zealand.

#### 4.2.1 Biocontrol effectiveness

The effectiveness of the biocontrol hinges on its ability to suppress growth and spread of the wasp infestation. In the model this occurs through two parameters: the reduction in wasp growth rate (*ρ*_*B*_) and the winter mortality of wasp nests (*φ*_*B*_) caused per adult parasitoid. Evidence to estimate values for these parameters comes from Barlow et al. (1996) for New Zealand, adjusted based on simulations to understand the effects of different combinations of parameter values for the Australian case (see section 4.3 for details of model calibration). These are uncertain parameters that have not been measured for Australian conditions – while some Australian release sites of *S. v. vesparum* were monitored (G. Lefoe, pers. comm.) there was no evidence in subsequent years that the parasitoid established at any of those sites (Lefoe et al., 2001). However, using the model we can explore combinations of parameter values that would make a biological control program feasible in the sense of *classical* biological control, where the biocontrol agent becomes established and maintains the population of the pest at a low level (Hajek et al., 2016).

Figure 5 shows the time cycles of the biocontrol and the wasp from simulations at different combinations of parameter values. The top panel (Figures 5(a) and 5(b)) represents relatively high growth rate (*α_B_*) and winter mortality rate (*μ_B_*) for the parasitoid, within the feasible areas illustrated in Figure 3, whereas the bottom panel (Figures 5(c) and 5(d)) represents relatively low*α_B_* and *μ_B_*. In all cases there is a cyclical behaviour in the population densities of both species as expected. However, at high growth and mortality rates, both the frequency and amplitude of the cycle are higher (top panel in Figure 5) than at low growth and mortality rates (bottom panel). Comparing the left and right panels, it is interesting that relative high effectiveness (*ρ*_B_ = *φ_B_* = 0.04) increases the fluctuations in both populations when *α_B_* and *μ_B_* are high (Figure 5(b)), resulting in a higher average wasp density. In contrast, when *α_B_* and *μ_B_* are low, high effectiveness produces the best results in the set (Figure 5(d)) in the long term, with the lowest wasp population density as the cycle converges to a stable equilibrium between wasps and parasitoids. However, the slow rate of decrease in the wasp population between years 0 and 30 for this scenario means that substantial damage from wasps may accumulate before reaching this low equilibrium.

**Figure 5.**
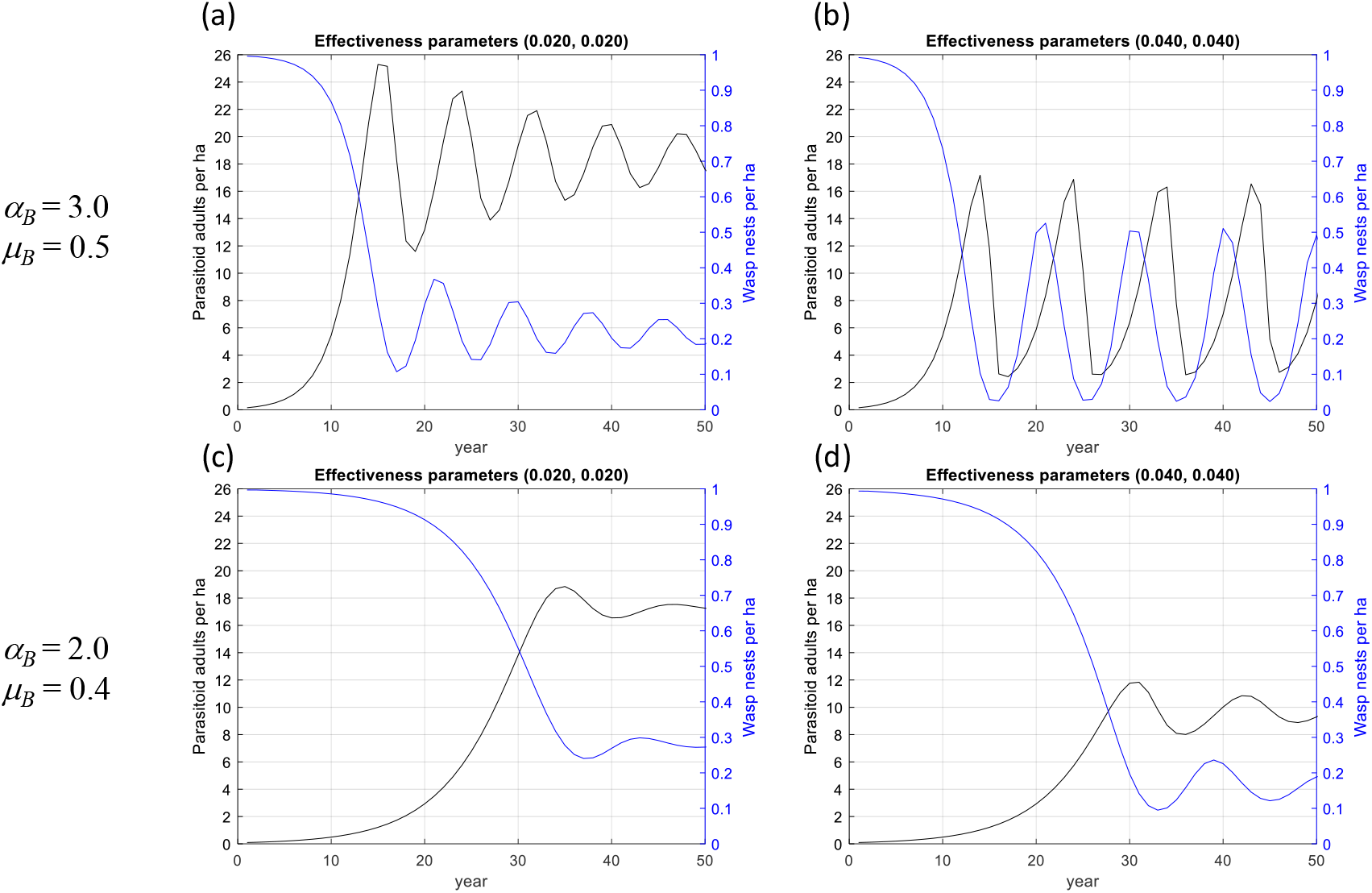
Time trajectories of adult *S. v. vesparum* parasitoids per ha and *V. germanica* wasp nests per ha, model solutions at alternative combinations of growth (α_B_, μ_B_) and effectiveness (*ρ*_B_, *φ_B_*) parameters.

The cases we are interested in are those that tend towards some sort of equilibrium between the two species, with wasp density maintained at a relatively low level. These are the cases for which classical biological control can succeed in the long term. This approach allows us to screen alternatives before considering the costs and benefits of the program. **At this stage, scientists may be able to assess whether it is likely that the biocontrol agent will reach the required levels of growth, mortality and pest suppression to make the program feasible**, perhaps through additional research targeted at measuring the most uncertain parameters. This applies to any biocontrol agent beyond this case study.

Figures 6 and 7 present the cycles above as phase diagrams. Plotting biocontrol agent density against wasp density for the full trajectory provides useful information on the dynamics of the biocontrol-pest interaction. In all cases selected for analysis, the system starts on the top left of the diagram (high wasp density - low parasitoid density) and moves down and to the right towards some sort of dynamic equilibrium. In some cases, the system tends towards a steady state (Figure 6(a)) and in others towards a stable cycle (Figures 6(b) to 6(d)).

**Figure 6.**
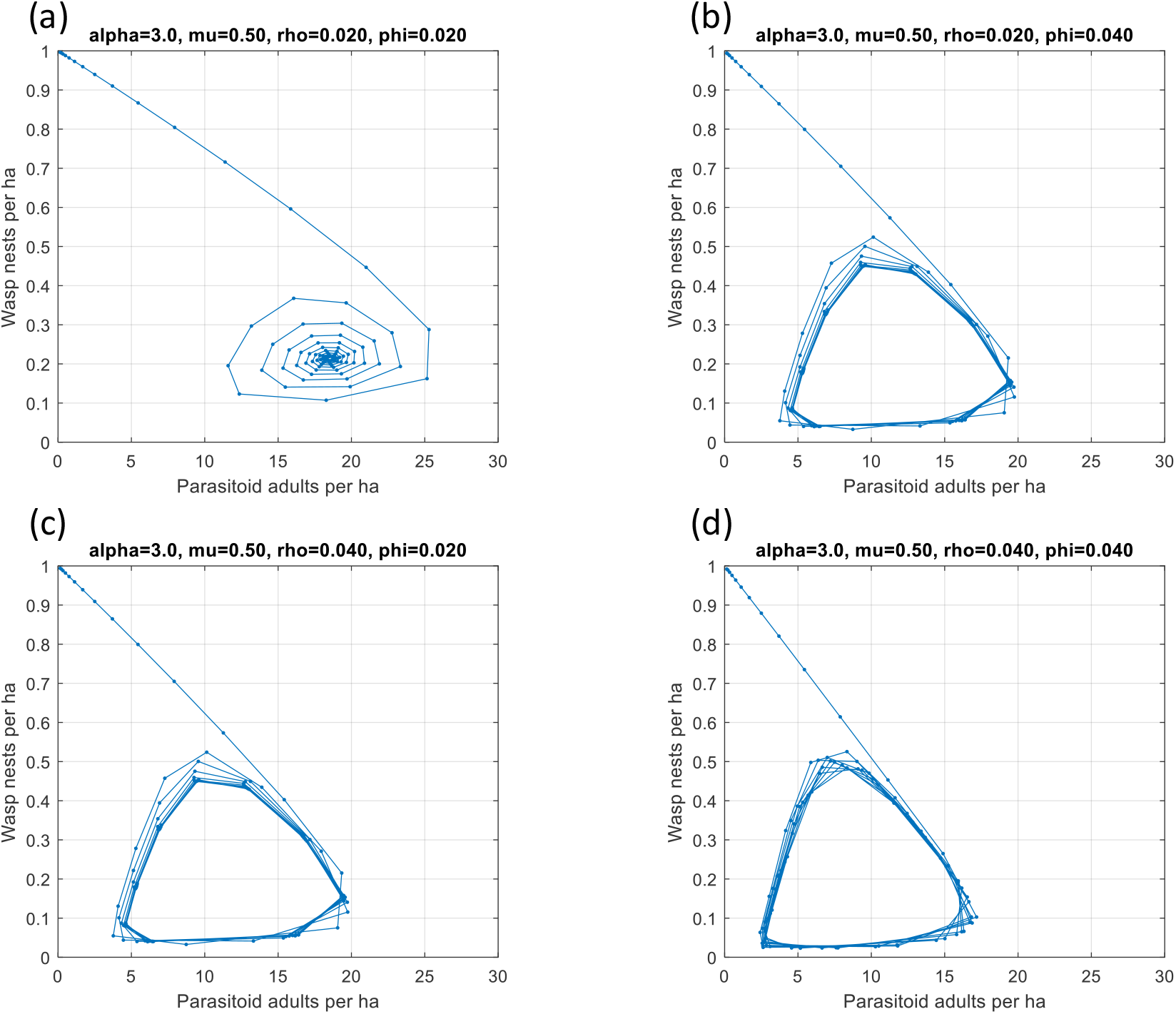
Phase diagrams of adult *S. v. vesparum* parasitoids per ha against *V. germanica* wasp nests per ha, model solutions at alternative combinations of growth effectiveness (*ρ_B_*, *φ_B_*) parameters, with *α_B_* = 0.3 *μ_B_* = 0.5.

**Figure 7.**
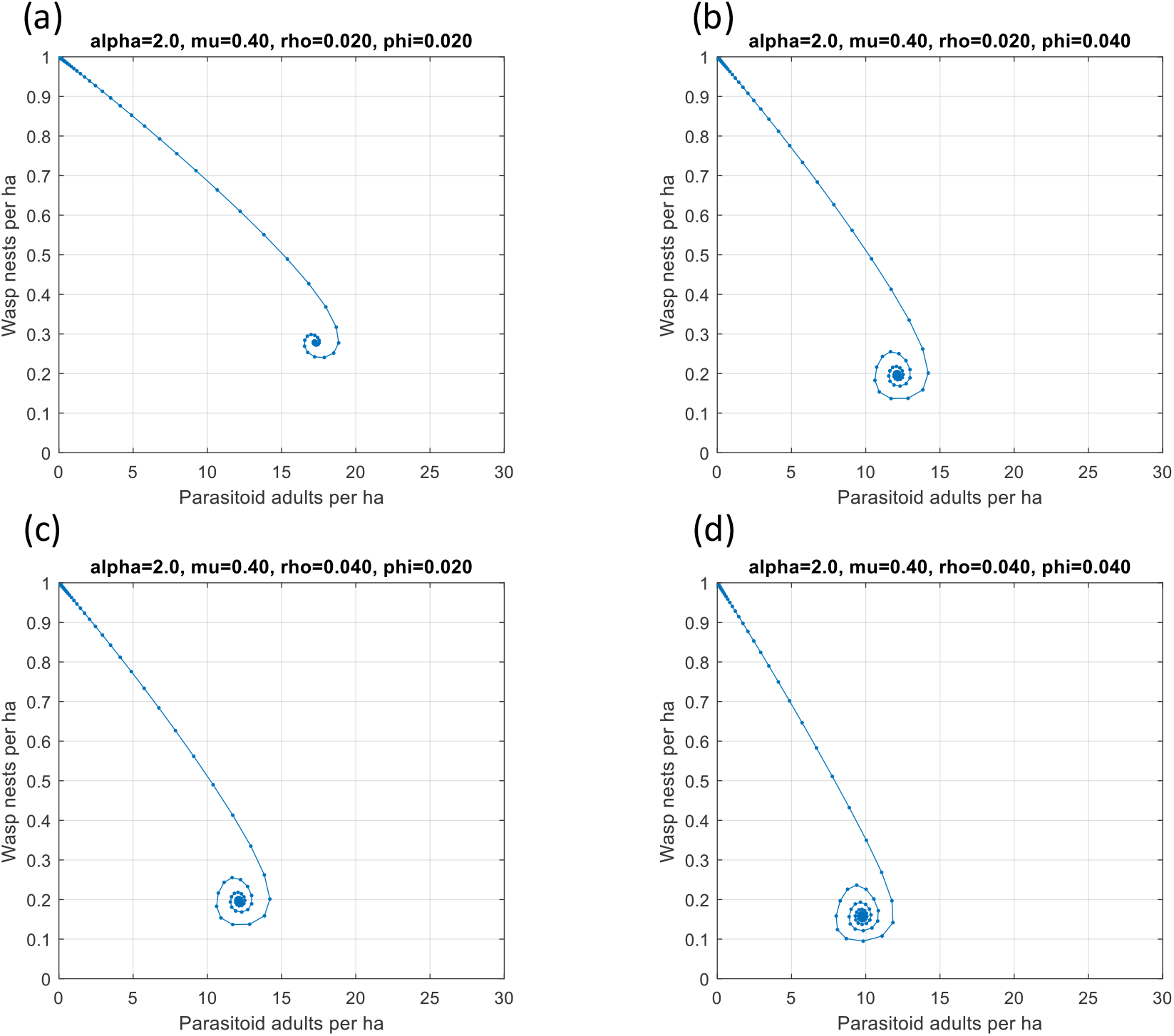
Phase diagrams of adult *S. v. vesparum* parasitoids per ha against *V. germanica* wasp nests per ha, model solutions at alternative combinations of growth effectiveness (*ρ_B_*, *φ_B_*) parameters, with *α_B_* = 0.2 *μ_B_* = 0.4.

The phase diagrams show that a combination of low growth and mortality rates leads to more stable states in the long term, with wasp densities between 0.15 and 0.3 nests per ha (Figure 7) depending on the effectiveness parameters, as compared to the case of high growth and mortality rates (Figure 6) where the tendency is towards broad cycles ranging between 0.05 and 0.5 nests per ha. In such case wasps would cause high damages in some years but not in others. The trade-off is that those damages are reduced to a steady state 10-20 years earlier than under the low growth-low mortality scenario, from Figure 5.

Based on evidence from New Zealand, expected winter mortality (*μ_B_*) is ∼0.85 (Barlow et al. 1996). There would be a problem if a similar value applies in Australia, since the maximum mortality for success of biocontrol is 0.5 (Figure 3(a)), given other parameter values. Based on the literature (Toft et al., 1999), leading causes of winter mortality of *S. vesparum* cocoons in NZ is by flooding of underground nests and predation by rodents. Perhaps cocoon mortality may not be as high in areas of Australia where flooding during winter is uncommon and this would work in favour of likelihood of success. A more detailed review of the scientific evidence in the context of feasibility of introducing the biocontrol was out of the scope of this project but may be worthwhile to consider before making a final decision.

### 4.3 Running simulations and estimating parameter values

Parameters for the population dynamics of European wasps are available for New Zealand (Plunkett et al., 1989; Barlow et al., 1996; Beggs et al., 2008) and England (Archer, 1985) but not for Australia. Some parameters reported in the literature provide a starting point for calibrating our model, but differences between countries need to be considered. Notably, the maximum number of wasp nests per ha (*κ*_*W*_) reported for beech forests in NZ (∼12 nests per ha) is too high for Australian conditions (Crosland, 1991; Kasper, 2004; Tennant et al., 2011). There is also uncertainty regarding the number of years it takes for the population to reach saturation in the absence of control, which is determined by *α*_*W*_ for a given *κ*_*W*_. Other important sources of uncertainty are the dispersal parameter (*γ*_*W*_) and the detection probability parameter (*δ*_*W*_). These four parameters were estimated through simulation using a genetic algorithm (GA) combined with what we know about the history of the invasion in Australia.

Simulations using the model (1) to (13) were undertaken as described in Box 1. To estimate likely values of uncertain parameters a likelihood function was created based on known facts and the simulation model. The known facts are:

1. The first European wasp nests were discovered in Tasmania in 1959 (Spradbery and Maywald, 1992).
2. By 1982 the wasp had spread to several areas in the southern part of mainland Australia (Crosland, 1991) and eradication was attempted in several locations.
3. Wasp detection records in the Atlas of Living Australia (ALA) were reported as early as 1960^3^, with increasing reports since 2010 (Figure 8).

**Figure 8.**
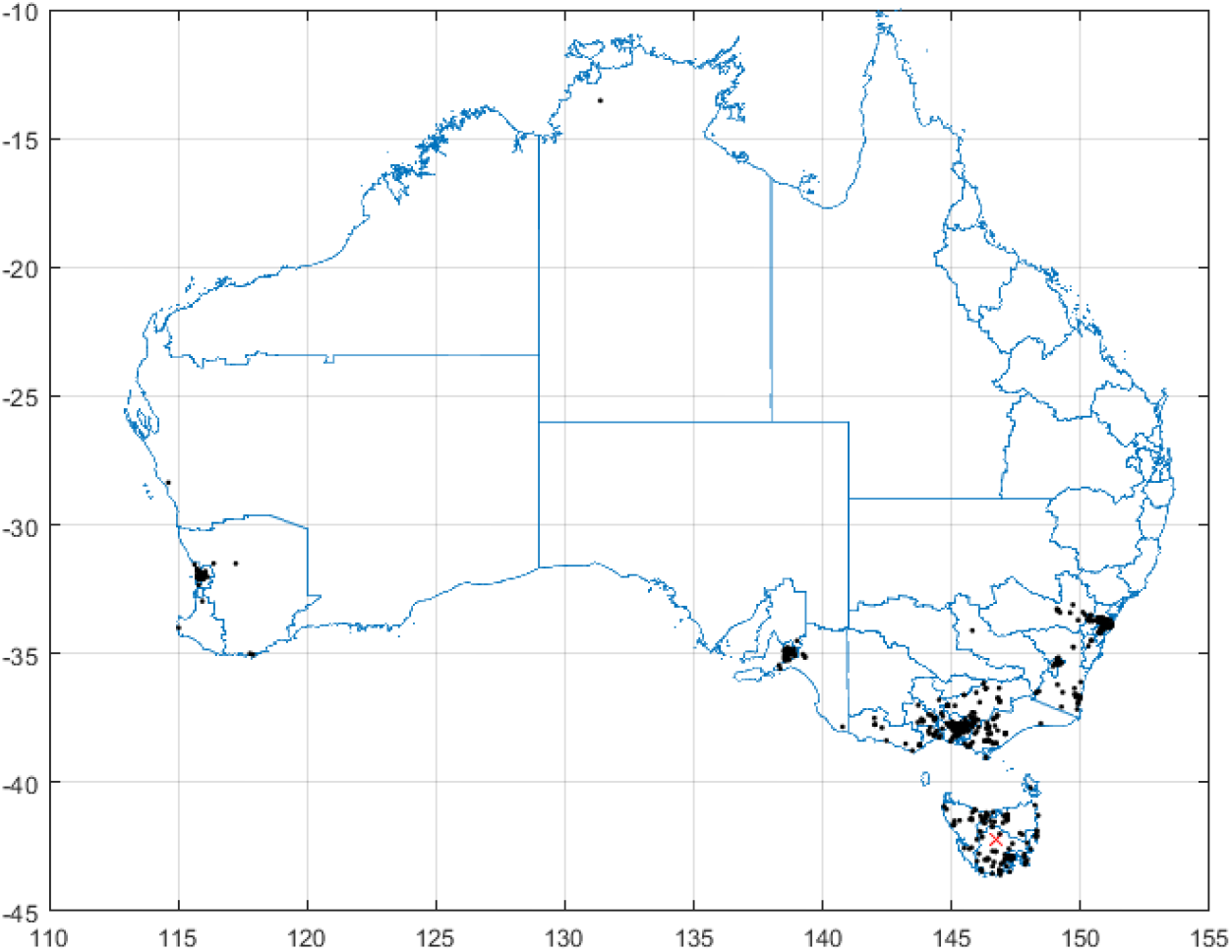
European wasp reports downloaded from Atlas of Living Australia (black points) and initial invasion in Tasmania (red x).

Box 1. Simulation algorithm.

1. Set the desired number of stochastic iterations *nr* and planning horizon (*T*) to the desired values and initialise random number generator to default value.
2. Set the initial number of parasitoids, *B_s_*_,0_ = *B*_0_, where *B*_0_ is an initial vector (522×1) of parasitoids placed within cells on the map where wasp nests are present. This is a management variable to be varied across scenarios.
3. Set the initial number of spring wasp nests for the given scenario, *W_s_*_,0_ = *W*_0_, where *W*_0_ is an initial vector (522×1) of wasp nests per ha for all cells on the map. This is the initial state of the invasion.
4. For each stochastic run *n* = 1, ..,*n_r_* select a parameter set at random from given distributions and go to step 5.
5. For each year *t* = 1, ..,*T* estimate wasp dispersal for *N_s,t_* using random sampling based on (12) and (13) and update *N_s,t_* map.
6. Run the growth model (1) to (11) and save results in column *t* of **W***_s,T_*, **B***_s,T_* and **K***_T_*.
7. Return to step 5 until *t* = *T*.
8. Return to step 4 until *n* = *n_r_*. Save results from each stochastic run for later analysis and application.

The likelihood function is

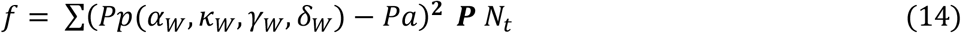

Where *P_p_* is the predicted probability of wasp nest presence and *P_a_* is the actual presence based on the ALA reports (Figure 8). The *Pp* vector is obtained using the simulation algorithm in Box 1 with additional steps involving the GA:

a. In Step 1 (Box 1) set *n_r_* = 100 and *T* = 60.
b. Define a parameter vector *z* = [*α_W_*, *κ_W_*, *γ_W_*, *δ_W_*] and corresponding bounds *z*^+^= [4.0, 1.2, 7.0, 0.6] and *z*^-^= [2.0, 0.7, 3.0, 0.0], these values are set within plausible limits based on the literature.
c. Pass *z, z*^+^ and *z*^-^ to the GA, which adjusts wasp population parameters prior to running simulations and uses the likelihood function to estimate the fitness of each solution.
d. In Step 2 (Box 1) set the biocontrol agent *B_s_*_,0_ = 0 for all cells, representing the case with no biocontrol program.
e. In Step 3 (Box 1) set the initial wasp invasion *W_s_*_,0_ = 0 for all cells except for *W_s_*_,0_ = 0.01 for cell number 20 which is located in Tasmania (Figure 8). This represents the initial invasion detected in 1959.
f. Run simulation Steps 4 to 8 (Box 1) for each stochastic iteration and save the final result vector *W_s,T_* as column *n* in matrix **W***_N_*.
g. Estimate 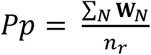 and apply likelihood function *f*. Use *f* as the fitness function in the GA.

Steps a to g are incorporated into the GA (Hester and Cacho, 2012), which starts with a population of 50 individuals with values of *z* selected randomly within the bounds. Each realisation of *z* represents a ‘gene’ consisting of 4 ‘chromosomes’ to be modified in a process analogous to natural selection to find the value of *z* that minimises *f*. When the GA converges, it produces not only the optimal parameter set *z*\*, but also the final population of the best 50 genes, based on fitness. These optimal parameter sets are saved in matrix **Z**\* and are used to sample randomly in further simulations. This approach allows us to account for uncertainty in parameter values while ensuring that the model explains the known data. Results of GA iterations are shown in Figure 9 showing that the solution tends to converge within 50 iterations. mean best

**Figure 9.**
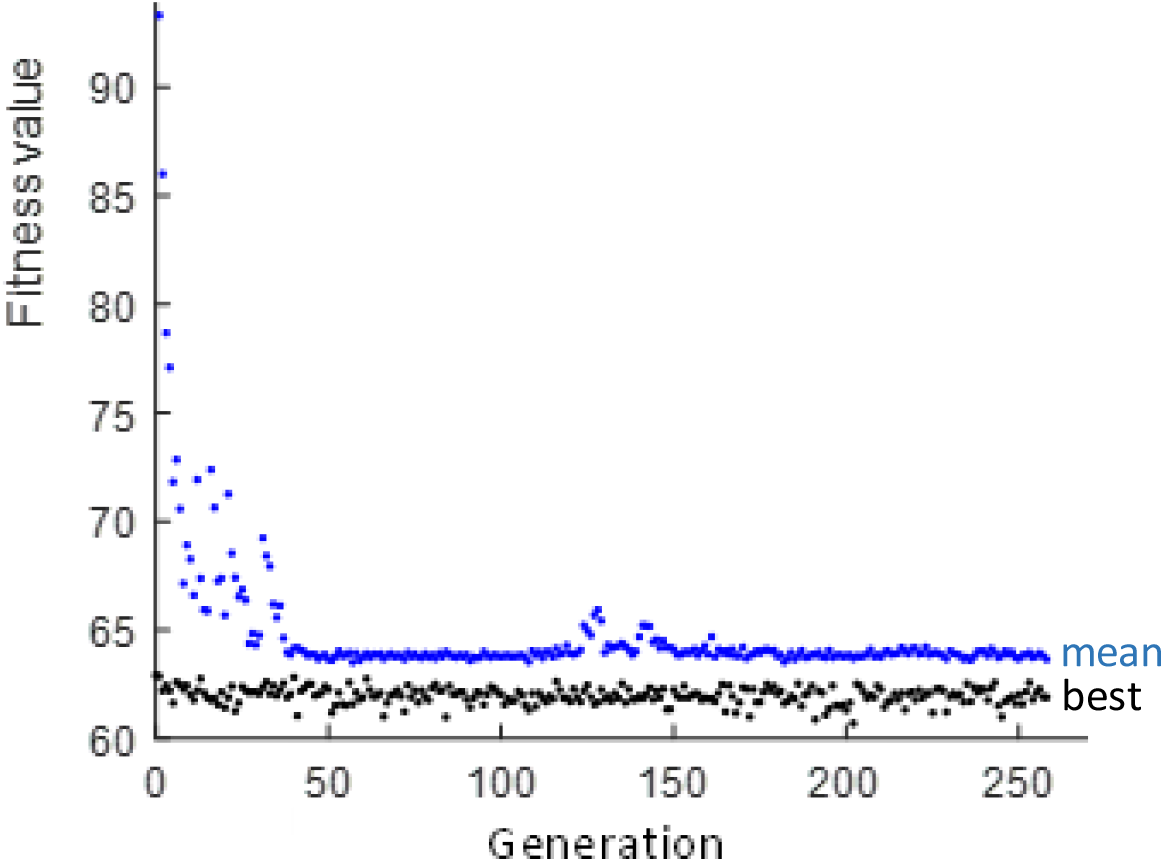
Genetic algorithm results showing convergence of the population within 50 generations.

The analysis was limited to cells in the Climex map that were suitable for wasp establishment in the Eastern states. Western Australia was excluded as the state still has an eradication program. The probability map for the relevant area along with the ALA reports of wasp presence is presented in Figure 10.

**Figure 10.**
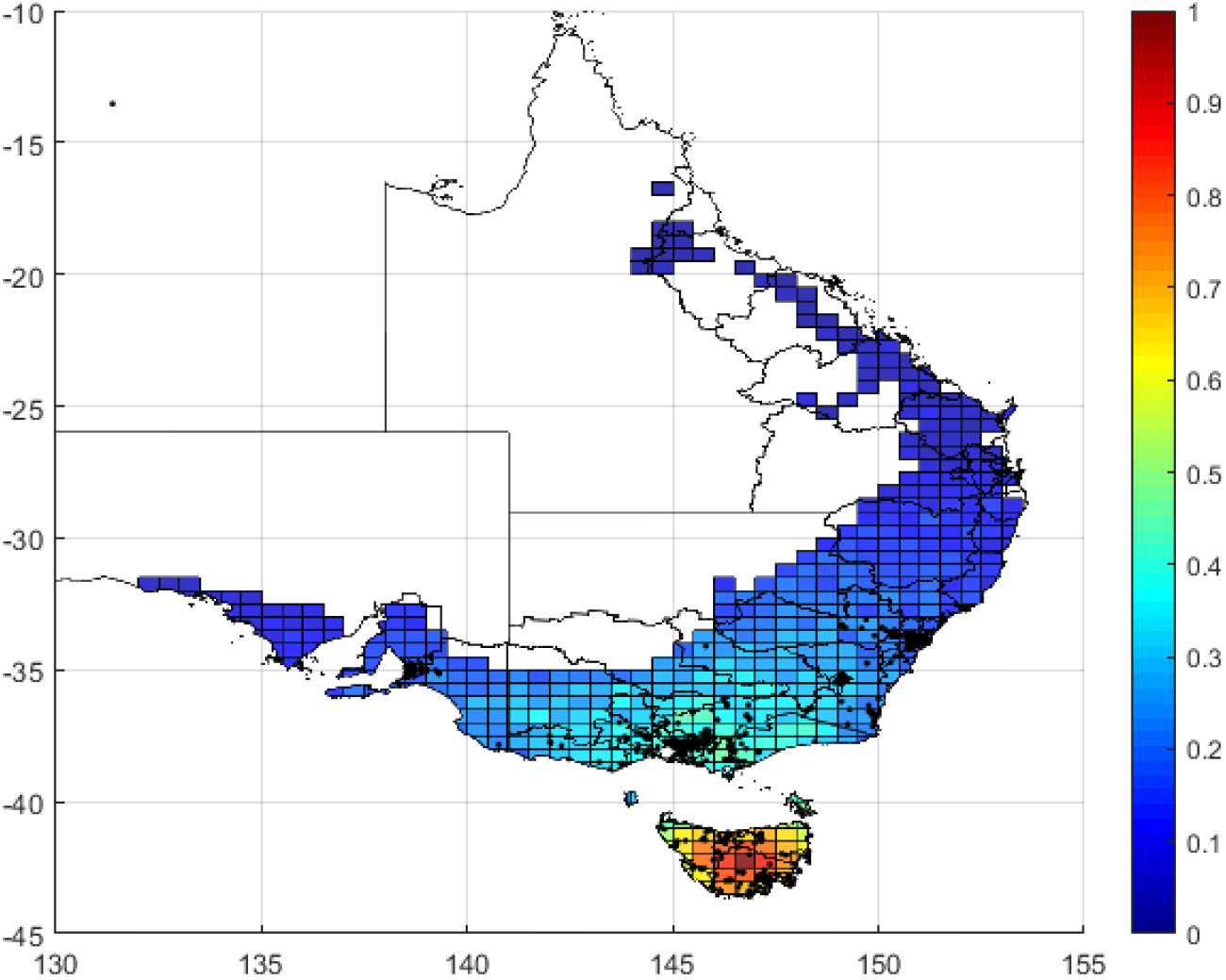
Probability map of wasp presence generated by the model (coloured cells) and ALA reports of wasp presence (black dots) overlaid on the relevant SA4 regions from ABS.

## 5. Data and estimation of damages

In order to understand the market and non-market impact of wasps, information was required on: the value of agricultural production; land use; habitat suitability, wasp detections over time, income and inflation.

### 5.1 Data sources

Figure 2 in Chapter 4 illustrated the input datasets as a set of maps overlaid on top of each other. Four spatial datasets were used as inputs:

1. Map of *V. germanica* habitat suitability in Australia based on Climex simulations from de Villiers et al. (2017) at a resolution of 0.5 degrees.
2. National map of land uses and agricultural commodities from ABARES^4^. The number of agricultural commodities in this dataset is 27.
3. Maps of agricultural commodities (2017-18) at SA4 level from the ABS for value of production, area planted and yields^5 6^.
4. Number of Households and income at SA1 level from the ABS 2016 Census Community Package^7^.

These maps are shown in Figures 11 to 14. The unit of analysis is determined by the habitat suitability map (the Climex map) and a series of matrix manipulations were used to convert the original source data to a final dataset where each row represents a Climex cell and columns represent commodities and land uses. The dimensions of the matrices involved in these calculations are:

*n_w_* = 522 number of Climex cells with wasp habitat suitability > 0. In the Eastern states of Australia.
*n_s_* = 76 number of SA4 regions that overlap with Climex cells in the *n_w_* set.
*n_c_* = 65 number of commodities in ABS datasets.
*n_u_* = 27 number of agricultural land uses in the ABARES dataset.

**Figure 11.**
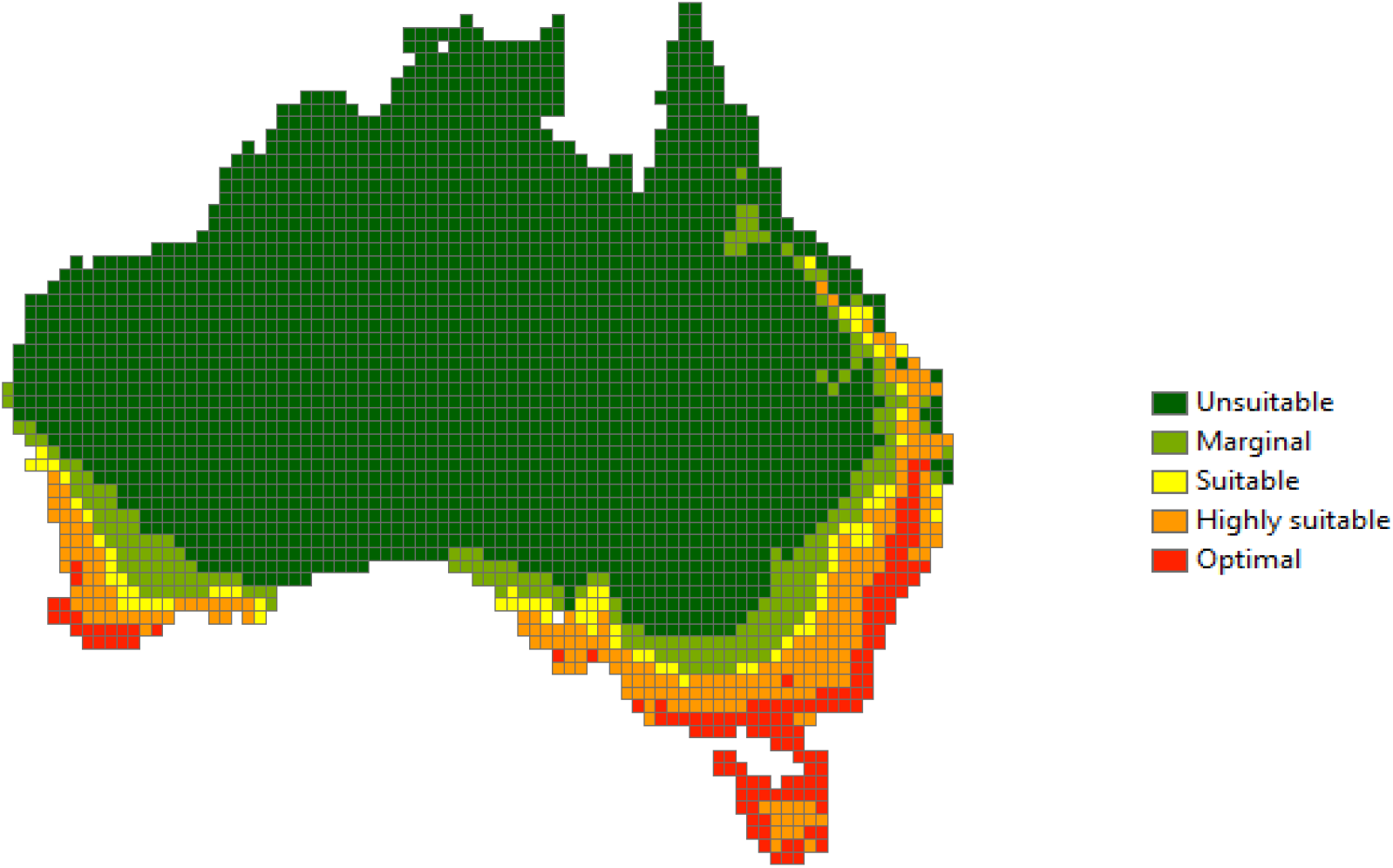
Habitat suitability of European wasp in Australia based on CLIMEX simulations

**Figure 12.**
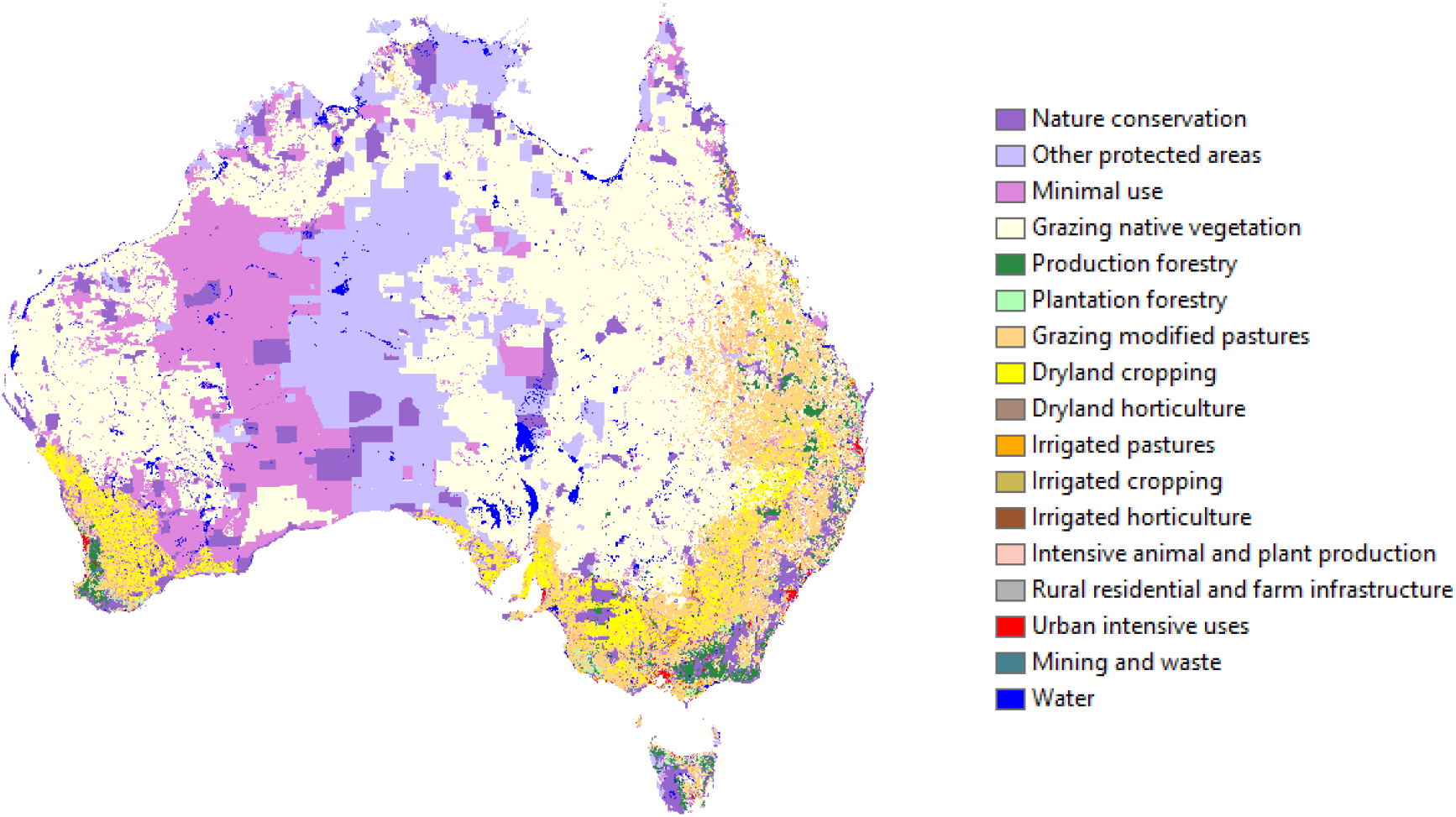
National map of land uses and agricultural commodities

**Figure 13.**
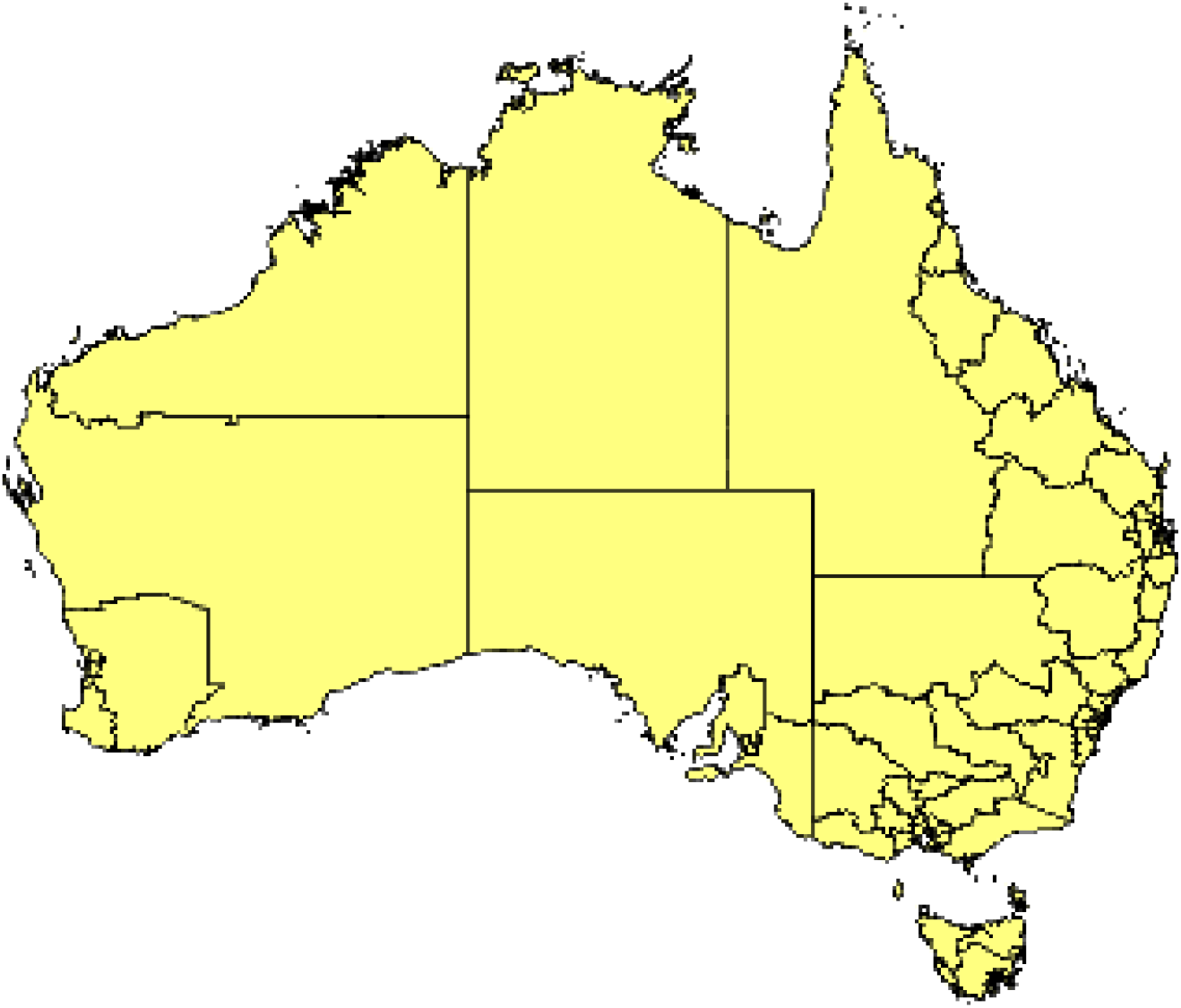
Map of SA4 regions.

**Figure 14.**
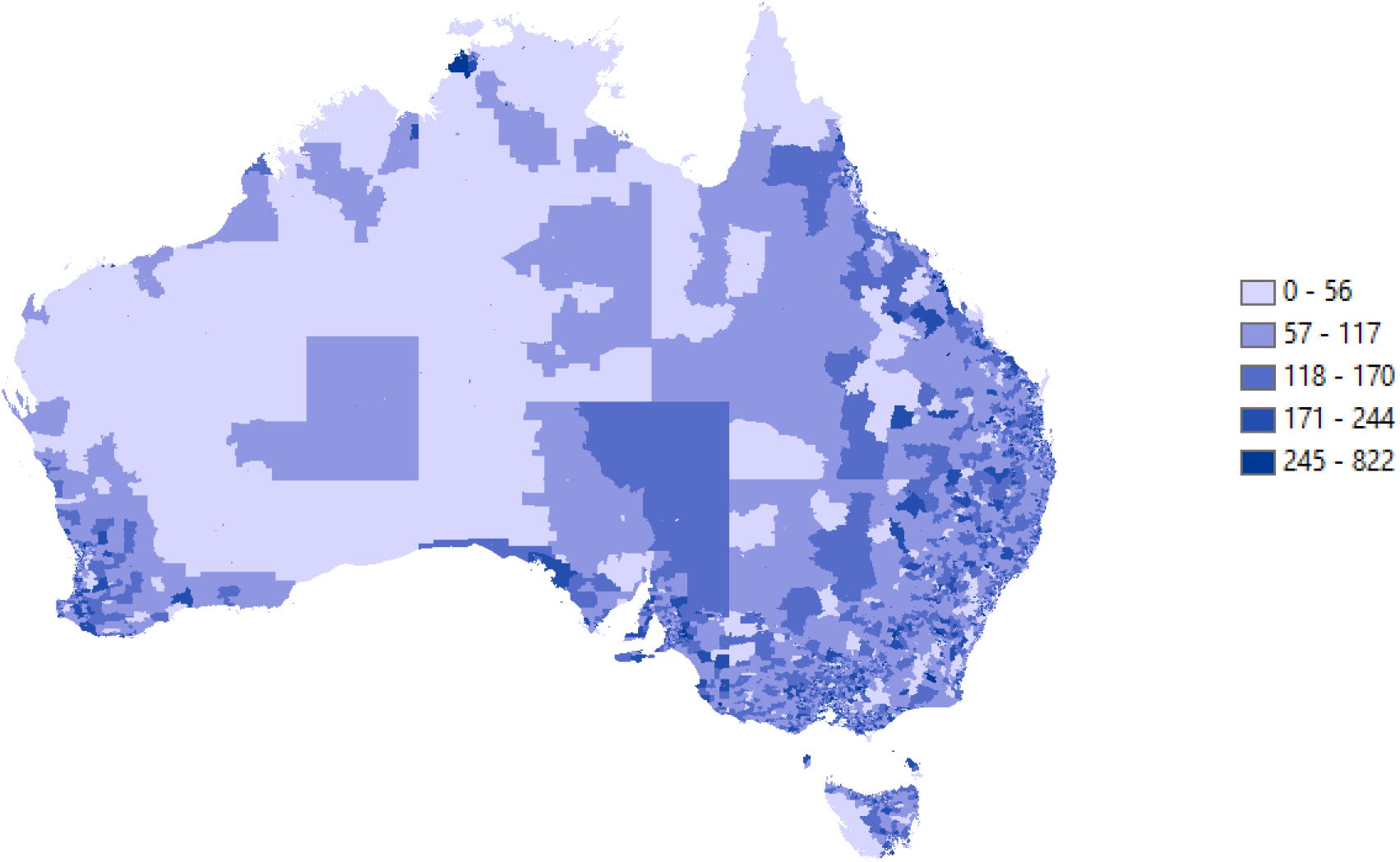
Number of households at SA1 level.

The matrices derived from the ABS data are:

**V***_sc_* = values of commodities by SA4 region and commodity (*n_s_* × *n_c_*)
**A***_sc_* = commodity areas by SA4 region and commodity (*n_s_* × *n_c_*)

These matrices were rearranged to the desired dimensions. There were some gaps in the data regarding the area of trees per SA4 region that had to be filled from alternative sources. This process is explained in Appendix A.

The area matrix derived from the ABARES data is:

**A***_wu_* = Areas of agricultural land uses by Climex cell and for each commodity as defined in ABARES commodities (*n_w_* × *n_u_*).

This matrix was generated by aggregating the 1 km cells of land uses from the ABARES National Map to the Climex cells.

To convert between dimensions two conversion matrices were derived:

**C***_cu_* = conversion matrix between ABS and ABARES commodity groups (*n_c_* × *n_u_*). This matrix consists of 0, 1 entries indicating the columns to which each ABS commodity (rows) belongs.
**C***_sw_* = weight matrix to convert ABS SA4 regions to Climex cells (*n_s_* × *n_w_*). This matrix was produced by overlapping the ABS map and the Climex map.

The matrix of returns per ha by SA4 region (*n_s_* × *n_c_*) is calculated as an element by element division of the value and area ABS matrices:

**R***_sc_* = **V***_sc_* / **A***_sc_*

Which is then converted to return per ha by Climex region and ABARES agricultural land use (*n_w_* × *n_u_*) as:

**R***_wu_* = [**C***_sw_* × (**R***_sc_* × **C***_cu_*)]’

**R***_wu_* and **A***_wu_* are then used to estimate damages as the value of reductions in productive area for each agricultural land use and each Climex cell based on the density of wasp nests on that cell (see below).

### 5.2 Wasp invasion

The wasp spread model is solved for the Climex map based on habitat suitability and growth and dispersal of wasp nests over time. Solution of the spread model for a planning horizon of *T* years results in a matrix of dimensions (*n_w_* × *T*) representing the number of wasp nests per ha for each Climex cell (rows) for each year of the simulation (columns). The wasp nest density for the planning horizon is:

**W***_t_* = matrix of wasp nests per ha over time for one iteration of the stochastic spread model (*n_w_* × *T*). This matrix is created by running the simulation algorithm explained in Section 4.3.

For multiple stochastic runs **W***_t_* matrices are stacked on the third dimension for each random iteration, resulting in a data cube that can be used to derive probability distributions and trajectories of the invasion in time and space.

### 5.3 Market Damages

As explained earlier, market damages caused by wasps include reduction of pollination, direct fruit damage and damage to honey production and apiculture. The data and assumptions used to estimate these damages are explained in Section 3.3. The pollination and fruit damages were defined as column vectors (*n_c_* × 1) for commodities.

**p***_d_* = pollination reliance for the given crop or fruit (0-1).
**f***_d_* = expected proportion of fruit damage caused by wasps (0-1).

These damages were converted to agricultural land use dimensions (*n_u_* × 1) as:

**d***_p_* = (**C***_cu_*)’ **p***_d_*
**d***_f_* = (**C***_cu_*)’ **f***_d_*

Where **d***_p_* and **d***_f_* are proportional damages to pollination and fruit for each of the 27 agricultural land uses from ABARES.

The damages to the honey industry were allocated spatially based on the distribution of land used in the Climex cells defined as a column vector (*n_w_* × 1):

**v***_h_* = value of honey production per Climex cell.

The market damage associated with wasp presence is represented by introducing three damage parameters:

*δ_p_* = proportion of area on which pollination is reduced per wasp nest/ha.
*δ_f_* = proportion of fruit area damaged per wasp nest/ha.
*δ_h_* = proportion of area on which honey production is reduced per wasp nest/ha.

The damages are calculated by multiplying the elements of each row of **A***_wu_* times the corresponding elements of (*δ_p_* **d***_p_*)’ and (*δ_f_* **d***_f_*)’ respectively, to obtain damage matrices **A***_dp_*, and **A***_df_* for pollination and fruit respectively. The damages to the honey industry are calculated by multiplying each column of **A***_wu_* times the corresponding elements of (*δ_h_* **v***_h_*) to obtain the damage matrix **A***_dh_*. These three damage matrices have the same dimensions as **A***_wu_* and represent the area affected for each commodity and in each Climex cell per wasp nest/ha.

Finally, the actual market damages caused by the wasps for the planning horizon *T* are:

**D***_p_* **= (**(**A***_dp_* ° **R***_wu_*)’ **W***_t_*) / 1×10^6^
**D***_f_* **= (**(**A***_df_* ° **R***_wu_*)’ **W***_t_*) / 1×10^6^
**D***_h_* **= (**(**A***_dh_* ° **R***_wu_*)’ **W***_t_*) / 1×10^6^

Where ° represents element by element multiplication as opposed to matrix multiplication. These three damage matrices have dimensions (*n_u_*, *T*), representing the actual damages to each land use in $ million for each of the *T* years (columns) attributed to pollination, fruit damage and honey production losses respectively. These damages are then discounted and expressed as present values for the given planning horizon. The damages can be partitioned spatially (by Climex cell) as well as by land use.

In plain language, the matrix manipulations described in this section involve these steps:

1. Start with ABS commodity tables of dimensions 76 rows (SA4 regions) by 65 columns (commodities) consisting as values ($) and areas (ha) and calculate table of returns per ha (**R***_sc_*).
2. Convert the returns table to dimensions 522 rows (Climex cells) by 27 columns (ABARES agricultural land uses) by applying conversion matrices derived through map overlays and aggregating commodities from ABS to ABARES. The resulting table of returns per ha (**R***_wu_*) is used to calculate damages per ha together with (**A***_wu_*) defined in step 3.
3. Create a table of dimensions 522 rows (Climex cells) by 27 columns (ABARES agricultural land uses) by overlaying Climex map on National Map, resulting in a table (**A***_wu_*) of commodity areas.
4. Simulate the wasp invasion for a planning horizon of *T* years using the algorithm described in Box 1. The result of one random iteration of the model is saved as a table (**W***_t_*) of dimensions 522 rows (Climex cells) by *T* columns (years).
5. Using results from 3 and damage parameters, create tables of dimensions 522 rows by 27 columns containing damages associated with pollination (**A***_dp_*), fruit damage (**A***_df_*) and honey (**A***_dh_*), expressed as area affected per wasp nest/ha.
6. Combine area damages from 5 with simulation results from 4 to calculate trajectories of damages for the *T* years in the planning horizon.
7. Depending on the analysis desired the results from 6 can be aggregated across time, space or commodity. The equations in the text represent the derivation of damage trajectories (**D***_p_*, **D***_f_*, **D***_h_*) for the 27 agricultural land uses (rows) for *T* years (columns).
8. Multiple iterations of stochastic simulations are conducted by random selection of parameter sets from given probability distributions and repeating steps 4 to 7.

### 5.4 Non-market damages

Resources were not available to undertake a primary study to determine the impact of wasps on outdoor activities, human health or nature conservation. Rather, a benefit transfer approach was used. Benefit transfer (BT) incorporates a set of methods for applying previously estimated values from a ‘study site’ to a ‘policy site’ of interest, that is, the area and environment effected by the incursion where no values are currently available. Values from the study site are typically adjusted for differences in income, prices and demographic variables, and scale. The full benefit transfer undertaken in this project (Tait and Rutherford, 2020) appears as Appendix A, and is summarised below. Rolfe and Windle (2014) was selected as the candidate study for the benefit transfer. That study involved a choice experiment (CE) survey around Brisbane residents’ willingness to pay (WTP) for reductions in impacts of the red imported fire ant (RIFA). The benefits of controlling RIFA are largely non-market in terms of avoiding health impacts, maintaining lifestyle and amenity values, and avoiding environmental impacts. The authors applied a CE survey of Brisbane residents to assess WTP for various outcomes (Table 6), defined by reductions in:

- the number of homes affected by stinging events;
- recreation, sporting and school areas affected; and
- protected areas affected.

**Table 6.** WTP estimates from Rolfe and Windle (2014) in 2009 and adjusted for inflation.

| Attribute | WTP/household |  |
| --- | --- | --- |
|  | 2009 | 2020 |
| Reduce number of homes affected | \$0.13/1,000 homes | \$0.16 |
| Reduce recreational, sporting and school areas affected | \$172/100ha | \$215 |
| Reduce protected areas affected | \$0.08/100ha | \$0.10 |

The description of the valuation context presented to survey respondents reveals a good level of consistency with the wasp control benefits considered for the benefit transfer in the current project (see Appendix A, Figure 4). Importantly, Rolfe and Windle (2014) explicitly considered how WTP estimates could be used in BT applications.

The WTP estimates from Rolf and Windle (2014) are presented in Table 6. Respondents to the survey were WTP $0.13 to reduce the impact of RIFA per 1000 homes, $172 per 100 hectares to protect public areas such as recreational, sporting and school areas, and $0.08 per 100 hectares to protect natural bushland. Apart from adjusting the original (2009) values in Table 6 for the effects of inflation, values need to be scaled to suit a larger quantity, population and area than was evaluated by the original study – treating per unit values as invariant to these changes would be incorrect.

The benefit-transfer methodology used to transfer values from the original study to the current context comprises four key components (Figure 15):

1. *Income*: this involves adjusting for differences in income levels between Brisbane households and households in policy target areas;
2. *Policy area*: the area of public land, bushland and number of households over which wasp impact occurs or might occur is defined;
3. *Distance decay:* the rate at which WTP falls with increasing distance from the management area must be defined, with consideration given to whether the chosen distance-decay is the same for each benefit valued; and
4. *Increasing scale*: the effect on WTP of increases in the scale of management outcomes relative to that which was conveyed in the original study is chosen.

**Figure 15.**
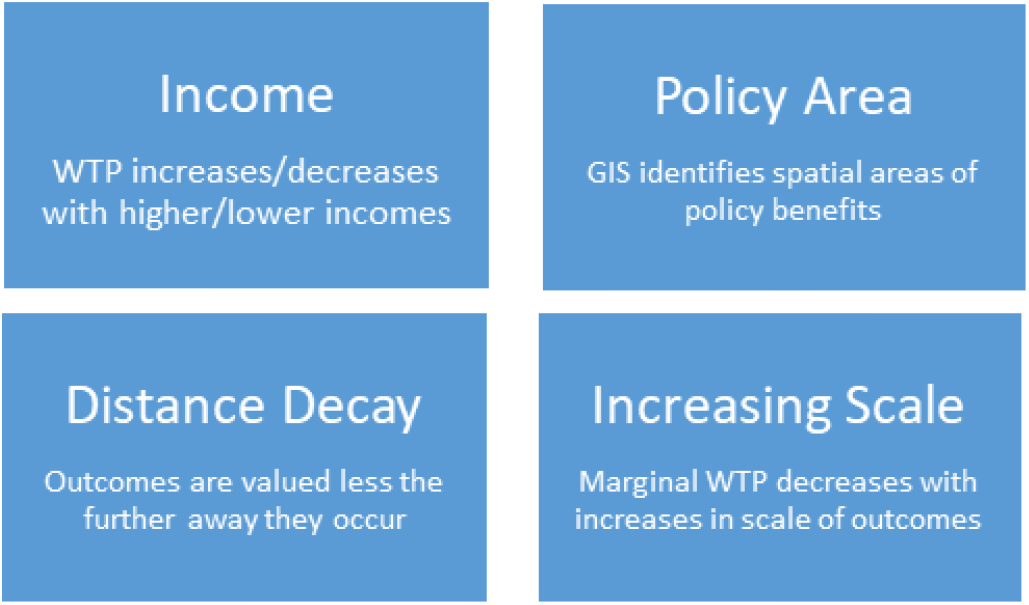
Benefit transfer method, main modules.

Details of steps 1-4 and resulting values are given in Appendix A.

The decision analysis model generates an estimate of wasp nest density at a spatial scale of a CLIMEX cell as explained above. As density increases, so too do the negative outcomes to health, amenity and biodiversity. Control options are modelled within the decision analysis model to affect a reduction in wasp nest density, and it is this change in nest density that is valued in the benefit transfer exercise.

Estimation of damages follows a similar approach to that explained above for market damages, except that in this case there was no need to change the dimensions of matrices from ABS commodities to ABARES land uses. The key matrix in this case is:

**A***_wn_* = matrix containing protected areas (ha), public areas (ha) and number of households for each Climex cell, with dimensions (*n_w_* × 3).

This matrix was created by matching the Climex map with the ABARES National Map and the ABS Statistical Area 1 (SA1) Census map as explained in Appendix A.

The WTP parameters for protection of nature conservation, protection of public areas and reduced household exposure to wasps from Table 6 were applied to the Climex map with the adjustments 1 to 4 above (Appendix 1). This resulted in the damage matrix **D***_wtp_* representing the WTP to avoid damages caused by wasps to protected areas, public areas and households, with the same dimensions as **A***_wn_*. Note that **D***_wtp_* is adjusted by income, policy area, distance decay and scale, as required by the benefit-transfer methodology. In essence, the elements of this matrix represent the aggregate willingness to pay in each Climex cell (rows of matrix) for reduction in non-market damages.

The non-market damage associated with wasp presence is represented by introducing three damage parameters:

*δ_nn_* = proportion of nature-conservation area affected per wasp nest/ha.
*δ_np_* = proportion of public areas affected per wasp nest/ha.
*δ_nh_* = proportion of households affected per nest/ha.

The damages are then calculated by multiplying the elements of each column of **D***_wtp_* times the corresponding elements *δ_nn_*, *δ_np_* and *δ_nn_* respectively, to obtain the adjusted damage matrix **D***_adj_*. Finally, the actual damages depend on the area invaded and the number of nests:

**D***_n_* **= (**(**A***_wn_* ° **D***_adj_*)’ **W***_t_*) / 1×10^6^

This matrix has dimensions (3 × *T*) and represents the non-market damages to protected areas, public areas and households (rows) in $ million for each of the *T* years (columns).

Note that in non-market valuation we are interested in changes produced by a policy rather than on absolute values. The benefit of any given policy is obtained by first solving the model for the baseline case with no biocontrol program, producing **D**_n_(0), and then solving the model for a scenario with biocontrol, producing **D**_n_(1). The non-market benefit from the policy is

**B***_n_* **= D***_n_*(0) - **D***_n_*(1)

Which is the same as avoided damages caused by the policy in scenario (1) in $ million per year. These benefits are then converted to present values for a given discount rate and compared with the costs of implementing the policy.

### 5.5 Costs of biological control

Biological control programmes typically involve significant periods of time and resources to select, test, import, rear and release biological control agents. While time devoted to agent selection is not required in the current context, it is unclear whether further host-specificity testing will be required. Given this uncertainty, the costs of biological control are presented for ‘with’ and ‘without testing’ under the broad categories of research and implementation (Table 7).

**Table 7.** Costs and length of time associated with the biological control programme.

|  | Testing |  | No testing |  |
| --- | --- | --- | --- | --- |
| | Cost (\$'000) | Time (Y) | Cost (\$'000) | Time (Y) |
| **Research costs (fixed)** |  |  |  |  |
| Submission seeking decision on requirement for host testing | 50 | 0.5 | 50 | 0.5 |
| Import agent; establish rearing colony | 646 | 1 | 646 | 1 |
| Host specificity testing | 646 | 1 | 0 | 0 |
| **Total** | **1,342** | **2.5** | **696** | **1.5** |
| **Implementation costs per 250,000 cocoons** |  |  |  |  |
| Mass rearing, release | 2,619 | 4 | 2,619 | 4 |
| Long-term monitoring and impact assessment | 1,220 | >3 y after release; x 5 y | 1,220 | >3 y after release; x 5 y |
| **Total** | **3,839** |  | **3,839** |  |
| **Total costs per cocoon (\$)** | **20.72** | | **18.10** | |

The research stage incorporates fixed costs associated with clarifying the need for testing, importing the agent, host-specificity testing if required. Testing adds an additional year to the research stage and an additional $650,000 (approx.). The implementation stage involves mass rearing and release of agents, and long-term monitoring. In this project we have assumed a 4-year mass rearing and release programme would be capable of producing 250,000 cocoons (R. Kwong, personal communication, July 14, 2020). With an additional 5 years for monitoring, the total costs of the biological programme amount to approximately $20.70 per cocoon with testing, or $18.10 without testing.

## 6. Scenarios and results

To understand the cost and benefits of introducing *S. v. vesparum*, damages from the baseline ‘no control’ simulation are compared with various ‘control’ scenarios reflecting introduction of the biocontrol agent in year 1. The starting point for all simulations reflects 60 years of past wasp spread throughout south-eastern Australia, with only *ad hoc* control undertaken by households and councils as wasp nests are located (see 4.3).

The model is implemented in Matlab language (The Mathworks, 2020). Each scenario is run for 1000 iterations over a planning horizon of 50 years.

### 6.1 Biological control scenarios

Four biological control scenarios were tested. These were based on parameter values for growth, mortality and effectiveness of the agent, as detailed in Figure 5:

a. LL: Low growth and mortality with low effectiveness (*α_B_*=2; *μ_B_* = 0.4; *ρ*_B_ = *φ_B_* = 0.02)
b. LH: Low growth and mortality with high effectiveness (*α_B_*=2; *μ_B_* = 0.4; *ρ*_B_ = *φ_B_* = 0.04)
c. HL: High growth and mortality with low effectiveness (*α_B_*=3; *μ_B_* = 0.5; *ρ*_B_ = *φ_B_* = 0.02)
d. HH: High growth and mortality with high effectiveness (*α_B_*=3; *μ_B_* = 0.5; *ρ*_B_ = *φ_B_* = 0.04)

For each of the above scenarios, the parameter that determines spread velocity of the agent (*γ_B_*) is assumed to have a value of 1, with 337,235 cocoons of *S. v. vesparum* introduced to infested sites in year 1. We assume that all costs of the biological control programme are incurred in year 1 for each biological control scenario.

### 6.2 The impact of biological control

When biological control is effective, damage caused by the European wasp is reduced. The level of reduction will depend on the values of parameters discussed in Section 4.2.1. Figure 16 illustrates the typical pattern of the reduction in damages that takes place for one set of ‘control’ parameter values (blue line), compared to a ‘no control’ scenario (red line). Damages are shown as cumulative distribution functions (CDF), indicating the range of values for damage and the cumulative probability that each value will occur. When biological control is applied, CDFs shift to the left showing that damages will be lower for any given probability band. The mean value of damages following application of biocontrol is also lowered (values not shown). Just how far the ‘control’ CDF shifts to the left will depend on the selected biocontrol parameters.

**Figure 16.**
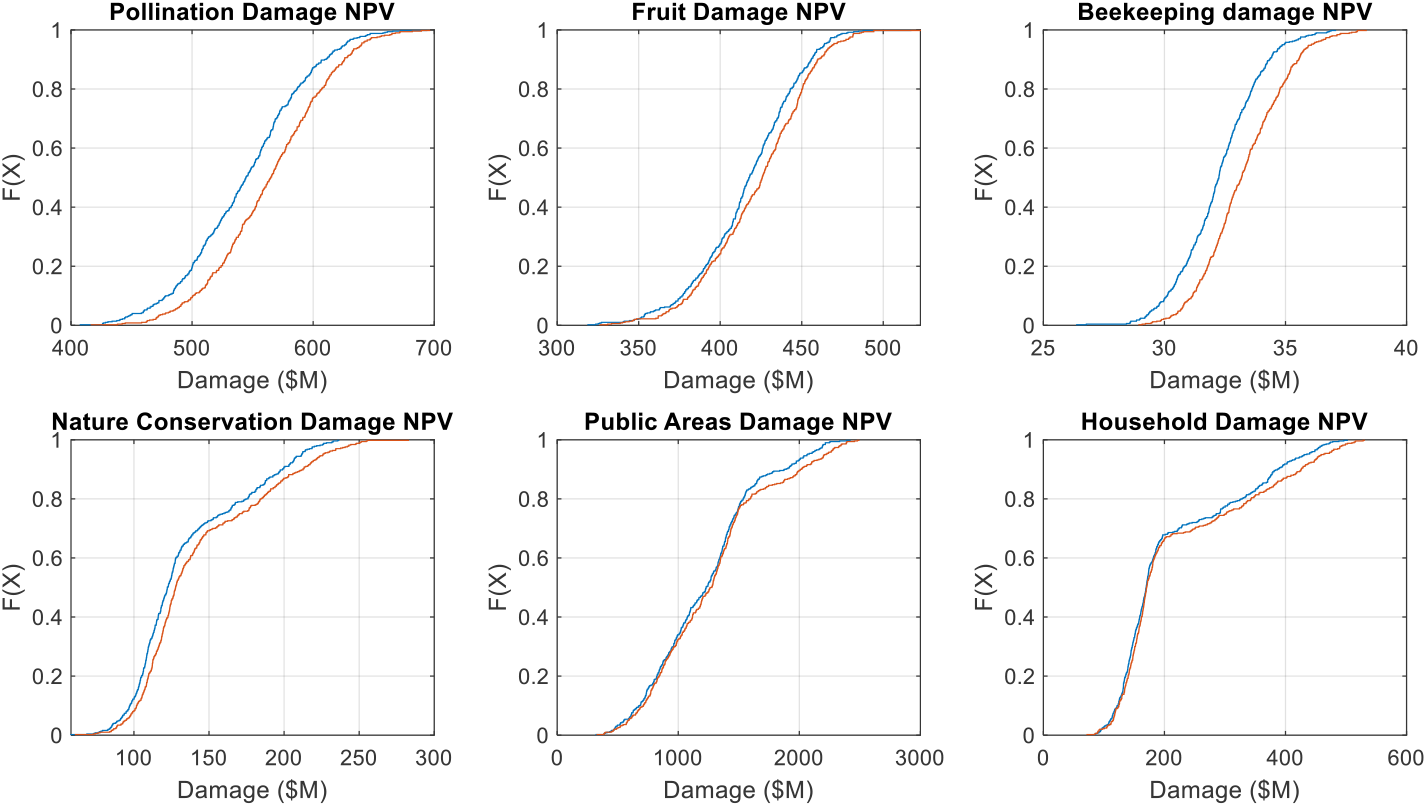
Cumulative distribution functions for damages under the ‘no control’ (red) scenario and for the ‘control’ (blue) scenario which achieves the greatest reduction in damage within the set tested in the base case. The control results are based on the HH scenario.

The mean reduction in damage and 95% confidence intervals from each of the four selected biological control scenarios was compared to the PV of damage under the baseline scenario (Table 8). The mean value of damages from 50 years of wasp spread without control is $2,659 m. Almost half this value is attributed to the damage caused by wasps to outdoor and sporting activities (use of public areas), followed by damage to pollination and ripened soft fruit.

**Table 8.** Reduction in damages (as PV) from the no-control and various biocontrol scenarios with a discount rate of 5% over a period of 50 years.

| | Mean<br>damages (\$m) | Reduction in damage* (\$m) | | | |
| --- | --- | --- | --- | --- | --- |
|  | No control | LL | LH | HL | HH |
| **Market damages** |  |  |  |  |  |
| Pollination | 564 | 2.0 (±3.7) | 5.1 (±3.7) | 14.0 (±3.3) | 20.4 (±3.5) |
| Fruit | 423 | 0.1 (±2.2) | 1.2 (±2.2) | 4.9 (±2.1) | 5.9 (±2.2) |
| Honey | 33 | 0.2 (±0.1) | 0.4 (±0.1) | 0.8 (±0.1) | 1.0 (±0.1) |
| **Subtotal** | **1,020.1** | **2.4** | **6.7** | **19.6** | **27.3** |
| **Non-market damages** |  |  |  |  |  |
| Nature conservation | 143 | 1.6 (±1.7) | 3.1 (±1.7) | 6.8 (±1.8) | 8.1 (±2.0) |
| Public areas | 1,272 | 7.3 (±30.5) | 16.3 (±30.8) | 40.3 (±32.8) | 49.0 (±30.3) |
| Households** | 225 | 2.8 (±4.1) | 4.6 (±4.2) | 10.7 (±4.6) | 10.6 (±5.2) |
| **Subtotal** | **1,639** | **11.7** | **23.9** | **57.8** | **67.7** |
| **TOTAL (\$m)** | **2,659** | **14.1** | **30.6** | **77.4** | **95.0** |
| **Killing costs** | **1,061** | **20.4 (±10.1)** | **30.1 (±10.3)** | **61.1 (±11.2)** | **70.0 (±11.2)** |

\*Numbers in brackets are the 95% confidence intervals.
\*\* “Households” refers to changes in the number of homes affected by stinging events (see Table 6).

Of the four biocontrol scenarios, the largest reduction in damages ($95m) is from scenario HH – high growth and mortality rates of the biocontrol with high effectiveness. The lowest reduction in damages ($14.1m) is from scenario LL – low growth and mortality with low effectiveness.

While there is a large mean reduction in damage to use of public areas there is also a large margin of error. In the case of LL and LH scenarios, the range in possible values would result in some model runs where damage actually increases as a result of the biological control programme. This would suggest there are cases where the low growth and mortality of the biological control agent results in spread of the European wasp across the landscape that is largely or completely unhindered.

Not only does the biological control programme lead to a reduction in market and non-market damages, it also reduces the number of nests that private citizens and public agencies will need to destroy, at an assumed cost of $250 per nest^8^. The value of this reduction (killing costs) is given at the bottom of Table 8. As would be expected, the scenario with the largest reduction in damage (HH) is the scenario with the largest avoided costs for nest control ($70m) as the biocontrol agent has a larger impact on nest density over the 50-year period. These savings to consumers and public agencies represent a loss to pest controllers, which are often small businesses. There could also be environmental benefits due to reduced use of chemicals, which in turn would result in reduced revenues to chemical companies. Given these additional complications we report the avoided control costs separately to distinguish them from unambiguous avoided damages.

The cost of the biocontrol programme for this base case is assumed to be fixed at $6.75m given the number of cocoons released and the fixed cost of the programme. Clearly the benefits of control outweigh the cost of the biological control programme for the biological control scenarios investigated. Sensitivity analysis of these results is undertaken in Section 6.4.

The damages from 50 years of wasp spread without biocontrol are unevenly distributed among state and territories (Figure 17a and Appendix B), reflecting differences in land area and use, population, habitat suitability, and age of the incursion. NSW bears the largest burden of the damage to public areas (78%), nature conservation (63%), households (48%), and honey production (42%) and the NSW public spend the largest amount of money killing wasp nests ($538m). Victoria bears a large burden of the damage to households (46%) and pollination (38%) while the damage to fruit production in South Australia is significant – its fruit producers would bear almost 40% of damage to this industry. Due to its small size only a relatively small amount of damage is experienced by the ACT and this occurs mainly to households and nature conservation. Almost $8m is spent by ACT residents in the control of wasp nests in the base scenario. Queensland is much-less affected, both because the wasp incursion takes time to spread across NSW to Queensland and because the habitat is less suitable in higher latitudes.

**Figure 17.**
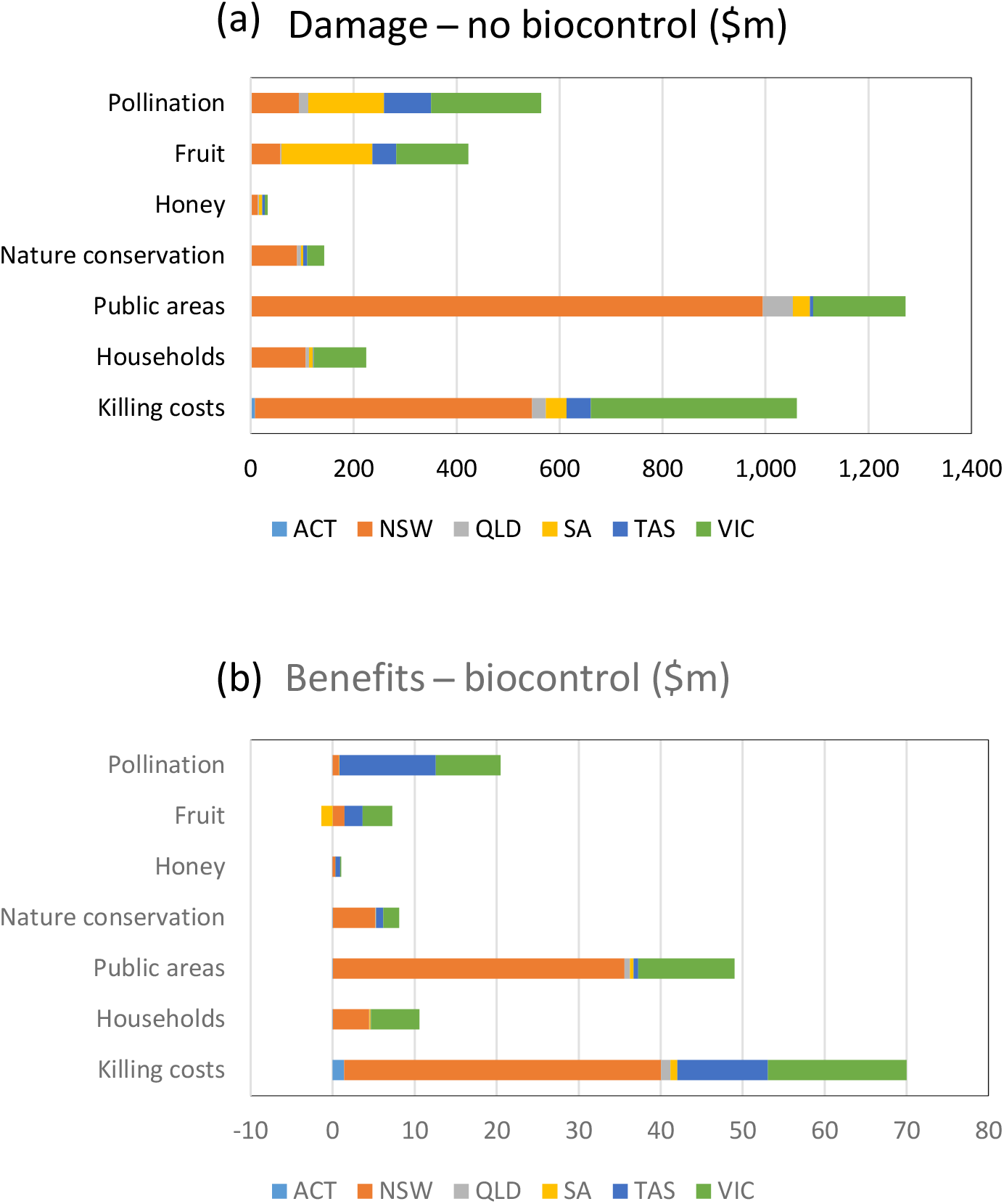
The distribution, by jurisdiction, of (a) wasp damage with no biocontrol and (b) benefits from biocontrol under scenario HH.

The jurisdictions bearing the largest share of damages from spreading wasps will also experience the largest benefits from biocontrol (Figure 17b and Appendix B). Combined, NSW and Victoria receive most of the benefits to households, public areas, nature conservation and fruit production (98%, 97%, 87% and 86% respectively) while honey production, pollination and fruit production in Tasmania are also significant beneficiaries of biocontrol under scenario HH. Interestingly, average negative benefits accrue to fruit production in South Australia and Queensland, and to pollination in Queensland (Figure 17b). This appears to be due to the low proportion of map cells inoculated with cocoons in scenario HH (Table 9).

**Table 9.**
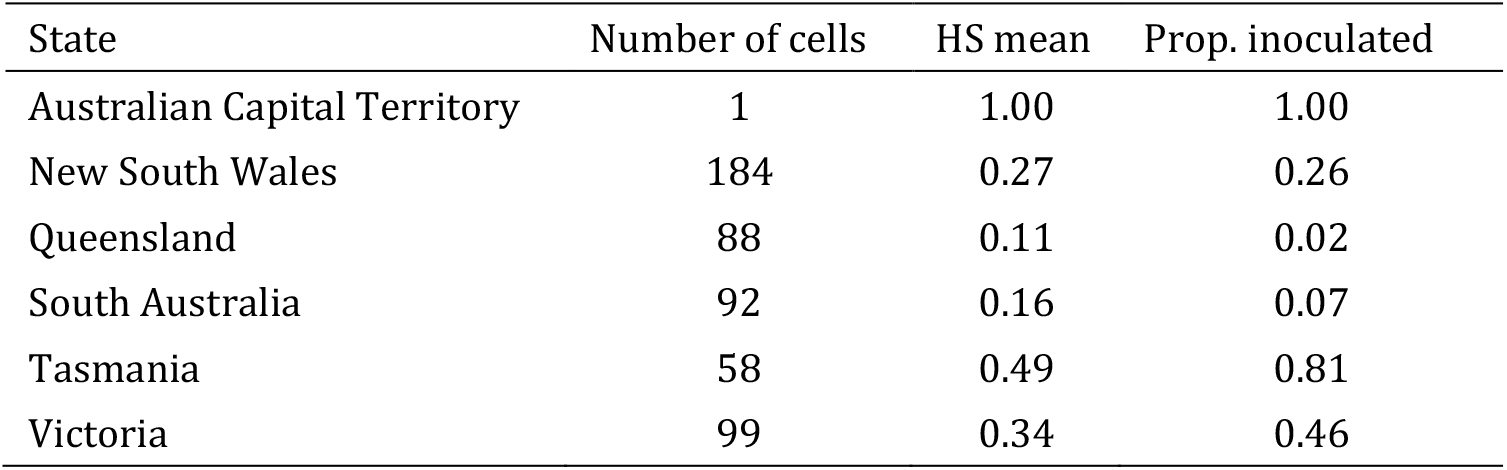
Number of Climex cells per state and mean habitat suitability (HS).

### 6.3 Decision analysis for biocontrol management

The model considers two decision variables for the biocontrol program:

- *x_p_* : the proportion of wasp nests that are inoculated with parasitoid cocoons on a given site.
- *x_c_*: the spatial coverage of the biocontrol release, expressed as the top percentile of infested sites selected for inoculation. For example, a percentile of 90 indicates that only the top 10 percent of sites in terms of wasp-nest density are selected for release of the biocontrol. The larger the percentile the smaller the number of cocoons released.

In the analysis above, the benefit of biocontrol for each scenario was estimated by comparing damages relative to the do-nothing case. In that case four scenarios were selected for comparison based on the assumed population parameters of the parasitoid. The decision variables were fixed at *x_p_* = 0.2 and *x_c_* = 95. In this section we work with Scenario LH and vary the decision variables to study their effects on the benefits and costs of the program. We also go beyond mean values and look at probability distributions.

Figure 18 shows the results of a 5×5 factorial experiment to test the effects of the decision variables within the ranges *x_p_*=(0.1,.., 0.3) and *x_c_*=(95,.., 99). Based on benefits, the best outcome is achieved with *x_p_* = 0.3 and *x_c_* = 96, indicated as point *A* (Figure 18 (a)). This solution requires an initial population of ∼423,000 cocoons (Figure 18 (b)). The option to reduce *x_p_* from 0.3 to 0.1 is indicated by point *B*. This solution requires an initial population of only ∼141,000 cocoons. Moving from *A* to *B* would reduce the number of cocoons required (and hence the cost of the biocontrol program) but would also reduce benefit from $110 million (*B_A_*) to $84 million (*B_B_*) in present value terms for the 50-year evaluation horizon.

**Figure 18.**
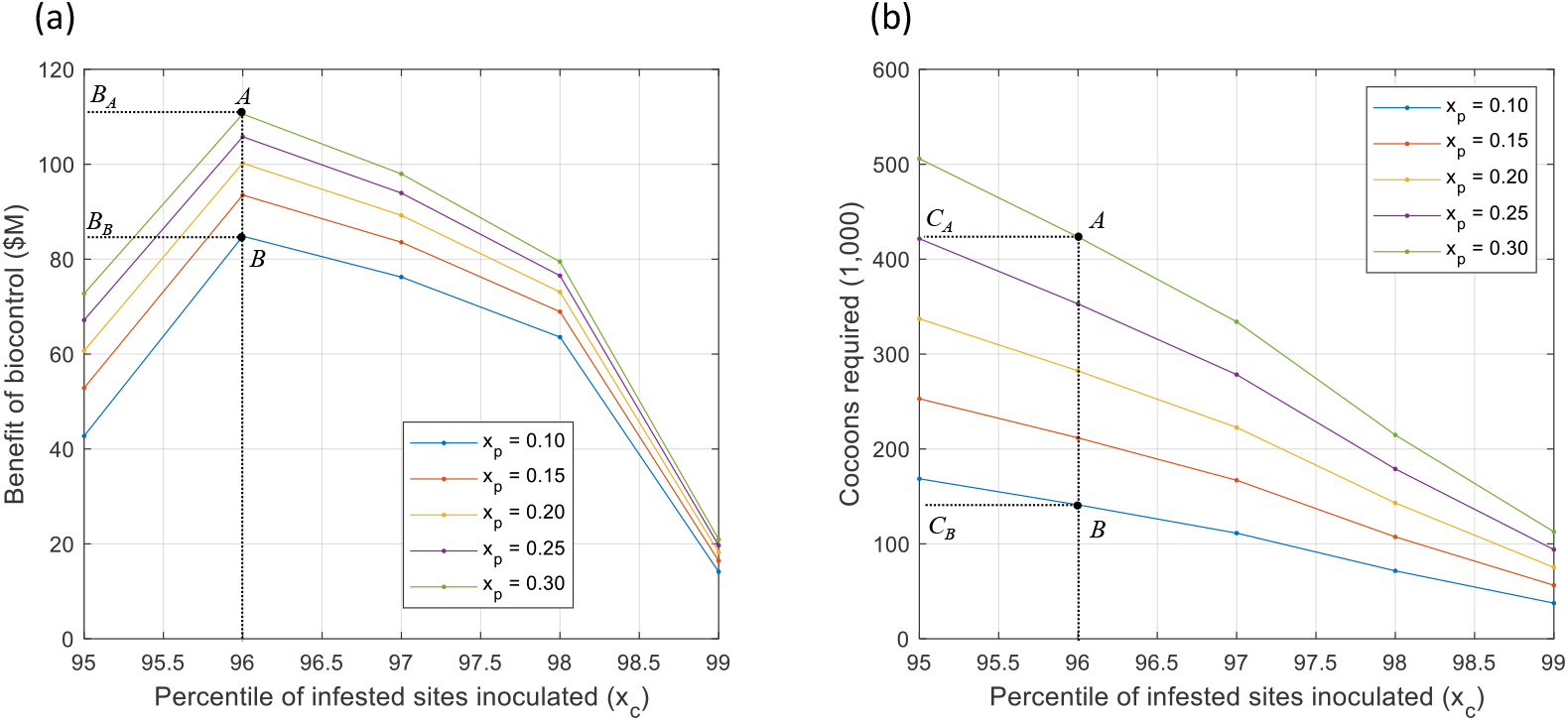
Effects of biocontrol decision variables on the benefits of the biocontrol program (a) and the number of cocoons required at the start of the program (b). Simulation results based on Scenario LH.

Figure 19 presents the full distributions associated with points *A* and *B*. The uncertainty associated with the project is evident by the wide range of values in the horizontal axis. To determine the best option, we need to know the expected cost of the biocontrol program and the likelihood of success. Using a simple scenario, assume that each ‘established’ cocoon costs $20.70, then the biocontrol cost of option *A* would be $8.8 million and for option *B* it would be $2.9 million. If plotted in Figure 19, these costs would be close to the zero-mark given the scale of the x-axis. This means that in about 98% of the simulations, the benefits would have exceeded the cost for both options. Of course, this would be the case only if the biocontrol parameters were those assumed in the LH scenario, and this does not account for the probability that the biocontrol could fail to become established. However, this example illustrates how the evaluation can be conducted as more solid data become available.

**Figure 19.**
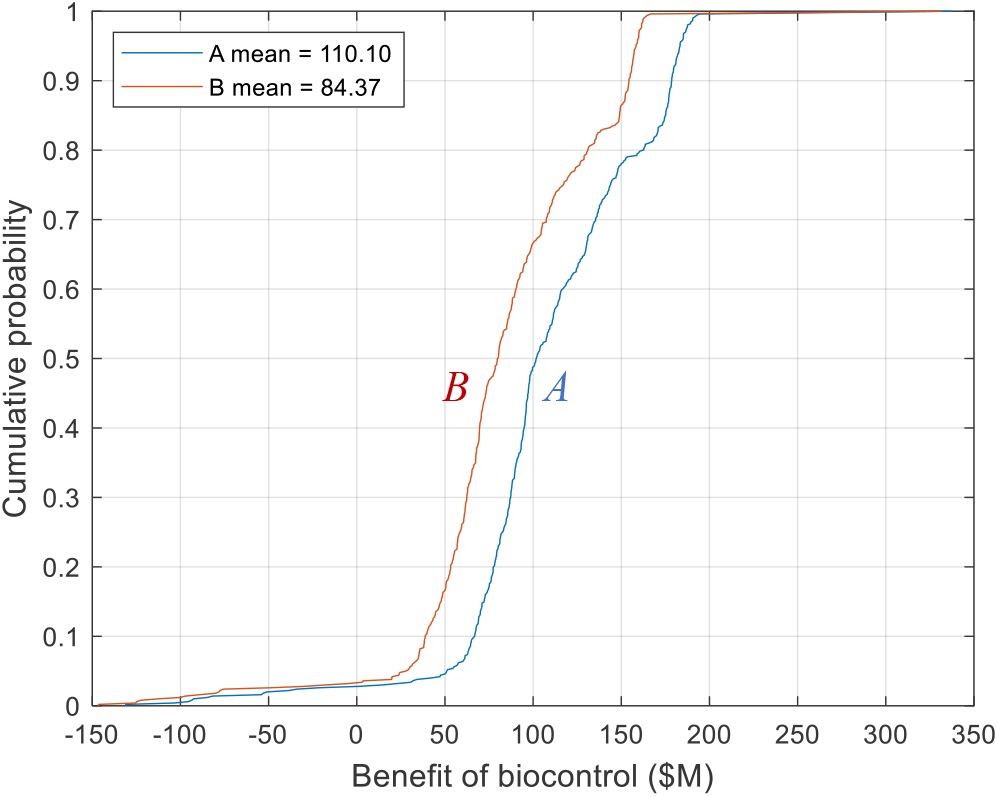
Cumulative probability functions of the benefit of biocontrol for two decision alternatives for (xp, xc): A = (0.1, 96), B = (0.3, 96). Simulation results based on Scenario LH.

Splitting the benefits of biocontrol into private and public (Figure 20), we can see that public benefits are about four times larger (with means of $59 million and $48 million for *A* and *B* respectively) than private benefits (with means of $14 million and $10 million).

**Figure 20.**
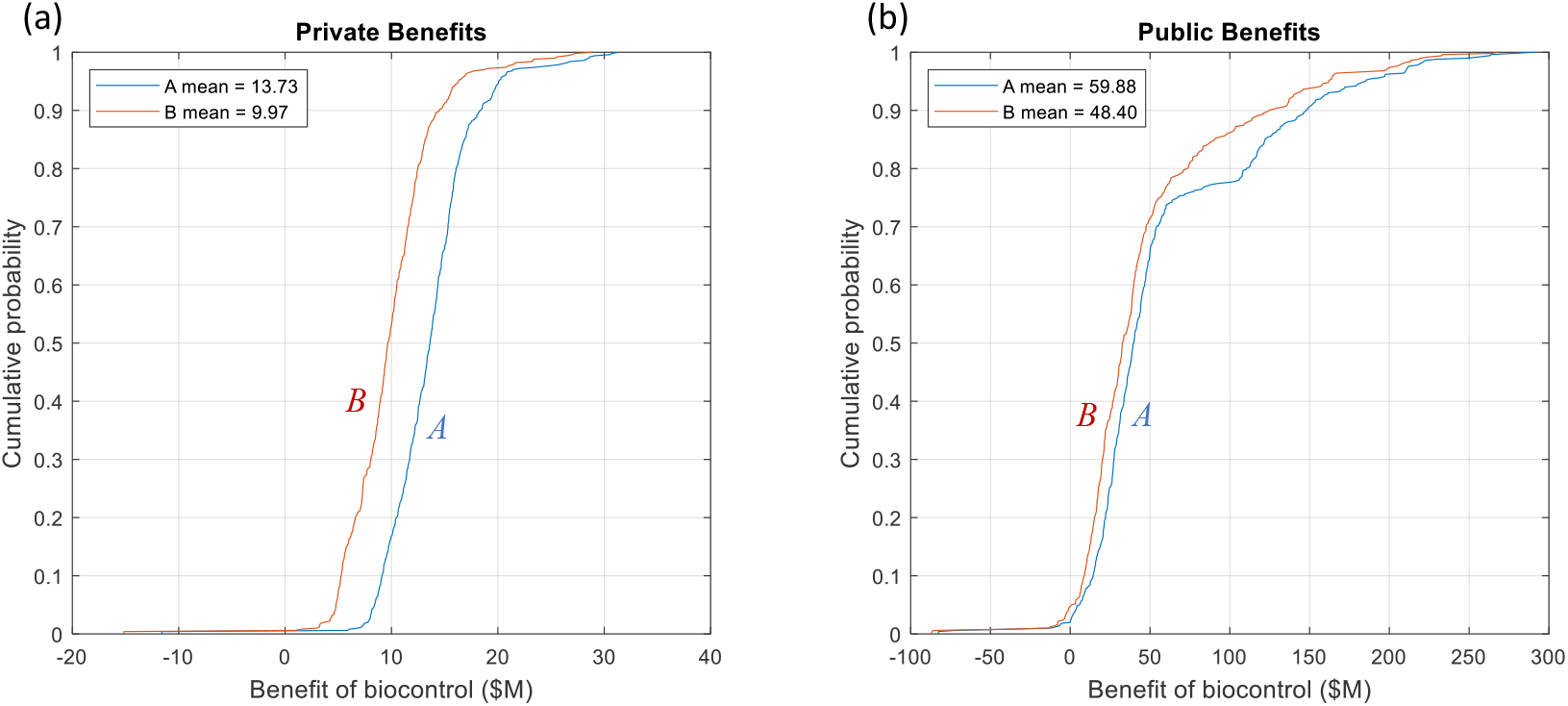
Cumulative probability functions of the benefits of biocontrol split into private (a) and public (b) benefits, for two decision alternatives for (xp, xc): A = (0.1, 96), B = (0.3, 96). Simulation results based on Scenario LH.

Additional analysis is required to understand (1) the likelihood that the biocontrol will be able to achieve the parameters assumed in these scenarios; and (2) the likelihood that the biocontrol agent will become established where inoculated and the associated costs.

### 6.4 Sensitivity Analysis

So far, we have identified ranges of parameter values that would make the biocontrol program feasible and estimated the potential benefits of the biocontrol program under alternative strategies regarding breeding and release of parasitoid cocoons. Given the uncertainty in the ‘true’ values of some of the parameters, in this section we carry out additional tests using an expanded range of strategies for biocontrol release.

For this analysis we focused on the two extreme biocontrol scenarios (LL and HH) and ran an 11 × 11 factorial experiment with *x_p_* in the range (0.05, 0.5) and *x_c_* in the range (90, 99). The full distributions for all the tests under the two scenarios considered, further confirm the uncertainty of the results. There are two interesting patterns in these results: (1) there are many management options that result in benefits > 0, and (2) the probability of benefits > 0 is greater under the HH scenario for all management options, but the distributions are more spread out under the HH scenario (Figure 21b). A break-down of distributions by state and industry is given in Appendix C.

**Figure 21.**
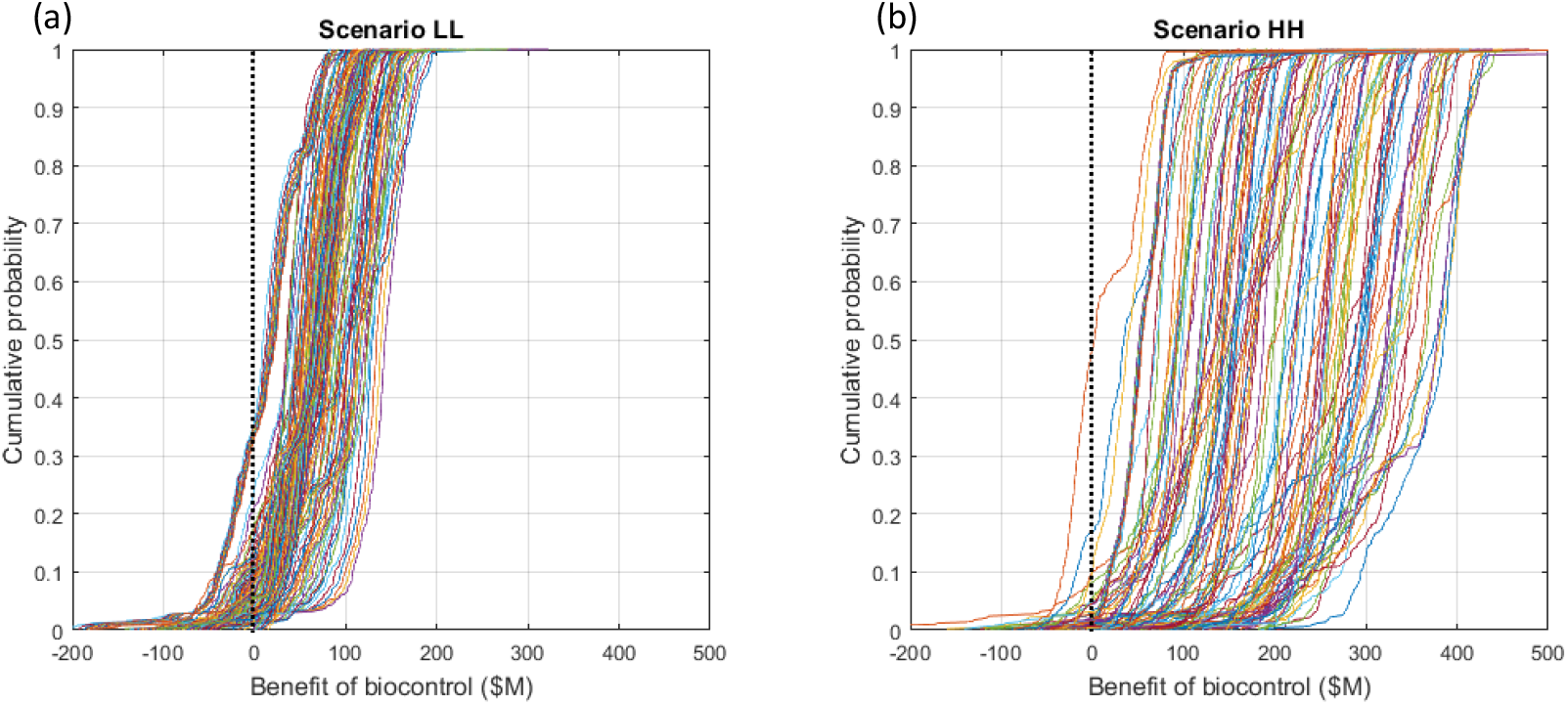
Cumulative probabilities of biocontrol benefits for multiple cocoon release strategies, ranging from (x_p_, x_c_): (0.05, 99) to (0.5, 90) in a full factorial design. Simulation results based on Scenarios LL and HH.

To determine whether supporting the program is justified based on benefit-cost ratios we need to compare benefits with the cost of the biocontrol program. However, the cost differs between the distributions in Figure 21, so we need to assess the benefit of biocontrol as a function of the number of cocoons (Figure 22).

**Figure 22.**
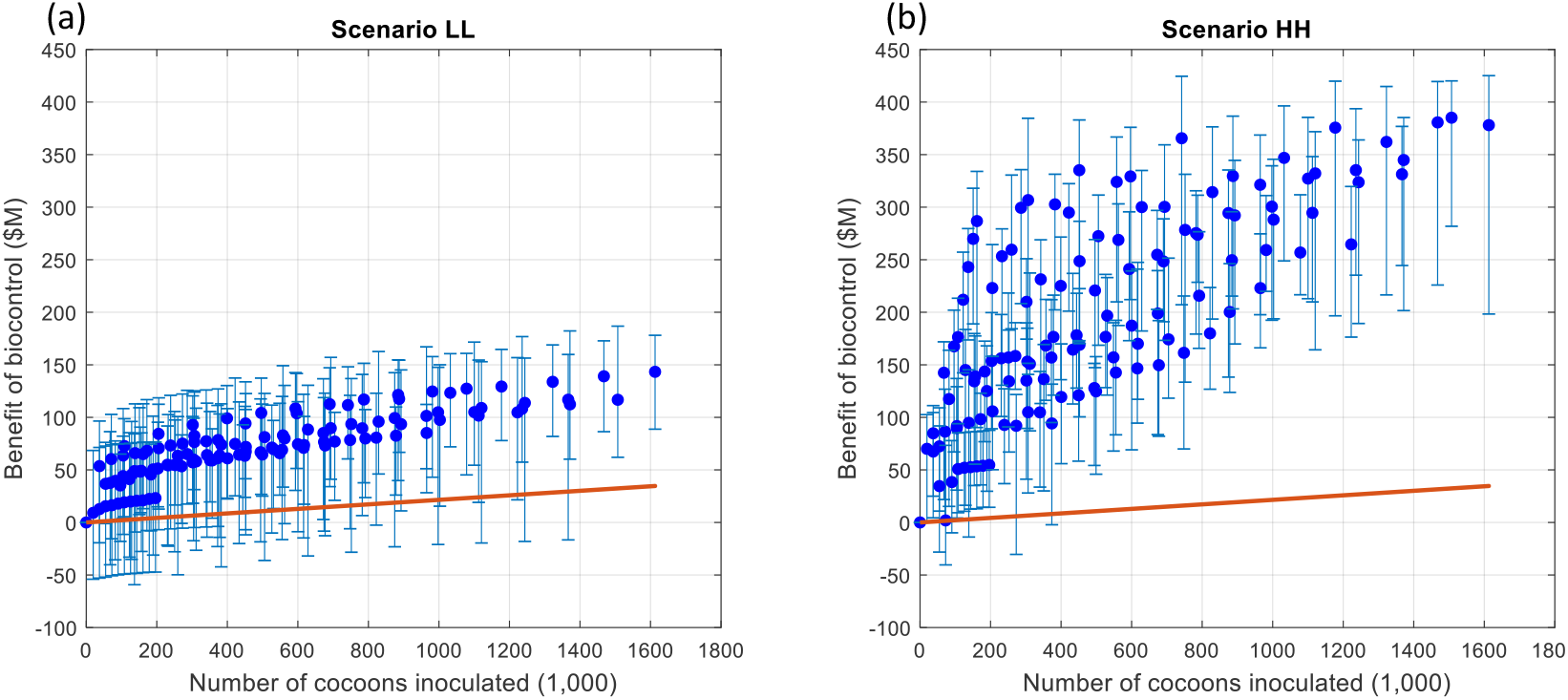
Benefits of biocontrol against the number of cocoons inoculated for multiple cocoon release strategies, ranging from (x_p_, x_c_): (0.05, 99) to (0.5, 90) in a full factorial design. Circles are medians and error bars are the 5th and 95th percentiles of the distribution of simulation results based on Scenarios LL and HH. The red line indicates the cost of the biocontrol programme.

Based on Figure 22, the highest median benefits are $143 million and $385 million for the LL and HH scenarios respectively, whereas the 5th percentiles are $89 million and $282 million. The number of parasitoid cocoons required to achieve these results is 1.6 million. Assuming a cost per cocoon of $20.7 (see Table 7 above) this program would have an estimated cost of $33 million, which is well below the 5^th^ percentile of $89 million under the LL scenario. This suggests that there is a high probability that the benefits will exceed the costs of the program, with benefit-cost ratios of 4.3 and 12.5 at the median, and 2.7 and 9.1 at the 5^th^ percentile, for LL and HH scenarios respectively.

These encouraging results assume that the biocontrol program succeeds (i.e. the parasitoid population becomes established). In terms of Figure 22, what is missing is a confidence band around the cost curve (red line) to represent the likelihood of successful establishment and the potential need for additional expenses to maintain the viability of the biocontrol agent. This is an empirical question that requires more research based on Australian conditions. As before, this also assumes that the parameters in the given scenario reflect reality.

## 7. Conclusion and recommendations

The original key questions are listed below along with comments on how they can be answered with the model developed for this project. But before answering any questions there are two key points to consider:

1. The relative rate of increase of the parasitoid population (*R*) needs to be > 1 for the biocontrol to be able to establish and spread. Evidence from New Zealand suggests that *R* for *S. v. vesparum*, may be > 1 for some sites but, the expected value tends to be relatively low (Barlow et al. 1996). This value is related to two model parameters: the growth rate (*α_B_*) and the winter mortality rate (*μ_B_*) of *S. vesparum* cocoons.
2. There are interactions between growth and mortality of *S. v. vesparum* on one hand, and its effectiveness in suppressing population growth of *V. germanica* on the other. High growth and mortality rates combined with high effectiveness can lead to instability in the population cycles of the two species, with alternating extremes in wasp nest and parasitoid density.

Regarding point 1, **μ*_B_* is a problem, with an estimated value of 0.85 in New Zealand (Barlow et al 1996 and others). Our analysis shows that *μ_B_* needs to be ≤ 0.5 for biological control using *S. v. vesparum* to be feasible. In our analysis we focus on feasible solutions to show the combinations of parameter values that will make the project feasible. This is the first hurdle: if the biological parameters indicate the biocontrol will be unable to establish, there is no need to proceed with the economic analysis. However, studies in NZ indicate that flooding is a leading cause of *S. vesparum* mortality in winter, and this factor may not be as important in Australia. Ultimately, scientific evidence will be required to gain confidence on the likely value of **μ*_B_* and hence on the technical feasibility of the biocontrol. Regarding point 2, biocontrol effectiveness depends on the ability of the parasitoid to reduce wasp-nest growth rates (**ρ*_B_*) and increase wasp winter mortality (**φ*_B_*). No information on these parameters is available for Australia. As before, we selected combinations of values based on their feasibility, in this case defined as being able to achieve a long-term equilibrium between the two species, which is the key to success in classical biological control. Once again, scientific evidence on realistic ranges for these parameters in Australia would help gain confidence on our estimates.

In the meantime, we account for uncertainty as best we can with the information available. Our results suggest that, provided conditions 1 and 2 above for technical feasibility can be met, economic feasibility is high, with benefits likely to exceed costs by a substantial amount in present value terms.

### 7.1 Key Questions

The key questions addressed by this project, and responses based on findings outlined in this project are:

1. **What are the costs and benefits of European wasp control, and how do these vary with pest density and spread?** Pest density and spread are endogenously determined within the model solution, but they ultimately depend on key parameters that have been analysed through a set of scenarios; further scenarios can be designed with advice from scientists. Based on analysing a set of biological control scenarios with two key decision variables, we found classical biological control to be a promising approach to controlling European wasp in eastern Australia. Benefit-cost ratios were between 2.7 and 12.5 for the scenarios analysed. It is important to note that all results assume the biological control agent, *S. v. vesparum* successfully becomes established. Additional research and experiments on agent survival will be required to refine parameters and reduce uncertainty around the benefits and costs of biocontrol.
2. **How much will it cost to reduce the population of European wasps to a particular density, with a particular likelihood of success?** The cost of reducing wasp density depends on i) the number of parasitoid cocoons that need to be released for a population to establish and survive and ii) their spatial distribution. Our model evaluated the cost of biological control for a range of scenarios over a 50-year time frame. As per comment 1, results are promising, but additional research is required on key parameters for both the European wasp and biological control agent.
3. **What are the time frames within which the European wasp could be reduced to a particular density for a given budget?** This question can be answered by placing a limit on the maximum number of cocoons that can be released for a given budget, time trajectories can then be analysed from the model results. Our results indicate the size of the budget to implement a biological control programme could range from as low as $3m to as high as $33m depending on the cocoon-release strategy and the desired benefits.
4. **How effective would the biological control agent need to be in order to reduce European wasp to a particular density over a particular period of time?** The question of effectiveness has been analysed at length above (Section 4.2.1). In summary, to answer this question adequately, additional scientific research is required on the values of the growth rate and winter mortality rate of the biological control agent, and the agent’s effectiveness at reducing wasp-nest growth rates and wasp winter mortality.
5. **How should populations of biological control agents be managed in order to maintain low population densities of European wasps?** Management of the biocontrol agent was represented based on two decision variables: the proportion of nests to be inoculated and the spatial extent of the cocoon release. Values of these variables were analysed in Section 6.3 to show the trade-offs between the benefits of biological control that accrue from releasing a large number of agents, and the costs of such a strategy. The model can be used to analyse additional scenarios once the value of biological parameters becomes clearer.
6. **What magnitude are the measurable non-market impacts, and does their inclusion change the business case?** The non-market impacts of European wasps were estimated to be more than one-and- a-half times the market impacts and, for the scenario analysed in the paper, would amount to $1.6 billion dollars if the wasp continues to spread across the landscape without any formal management programme. This can be compared to market impacts of around $1 billion. Under the biological control programmes assessed in this research, the BCR are therefore significantly higher with the inclusion of non-market impacts.

### 7.2 Biocontrol programmes for other established pests

This study on the economic feasibility of European wasp control could provide useful general information about the control of other pests that are widespread. The following should be useful in this regard:

- For any potential biological programme, scientists must determine whether the relative rate of increase for the biocontrol agent, R in this study, will be > 1. If *R* < 1 for the biocontrol agent, there is no need to undertake further evaluation of its potential benefits, as the organism will be unable to establish a viable population. In this case the main question is whether there are mechanisms for increasing the value of *R*, and at what cost.
- The modelling approach presented in this study could be transferred to other pest-biocontrol agent scenarios. In some cases, this would require modifications to the population dynamics model and calculation of damages (e.g. where pests reproduce more frequently than once per year). Regarding the calculation of damages, the conversion of data from maps to matrices of values that are easily manipulated provide a readily applicable framework for other cases.

### 7.3 Recommendations

The economics of controlling European wasp using the biological control agent *S. v. vesparum* looks promising based on the modelling undertaken in this research. Results, however, ultimately depend on key biological parameters whose values are uncertain. These parameter values need to be refined before a formal recommendation on a biocontrol programme can be made. We therefore recommend the following actions:

1. **Additional scientific research** Additional scientific research and experiments should be undertaken to refine the growth rate and winter mortality rate of *S. v. vesparum*, and its effectiveness at reducing wasp-nest growth rates and wasp winter mortality.
2. **Case studies** Case studies should be undertaken to gain additional insights into key parameter values and the spatial aspects of a biological control programme. A first step would be to analyse the ACT’s eWasp dataset. This dataset is a resource that contains detailed information about nest locations over time, method of detection and stinging events.
3. **Develop closer contact with NZ experts.** Recent research in New Zealand suggests that introducing *S. v. vesparum* collected from the United Kingdom, thought to be the origin of New Zealand’s European wasps, will increase the parasitoid’s effectiveness in controlling the wasp. The newly imported parasitoids are in containment in New Zealand and once approved for release, will be released at multiple locations in the South Island (R. Kwong, personal communication, September 23, 2020). The Australian biocontrol effort could leverage off this research. We therefore recommend that closer contact with the New Zealand biological control team be established in order to benefit from the NZ research and improve our understanding of likely agent performance in Australia.
4. **Investigate other biological control agents** Based on research currently underway in New Zealand, there appear to be several biological control agents that might provide additional control of the European wasp (Ward, 2014; R. Kwong, personal communication, September 23, 2020):

- the mite *Pneumolaelaps niutirani*, first discovered in New Zealand in 2007 but widespread throughout the country (Fan et al., 2016). It has been found in wild nests of the common wasp and European wasps as well as on overwintering queens. Research to date suggests that mite-infested wasp nests are smaller than uninfected nests. However, these mites have also been collected on sticky traps in honeybee hives.
- A mermithid nematode in the genus *Steinernema.* High mortality rates of European wasps were observed where there were high rates of inoculum by the nematode, suggesting this agent may be a candidate for *inundative* biological control.
- *Volucella* hoverflies. These hoverflies are common across the UK and Europe and are known to be associated with the nests of, or to attack, vespid wasps such as the European wasp.

## Acknowledgements

This report is a product of the Centre of Excellence for Biosecurity Risk Analysis (CEBRA). In preparing this report, the authors acknowledge the financial and other forms of support provided by the Department of Agriculture, Water and the Environment, and the University of Melbourne.

The authors are grateful to the following people who generously gave their time to attend project workshops: Raelene Kwong, Greg Lefoe (Agriculture Victoria, DJPR); Con Goletsos, James Woodman, Mona Akbari, Brendon Reading, Catherine Mathenge, Zamir Hossain (ACPPO, DAWE); Milena Rafic, Philippe Frost, Bryce Neville (AECBO, DAWE); Lina Tze, Ian Naumann, Tara Dempsey, Narelle Tyler, Owen Daniel, Ashmeet Malhi (DAWE); Tony Arthur, Kasia Mazur, Belinda Barnes (ABARES); Jim Bariesheff (CoreEnviro Solutions); and Darren Kriticos (CSIRO).

We are also particularly grateful to Raelene Kwong and Greg Lefoe for sharing their knowledge on biological control and the earlier release of *Sphecophaga vesparum vesparum*; to Christina and Jim Bariesheff from CoreEnviro Solutions who shared data from Ewasp and practical knowledge about European wasp control; to Darren Kriticos for providing the CLIMEX dataset; and to Chris Baker who assisted with equation (9). The authors are also grateful to Paul Rutherford for undertaking the GIS component of the modelling.

The authors are also grateful for the insightful comments provided by two external reviewers ‒ Dr David Cook (WA Department of Primary Industries and Regional Development) and Dr Bob Brown (Landcare Research NZ) ‒ on an earlier version of the report. This final report is much improved as a result.

## Appendix A: Valuing Non-market Benefits of European Wasp Management in Australia: a Benefit Transfer Approach

### 1 Introduction

The impacts of the invasive social wasp *Vespula germanica* on human and ecological systems have been researched globally (Beggs et al. 2011. With New Zealand’s beech forests proving particularly suitable habitat, research there has taken a lead in this space (Beggs 2001). In Australia, control of wasp populations can generate a range of benefits to society including for agricultural production, health, recreation, and environmental outcomes (Bashford 2001; Cook 2019). Valuation of control benefits has to date focused predominantly on values to agricultural industries (Cook 2019; MacIntyre and Helstrom 2015) while estimates of non-market values (NMV) of wasp control, such as environmental, have yet to be estimated. When economists approach a valuation exercise they typically assess the types of values, and how they will be estimated, using the Total Economic Value (TEV) framework (Figure 1). Agricultural values are an example of direct-use values, measured using market-prices on foregone production. A range of methods can be used to estimate of non-market values but these are typically resource intensive to apply. An alternative is to conduct a Benefit Transfer, where values from a previously undertaken primary study are transferred to a new subject site where values are required to be generated. This method can take values estimated from any of the other valuation approaches and is useful because it may require less resources to undertake compared to a primary study. This report’s central objective is to apply BT to provide economic values of non-market benefits of wasp control to Australian households, that can be compared against control costs and market-based values.

This report contributes to a project developing methods for prioritisation of biocontrol efforts in Australian (Hester 2019). The BT method developed here is designed to fit within the practical framework of a Decision Analysis Model (DAM) that is used to assess wasp management feasibility (Oscar and Hester 2019).

**Figure 1.**
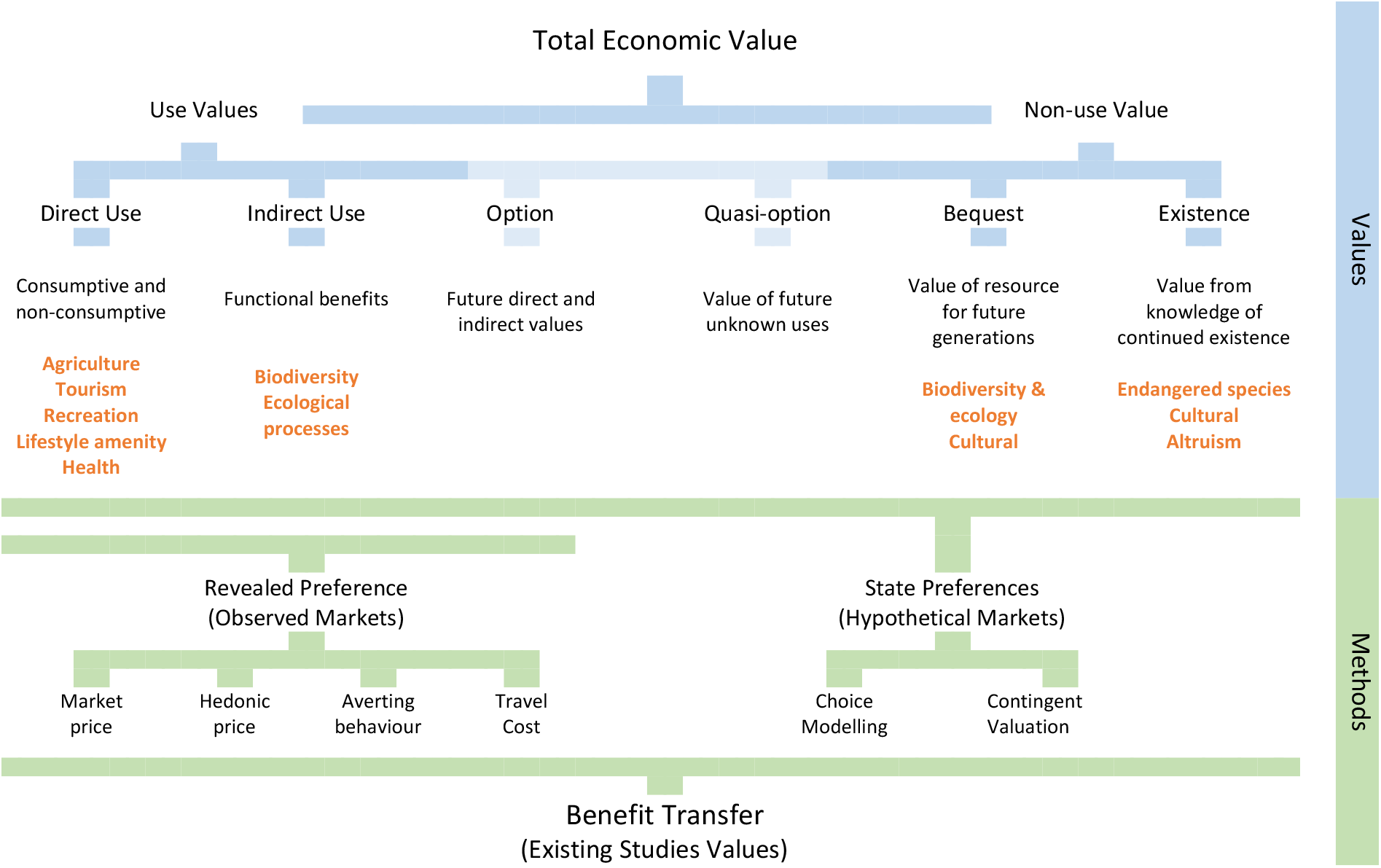
Economic values and relevant valuation approaches

### 2 Benefit Transfer Steps

Benefit Transfer (BT) is a set of methods for applying previously estiamated values from a ‘study site’ to a ‘policy site’ of interest, that is, the area and environment effected by the incursion where no values are currently available. The main caveat to using BT is that, given the current limited availability of suitable source studies, estimates of WTP are unlikely to achieve equivalence with conducting a primary valuation study. The main steps in conducting a BT are described in Figure 2. The remainder of this report describes these steps in developing the BT method applicable to wasp management in Australia.

**Figure 2.**
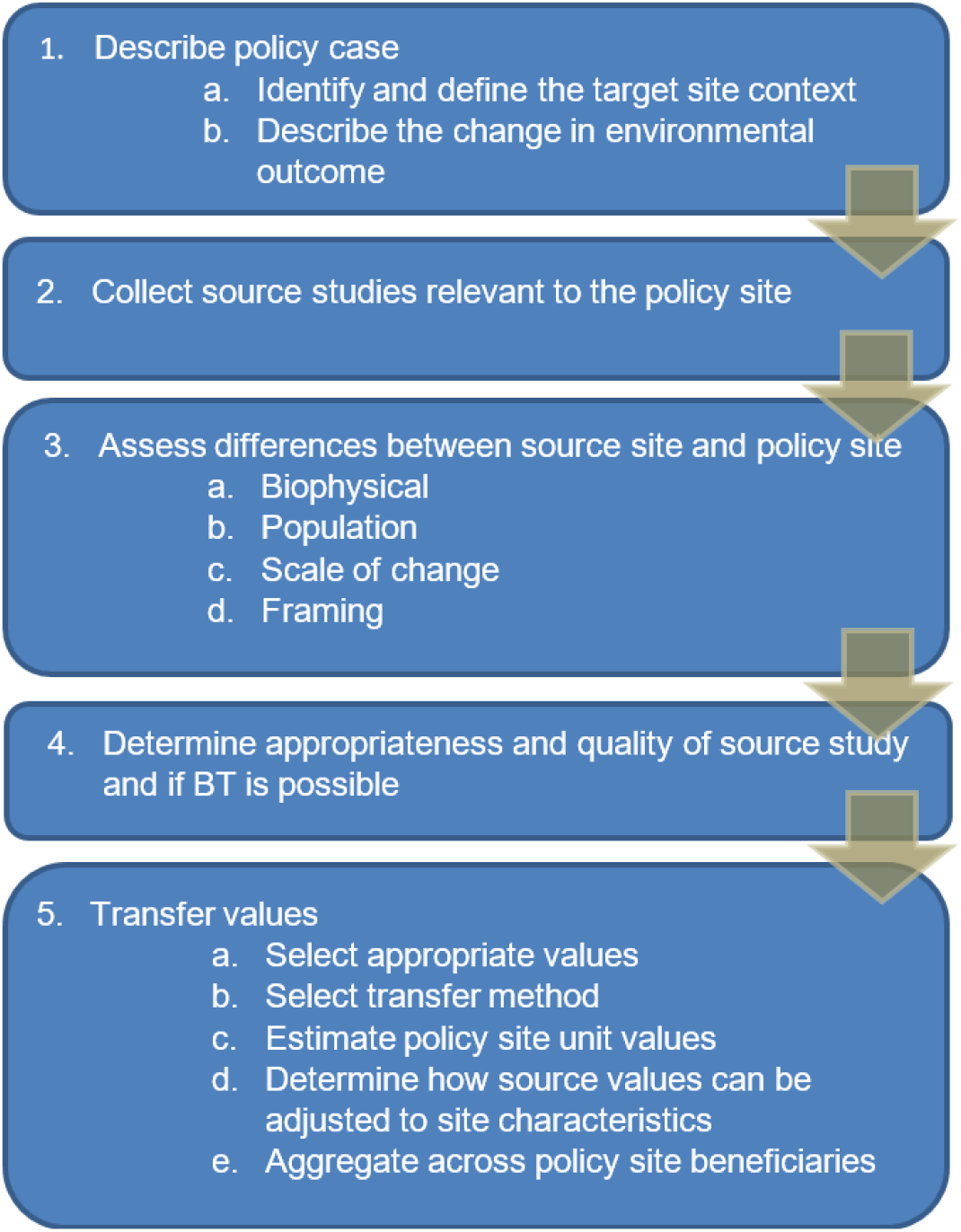
Main steps in Benefit Transfer process

### 3 Benefit Transfer Application

#### 3.1 Target Site and Non-market Benefits of Management

The spatial extent of management outcomes will be defined by the spread dynamics component of the decision analysis model (DAM). This uses CLIMEX modelling of habitat suitability (de Villiers et al. 2017) (Figure 3) as well as other variables. This means that the spatial resolution of the decision analysis model is defined by the size of each CLIMEX cell (50km x 50km). The DAM estimates wasp hive density on a one-year time step. To be appropriately incorporated into the DAM, design of the BT method needs to consider how values change spatially in relation to the management outcomes modelled, and how this can be accommodated in the transfer.

**Figure 3.**
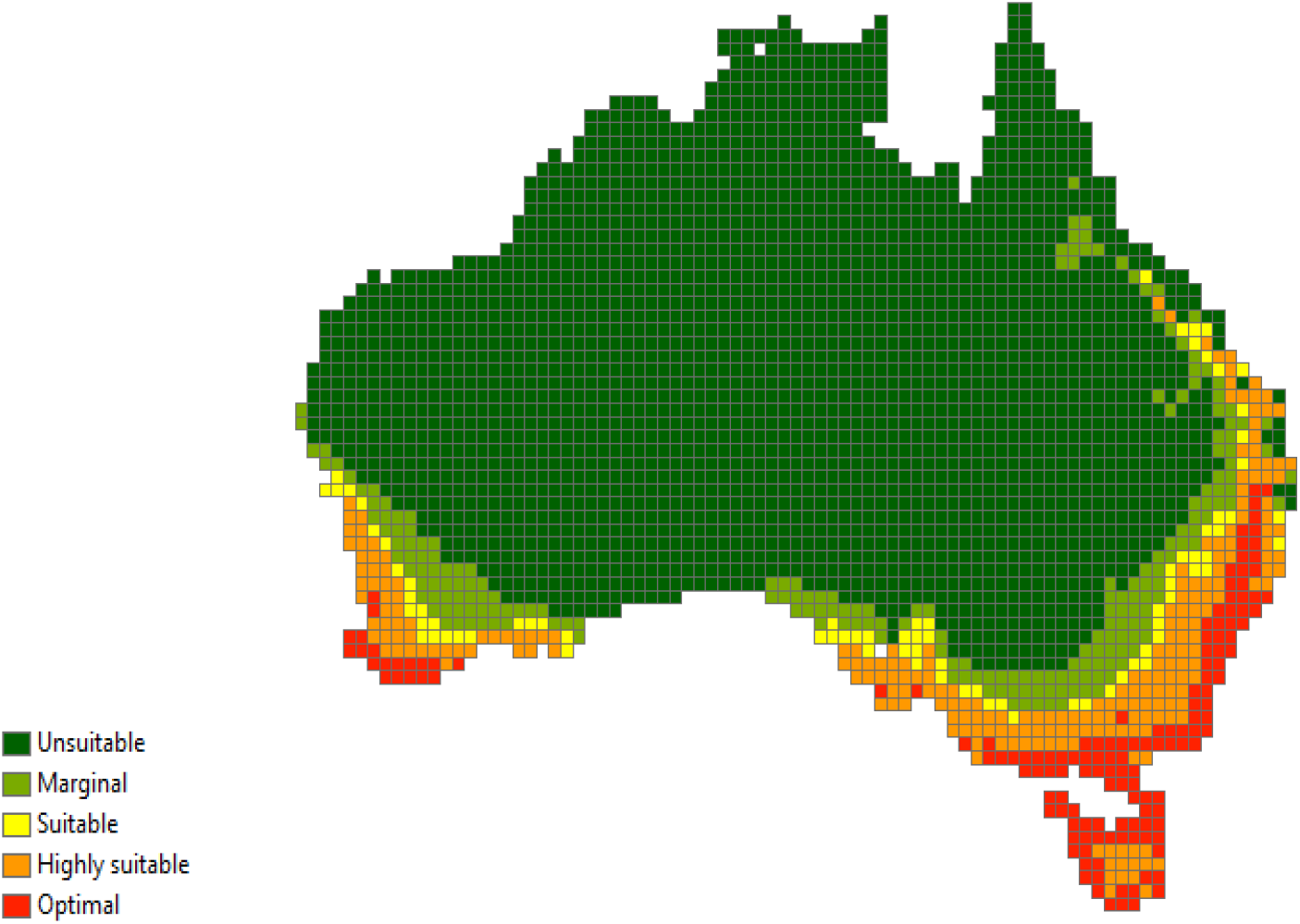
CLIMEX modelling of Wasp habitat suitability

Management of *Vespula germanica* is expected to provide two main types of non-market benefits.

- Due to their scavenging habits and choice of nesting sites, they are frequently attracted to homes, recreation areas, gardens and picnic areas, where they may pose a **sting threat and/or create nuisance** (McGain et al. 2000; Perez-Pimiento et al 2007; Welton et al, 2016).
- In common with all other social wasps, the workers catch various insects and spiders which are malaxated (chewed up) and fed to the larvae. **Biodiversity impacts** on Dipterra (winged insects), Araneae (spiders) and Lepidoptera (butterflies). (Madden 1981; Kaspar 2004; Potter-Craven et al. 2018).

There is also the possibility of ecological and further biodiversity impacts on species that compete with wasps for these insect food sources. While this has been ecologically established in the New Zealand beech forest context, equivalent analysis is not available for Australia and so the scope of biodiversity impacts is restricted to immediate impacts.

#### 3.2 Selection of Candidate Source Studies

The next step is to find relevant primary studies that provide marginal values estimates representing the types of values identified. This follows a literature review process based on indexed database searches. This was undertaken through the Lincoln University Library and covered a range of relevant databases including EconLit, IDEAS, Google Scholar, AgEcon, ABI/Inform. The following specialised databases were also searched, including those dedicated to non-market economic valuation:

- Environmental Valuation Reference Inventory www.evri.ca
- Envalue www.environment.nsw.gov.au/envalue
- Review of Externality Database www.red-externalities.net.
- New Zealand Non Market Valuation Database http://selfservice.lincoln.ac.nz/nonmarketvaluation
- EconPapers http://econpapers.repec.org
- Biodiversity Economics www.biodiversityeconomics.org
- Environmental & Cost Benefit Analysis News http://envirovaluation.org
- The Economics of Ecosystems and Biodiversity Valuation Database www.teebweb.org
- Australian Bureau of Agriculture and Resource Economics Publication Library https://daff.ent.sirsidynix.net.au/

The main criteria applied to select candidate primary studies are listed below. Essentially, the search is for a primary valuation study that comes as close as possible to the context of wasp management being considered in the DAM. A central theme in this process is the concept of commodity consistency. The basic notion is an assessment of whether what has been valued in the primary study, is consistent with what is trying to be valued in the BT exercise. This includes considering how management attributes are characterised and changes defined, the framing of value elicitation, increases in scale and/or scope, demographic representation, and biophysical differences in policy and site contexts.

- The biophysical conditions in the source case is similar to those in the target case
- The scale of environmental change considered in the source approximates the target case
- The socioeconomic characteristics of the population impacted by the change investigated in the source approach those of the target population
- The frame or setting in which the valuation was made at the source is close to that of the target
- The source study has to have been conducted in a technically satisfactory fashion

Expanding on the last criteria, candidate studies must also be of high quality to be reliably used. The quality of candidate studies is assessed based on consideration of:

- Detailed and transparent reporting of data and methods
- Detailed reporting of site and population characteristics
- Quality of underlying biophysical data for modelling
- Restrictiveness and realism of assumptions
- Clear specification of goods and quantities/qualities
- Empirical methods and development (e.g., use of accepted valuation methods)
- Modelling detail (i.e., model includes all elements suggested by theory)
- Data collection methods
- Sample sizes and representativeness
- Statistical techniques and model specifications
- Evidence of selectivity bias
- Robustness of results
- Evidence of peer review or other recognized quality indicators

A single study was found that focused specifically on attributes of wasp management, at Lake Rotoiti in the upper South Island of New Zealand (Kerr and Sharp, 2008; 2013). Lake Rotoiti is the main entrance point to the Nelson Lakes National Park with the main campground on its shores and facilities located in the small adjacent settlement of St Arnaud. The park is approx. 1,000km^2^ and is popular for camping, tramping and anglers, and also contains another large lake, Lake Rotoroa. Beech forest surrounds contain a network of tracks ranging from short nature walks to multi-day back-country tramping. The area experiences some of the highest wasp densities in New Zealand, and has been the focus of detailed ecological studies and control efforts. Estimation of WTP for non-market benefits of wasp control used a Choice Experiment study with identification of control benefits based on consultation with wasp and wildlife management experts, reviewed literature on wasp ecology, wasp management and Lake Rotoiti conservation reports, and two focus groups (Table 1).

The study area experiences well known problems with wasp impacts, this framing effect may elevate preferences and values for reduced impacts relative to other areas not as impacted. This view comes through in the CE design with a relatively high probability level of 20% being selected as the status-quo level in choice sets presented to respondents. The estimates of WTP relevant to the BT are for reducing sting probability (Table 2). that could be linked to wasp hive density in the DAM. However, the WTP estimate of $5.25 should be interpreted as a 1% reduction in probability from the 20% probability base level, and it is not evident how to adjust WTP for probability changes over a lower range of outcomes.

**Table 1.**
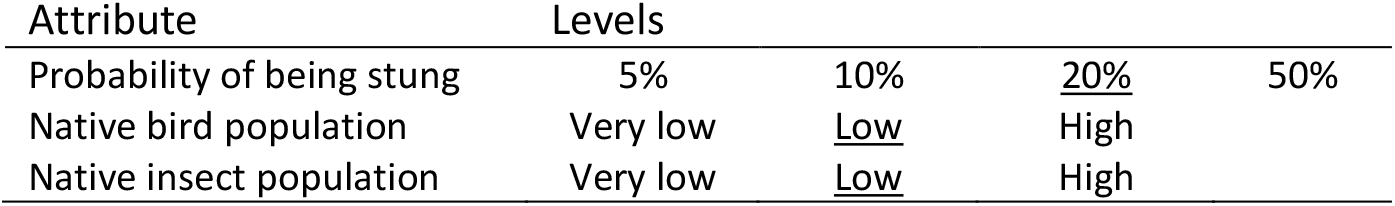
Attributes and levels for Kerr and Sharp (2008; 2013)

The CE was administered during two meetings held with a primary school community in Christchurch and Nelson. It is this aspect of the study that prompts the authors to recommend caution in using their results to draw inference on wider public preferences. The samples were not designed to be representative of the broader community and so are not suitable for drawing inferences about community WTP. The samples were drawn from Nelson which is about 100km distance away, and Christchurch at about 350km distance however no indication is provided as to what proportion of survey respondents were direct users of the lake.

**Table 2.** WTP estimates from Kerr and Sharp (2008)

| Attribute | Annual MWTP/household (sd) <sup>1</sup> |  |
| --- | --- | --- |
|  | Nelson | Christchurch |
| Reduce probability of being stung by 1% | \$5.25 (\$0.31) | \$7.22 (\$0.68) |
| Avoid native bird population becoming very low | \$324 (\$26.09) | \$256 (\$29.32) |
| Low to High native bird population | \$120 (\$8.00) | \$125 (\$13.91) |
| Avoid native insect population becoming very low | \$197 (\$14.14) | \$120 (\$19.19) |
| Low to High native insect population | \$87 (\$7.72) | \$101 (\$14.95) |

Search criteria that included ‘insects’ revealed a small non-market valuation literature applied to benefits of mosquite management. One of these is the only study found that estimates values for reductions in *‘daytime biting nuisance’* levels, an attribute relevant to wasp management benefits. In a CE study conducted in 2015, Bithas et al. (2018) estimate the WTP of residents of Athens, Greece, for non-market benefits of mosquito management. Selection of relevant attributes was initially informed by management experts, and further developed through two rounds of survey pilot testing, using online and face-to-face modes (Table 3).

**Table 3.** Attributes and levels for Bithas et.al. (2018)

| Attribute | Levels |  |  |  |
| --- | --- | --- | --- | --- |
| West Nile virus risk |  | <u>High</u> | Average | None |
| Tiger mosquito health risk |  | Extra measures | <u>No measures</u> |  |
| Biting nuisance - Night | Intolerable | <u>No improvement</u> | Average nuisance | Low nuisance |
| Biting nuisance - Day | Intolerable | <u>No improvement</u> | Average nuisance | Low nuisance |

Face-to-face interviews were also used to administer the final survey, yielding 458 responses. The sampling process generated a representative sample of the Athens population regarding age and gender, while deviating regarding education, income and employment. With a population of approx. four million in an area of just under 3,000km^2^, Athens is one of the world’s oldest cities, and has a global influence. The estimates of WTP relevant to the BT are for changes for changes in nuisance (Table 4) that could be linked to wasp hive density in the DAM. The authors consider these values to be relatively low, and insignificant in some model specification. They reason is that this result is a reflection of the low day-time nuisance reported by survey respondents, with much higher levels of nuisance reported for night time.

**Table 4.** WTP estimates from Bithas et al. (2018)

| Attribute |  | Annual MWTP/household |
| --- | --- | --- |
|  | Intolerable to low | €7.98 (€1.4-€14.5) |
| Improvement in daytime biting nuisance | High to low | €5.32 (€0.96-€9.68) |
|  | Average to low | €2.66 (€0.48-€4.84) |

The third study included in the candidate list focused on an ongoing red imported fire ant (RIFA) incursion in Brisbane, Australia where significant private and public effort continues to be invested (Rolfe and Windle, 2014). Brisbane is Australia’s third largest city with a population of approx. 2.5 million spread over 16,000km^2^. The authors consider that the benefits of controlling RIFA are largely non-market benefits in terms of avoiding health impacts, maintaining lifestyle and amenity values, and avoiding environmental impacts. They apply a CE survey of Brisbane residents using management outcomes defined by health apply a CE survey of Brisbane residents using management outcomes defined by health benefits in people’s homes, recreation in public areas, and environmental impacts in native bushland. Changes in the levels of outcomes are described by the number of houses affected, and hectares of area affected respectively (Table 5). The frame of the CE survey was aligned with modelled infestation and spread scenarios, including modelling assuming no further control efforts. The attributes in the CE were also explicitly matched to the outputs of these biological spread models.

**Table 5.** Attributes and levels from Rolfe and Windle (2014)

| Attribute | Levels |  |  |  |
| --- | --- | --- | --- | --- |
| Number of homes affected | 17,000 (1%)<sup>1</sup> | 83,000 (5%) | 167,000 (10%) | <u>500,000 (30%)</u> |
| Recreational and school areas affected | 250ha (1%)<sup>2</sup> | 1,250ha (5%) | 2,500ha (10%) | <u>7,500ha (30%)</u> |
| Protected areas affected | 12,000ha (5%)<sup>3</sup> | 24,000ha (10%) | 36,500ha (15%) | <u>73,000ha (30%)</u> |

<sup>1</sup>Percent of homes in Brisbane. <sup>2,3</sup>Percent of relevant areas in Brisbane

Description of the valuation context presented to survey respondents is shown in Figure 4. This reveals what can be considered as a reasonably good level of consistency with the wasp control benefits considered for the BT. Unlike other studies included in the candidate list, the authors of this study have explicitly considered how WTP estimates could be used in BT applications (Figure 4).

**Figure 4.**
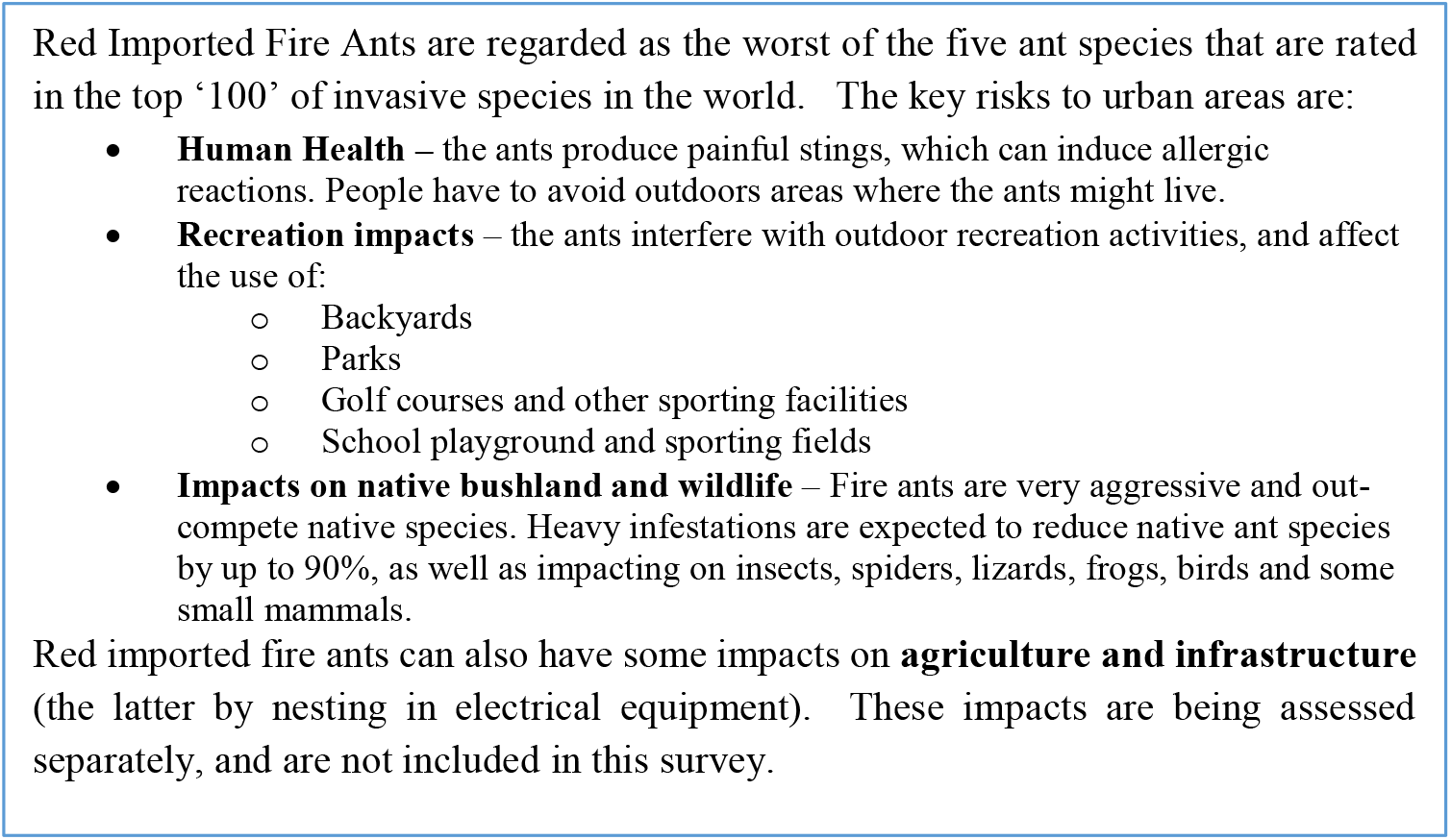
Information shown to respondents in RIFA study (Rolfe and Windle 2014)

**Figure 5.**
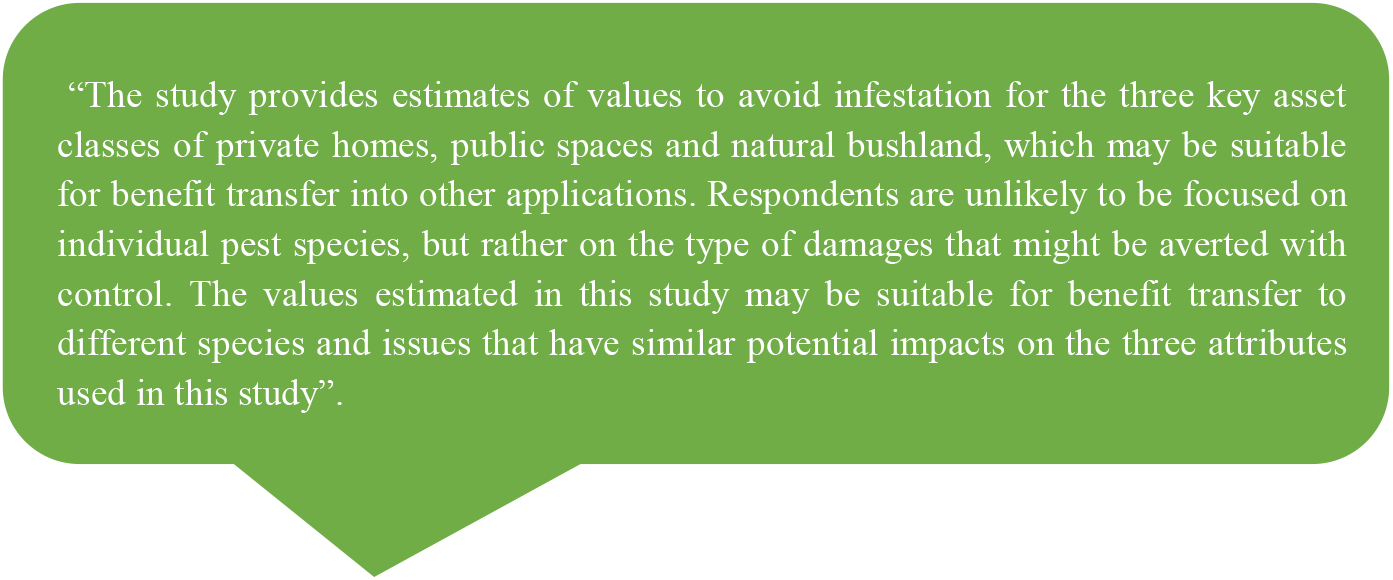
Design of study with future BT consideration in Rolfe and Windle (2014)

This approach has informed the definitions of the management attributes used, so that WTP estimates may be applied at differing scales more readily (Table 6).

**Table 6.** WTP estimates from Rolfe and Windle (2014)

| Attribute | Annual MWTP/household<sup>2</sup> |
| --- | --- |
| Reduce number of homes affected | \$0.13/1,000 homes (-\$0.17:\$0.37) |
| Reduce recreational, sporting and schools areas affected | \$172/100ha (\$0.52:\$250) |
| Reduce protected areas affected | \$0.08/100ha (-\$0.01:\$0.13) |

A summary of candidate study suitability is given (Table 7) where each study is scored qualitatively against overall criteria, from one indicating a study relatively *unsuitable*, and five relatively *suitable*.

**Table 7.**
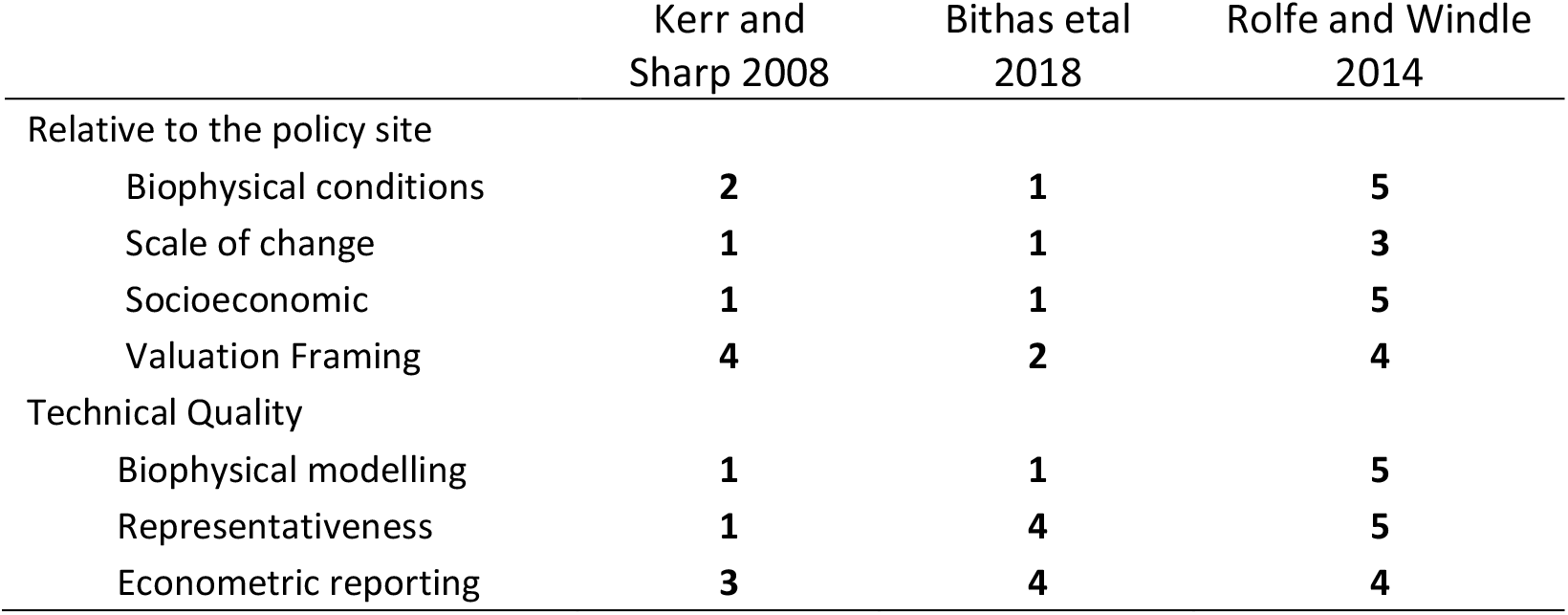
Summary of candidate study suitability

|  | Kerr and Sharp 2008 | Bithas et al 2018 | Rolfe and Windle 2014 |
| --- | --- | --- | --- |
| Relative to the policy site |  |  |  |
| Biophysical conditions | 2 | 1 | 5 |
| Scale of change | 1 | 1 | 3 |
| Socioeconomic | 1 | 1 | 5 |
| Valuation Framing | 4 | 2 | 4 |
| Technical Quality |  |  |  |
| Biophysical modelling | 1 | 1 | 5 |
| Representativeness | 1 | 4 | 5 |
| Econometric reporting | 3 | 4 | 4 |

As the summary suggests, all of the candidates have some deficiencies in the criteria, emphasising the compromise necessary in conducting BT. What does become apparent is that the Rolfe and Windle (2014) study is the preferred candidate for the BT application. Considering the scale of change, the CLIMEX (Figure 3) map suggests that the policy area (scope), and subsequent quantities of goods to be valued (scale), are likely to of be of a much larger magnitude than that considered in any of the candidate listed studies.

#### 3.3 Development of Benefit Transfer Method

Different types of BT can be applied dependent on the extent of available primary study sources. *Function* transfers can facilitate adjustments to values between study and policy sites based in empirical relationships. The function may be estimated from a single study, or from combining values from a set of studies. The list of relevant candidate studies found here indicates that function transfer is not currently possible, and therefore an adjusted unit transfer approach is developed.

The following section describes a BT approach using the WTP estimates from Windle and Rolfe (2014), considered to be the most suitable value estimates found. The DAM generates an estimate of wasp hive density at a spatial scale of a CLIMEX cell. As density increases, so too the negative outcomes to health, amenity and biodiversity. Control options are modelled within the Dam to affect a reduction in wasp hive density, it is this change in hive density that is valued in the BT exercise. To operationalise the Rolfe and Windle (2014) WTP estimates a threshold hive density must be specified so that once the threshold has been reached then management effort will be required to ameliorate impacts. This implies that impacts, although they may occur, are not sufficiently detrimental to warrant WTP to ameliorate them at densities below the threshold.

The BT requires scaling of per unit values by a larger quantity, population and area than was evaluated by the original source study. To treat per unit values as invariant to these changes would be incorrect in light of geographical proximity effects such as distance decay (WTP decreases with increasing distance from the policy area) and diminishing marginal utility (per unit values such as WTP/hectare decrease as quantity increases). Empirical analysis of these effects finds per unit values to be higher in small local case studies than regional or national ones. Therefore, unit values should not be scaled to significantly larger geographic areas or scales without adjustments (Johnston and Duke 2009).

The BT approach comprises four main modules (Figure 6). A Geographic Information System (GIS) forms the basis for the mechanics of conducting the BT within the DAM. Affordability considerations contribute to income and WTP being positively correlated, to account for this marginal WTP estimates are adjusted for differences in income levels between Brisbane and affected areas. The GIS is used to identify the hectares of protected and public areas and number of households effected each year. As the DAM runs over its multi-year time horizon, wasp incursion is modelled to spread into adjacent CLIMEX cells - increasing the quantity of areas and households effected, and hive density increases within cells - also increasing the area and households effected. A distance-decay function is specified that determines how households value benefits as they occur further away. A scale calibration is also applied to adjust marginal WTP relative to increases in the scale of management from that of the source study.

**Figure 6.**
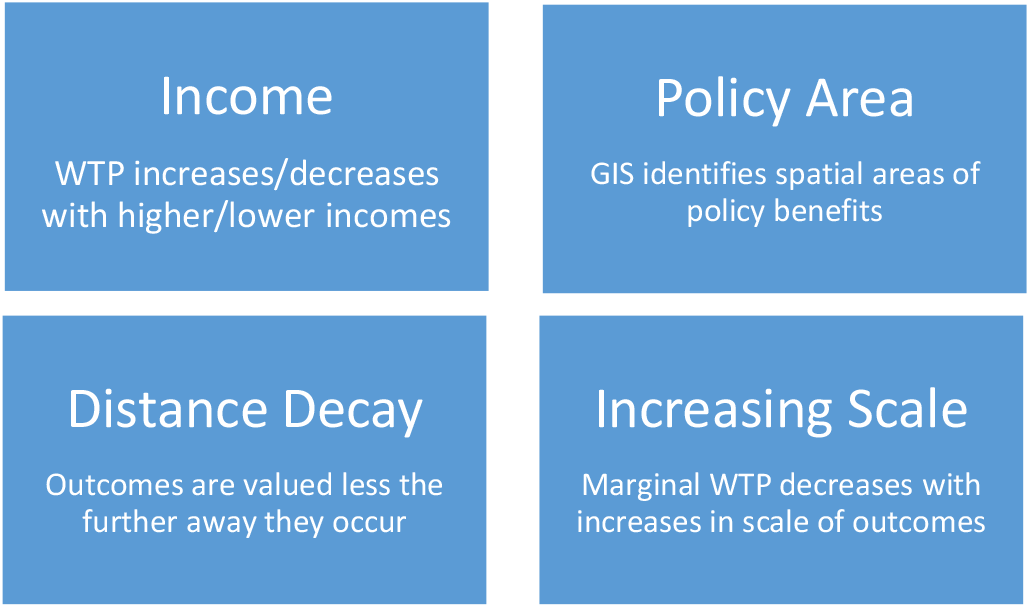
Benefit Transfer method, main modules

##### 3.3.1 Inflation

The first step, when transferring WTP from past studies to the current time period, is an adjustment made to reflect the effect of inflation on general price levels in the economy. The following formula is used to adjust estimates for changes in inflation over the time period between when the study was conducted and when the policy estimates are calculated:

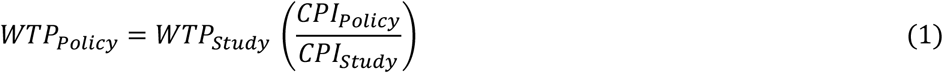

Where:

*WTP_Policy_* = WTP at the policy site
*WTP_Study_* = WTP at the study site
*CPI_Policy_* = The value of the Consumer Price Index for the year of the policy site estimates^3^
*CPI_Study_* = The value of the Consumer Price Index for the year of the study site estimates

Applying equation 1 to Rolfe Windle (2014) increases WTP values by approx. 25% (Table 8).

**Table 8.** Inflation adjusted WTP for Rolfe and Windle (2014)

| Attribute | WTP in 2009 | WTP in 2020 |
| --- | --- | --- |
| Households | \$0.13/1,000 houses | \$0.16 |
| Public areas | \$172/100 ha | \$215 |
| Protected areas | \$0.08/100 ha | \$0.10 |

##### 3.3.2 Income

The next step is to adjust for differences in income levels between Brisbane households and households in policy target areas. Demand for goods and services generally increases in line with income and accounting for differences in income levels between study and policy sites can be an important consideration to improve the accuracy of estimates. Unit values are adjusted using equation 2. With this adjustment, WTP will increase at the policy site if incomes of beneficiaries are higher relative to those of Brisbane households. The income elasticity of WTP conditions whether WTP increases in the same proportion as rises in income, less than proportionally, or greater than proportionally. We set this elasticity equal to one, meaning that WTP is a constant share of income.

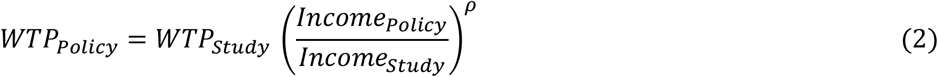

Where:

*WTP_Policy_* = WTP at the policy site
*WTP_Study_* = WTP at the study site
*Income_Policy_* = Income of beneficiaries at the policy site
*Income_Study_* = Income of beneficiaries at the study site
*Ρ*= Income elasticity of WTP

The income adjustment is operationalised spatially using median household income at SA1 level^4^ aggregated up to the CLIMEX resolution based on the proportion of SA1 shape intersecting with a CLIMEX cell.

**Figure 7.**
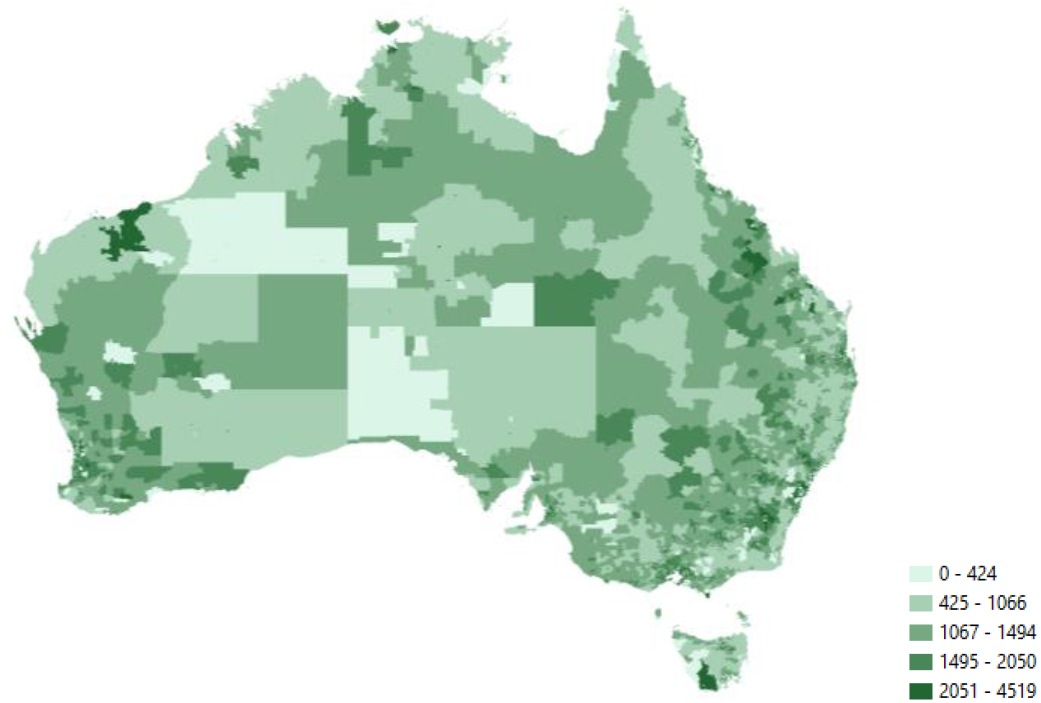
Median weekly household income at SA1 level

To illustrate, we use median weekly household income for Australia ($1,438) and Brisbane ($1,562) to adjust estimates in (Table 9). The ratio is less than one and so values are adjusted downward (Table 9).

**Table 9.** Income and Inflation adjusted WTP for Rolfe and Windle

| Attribute | WTP |
| --- | --- |
| Households | \$0.15/1,000house |
| Public areas | \$198/100ha |
| Protected areas | \$0.09/100ha |

##### 3.3.3 Policy Area

To define the policy area to be valued, we use the Land Use of Australia 2010-11 data^5^ (Figure 8). From the list of classes (Table 10) we form a spatial concordance with the definitions of affected public areas, and protected bushland (Figure 7). The number of households affected within each CLIMEX cell is derived using SA1 level data (Figure 7). As mentioned above, this may change at each time step of the DAM. Excluding Western Australia, there are 522 CLIMEX cells with suitable habitat (Figure 8).

**Figure 8.**
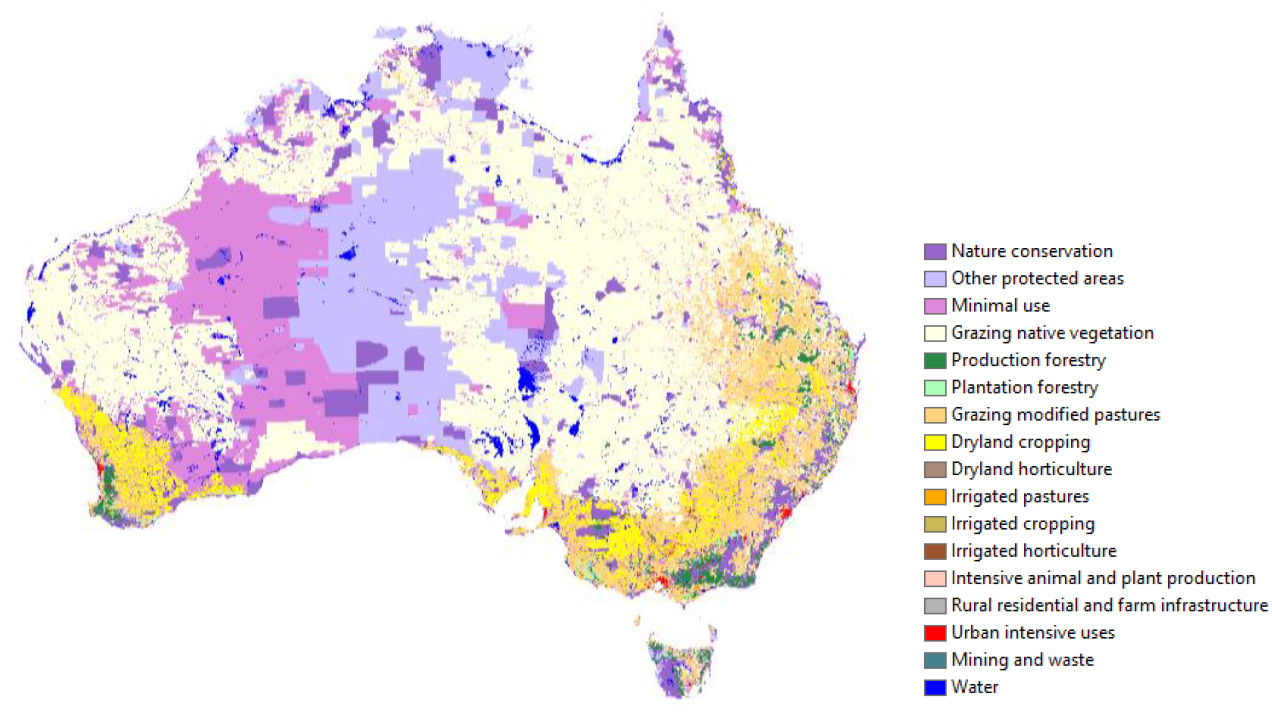
Land Use of Australia 2010-11

**Table 10.**
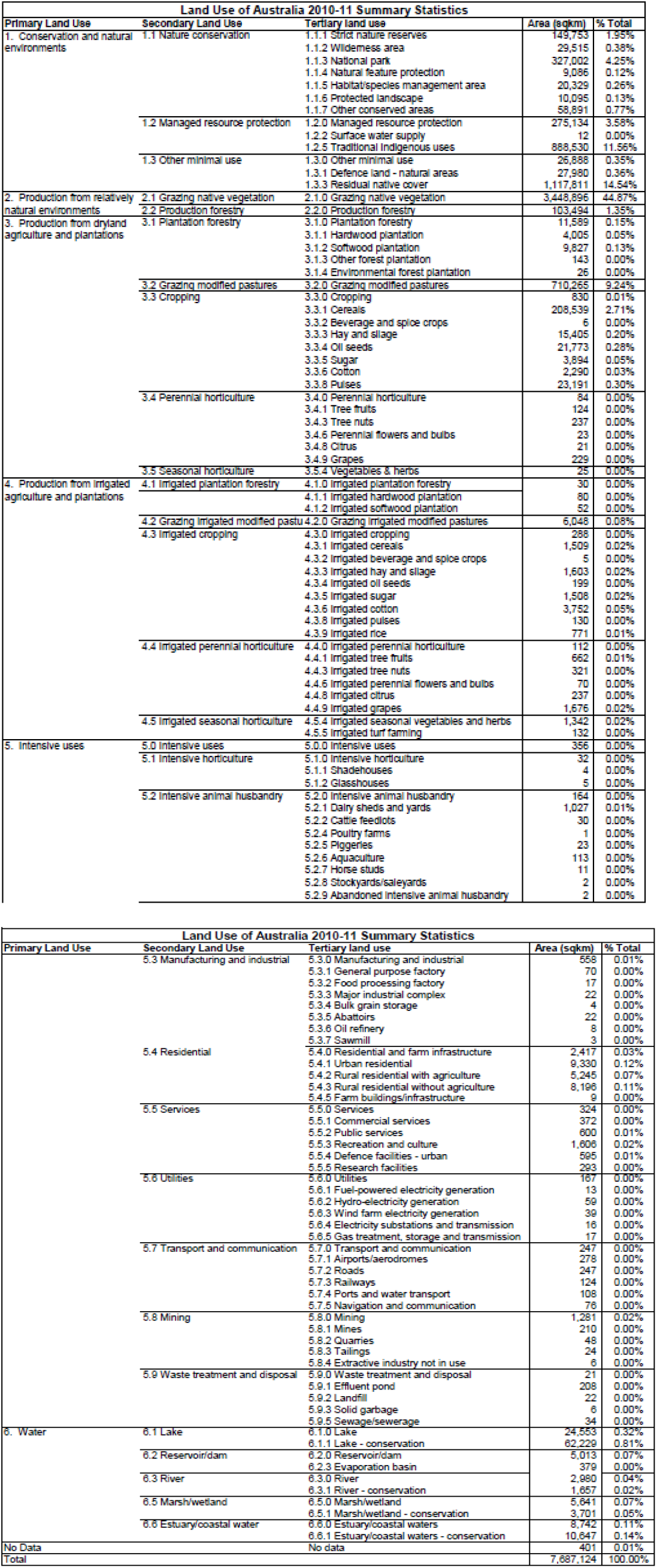
Land Use of Australia 2010-11 classes

**Figure 9.**
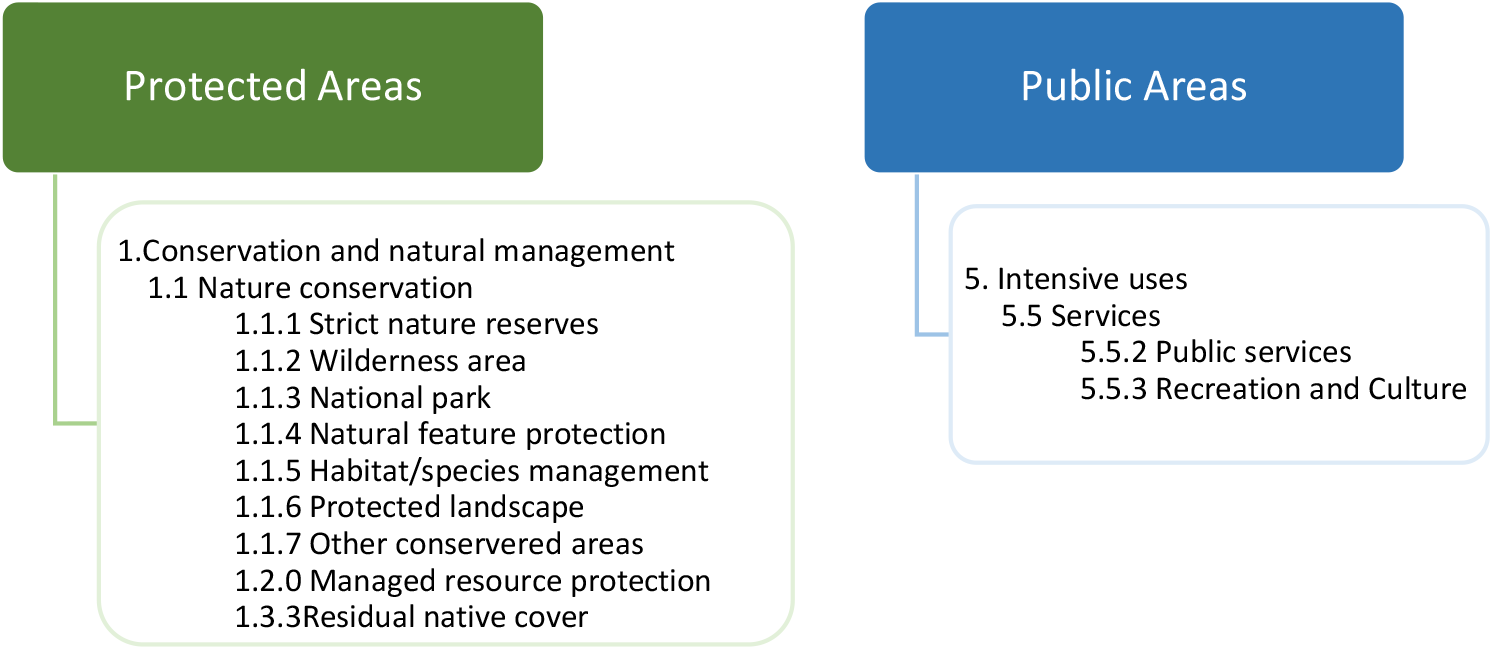
Spatial concordance of LUA and Windle and Rolfe RIFA management attributes

The number of homes protected is calculated using SA1 level Census data (Figure 10).

**Figure 10.**
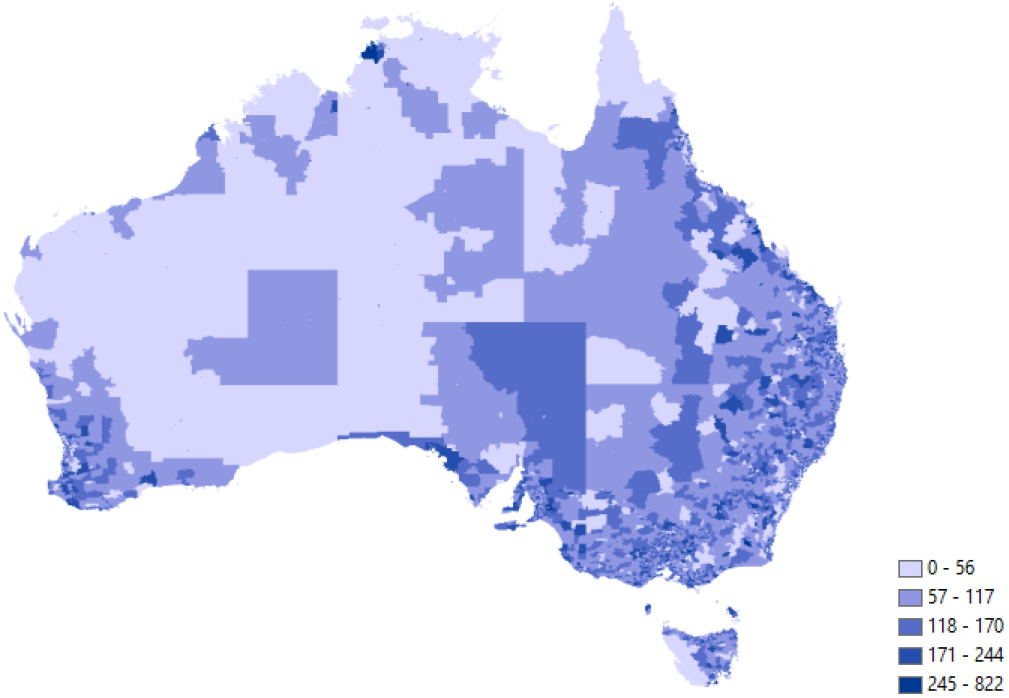
Number of households at SA1 level

Calculating the number hectares of protected bushland, public areas, and number of households for each CLIMEX cell reveals that most of the nations protected bushland is located in areas unsuitable as wasp habitat (Figure 3). Conversely, over 80% of public areas and households are located in suitable habitats.

**Figure 11.**
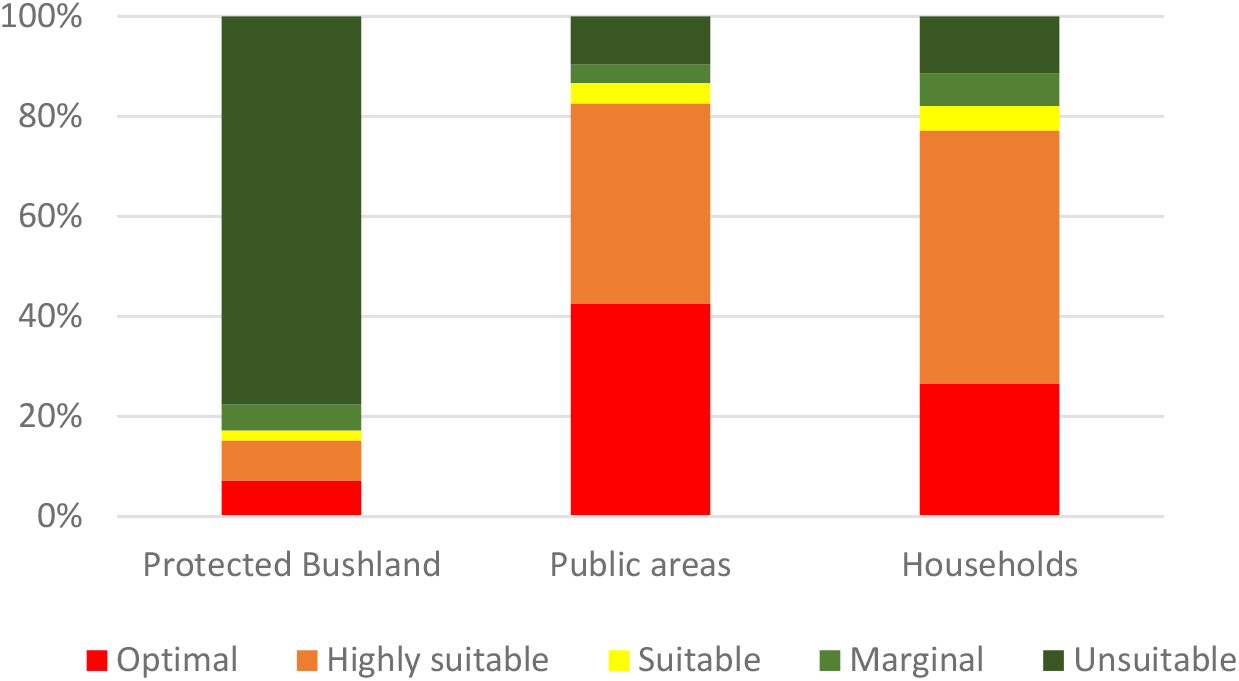
Areas of protected bushland, public areas, and households by CLIMEX suitability

##### 3.3.4 Distance Decay

As we move further and further away from an affected area, households WTP for benefits of managing that area tend to fall. The cost of accessing more distant areas increases in time commitment and expenditure, and for many environmental goods the availability of substitutes increases with distance from a site, contributing to distance-decay effects. Therefore, if we were to naively aggregate marginal WTP from the original source study ($X in Figure 12) that was estimated based on an assumed distance of A between households and the policy site(s), over larger distances we would likely overestimate the aggregate value of benefits.

Accounting for distance-decay effects can also be important in establishing the applicable geographical delineation of the population of beneficiaries (Jorgenson et al. 2013). One of the central considerations in the BT exercise is determining the population whose benefits should be counted. Using distance decay provides a solution to this problem in defining the distance from an affected area at which WTP falls to zero ($0 in Figure 12). This area is called the economics jurdisction, and tells us who is ‘in the market’ for the benefits of management outcomes at a site. As an alternative, the political jurisdiction is also commonly used to define the number of households WTP for policy benefits. Usually because this describes the catchment of those voting for, and/or funding the policy.

**Figure 12.**
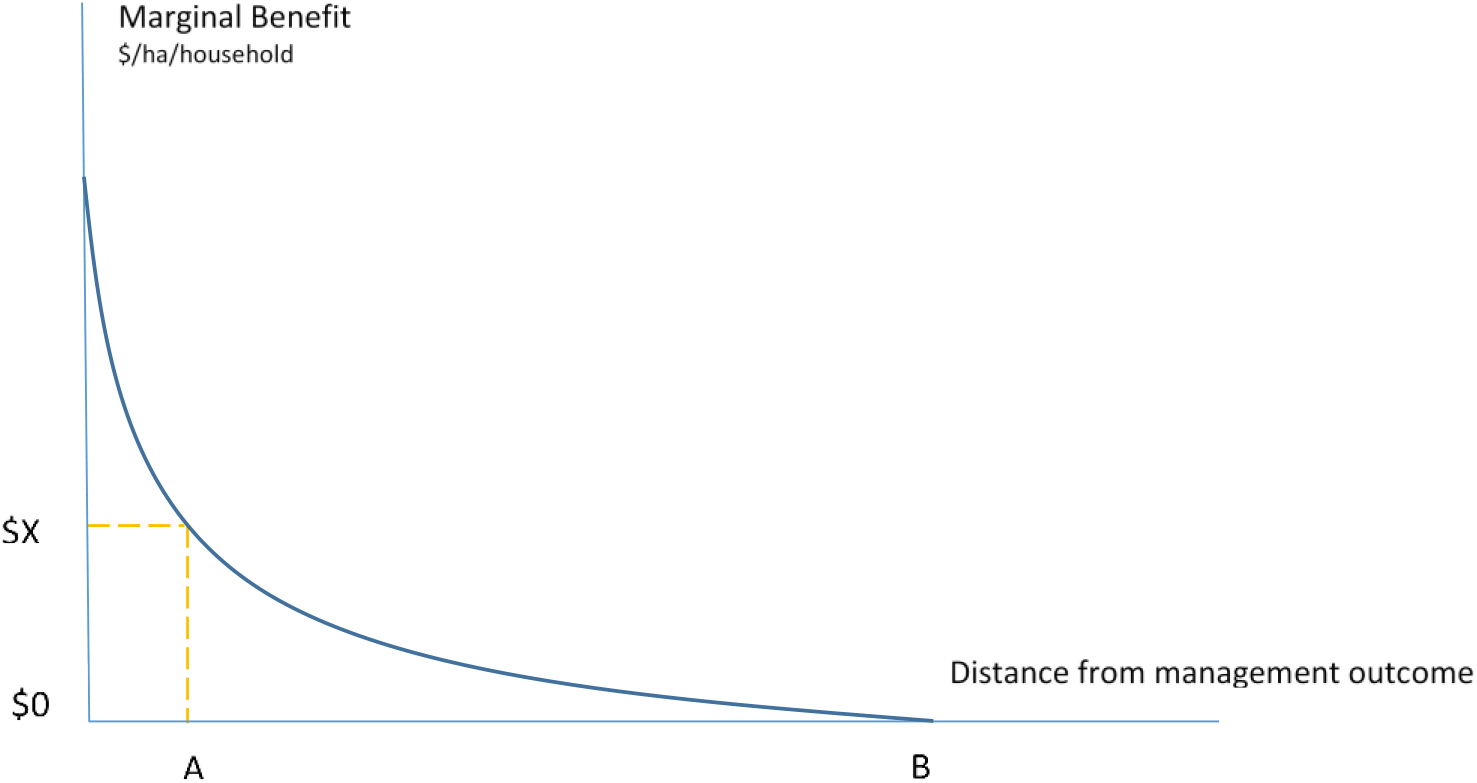
Distance-decay of marginal benefits

To implement a distance-decay relationship we need to specify the rate at which WTP falls with increasing distance from a management area. We also need to consider whether a distance-decay relationship is the same for each benefit valued. For example, we might expect that protection of homes is characterised by more private than public good benefits compared with managing areas of protected bushland or protection of public recreation and school areas. While there is a network effect in control of homes as those not managed can provide habitat for re-infestation, a majority of the benefits are likely consumed by each individual homeowner. Whereas, recreational areas and protected bushland generate benefits for a larger number of people that are more public good in nature. The implication is that a distance-decay function for reducing the impact on private homes is steeper than one relating to protected bushland or public areas.

Examining studies that estimate distance-decay functions demonstrates that the relationship varies mainly in line with the significance of the affected area. Protecting iconic assets, for example, are typically associated with a shallower decline and WTP may not reach zero before the political jurisdiction is reached. While, for example, improvements in degraded less significant areas are valued predominantly by those close by, who are aware of them and can enjoy improvement benefits.

In an Australian CE estimating WTP for protection of Great Barrier Reef, Rolfe and Windle (2012) find that WTP declined by about 50% from the local population to out-of-state populations, and then remained relatively constant at further distances. They suggest the possibility that WTP may increase at the furthest distances considered. This is plausible given the iconic status of this asset, and a lack of realistic substitutes. Estimates of WTP for protecting the Norfolk Broads in England declined from a mean value of £39 at a distance of 20 km, to £14 at a distance of 110–150 km away from the Broads area (Bateman et al. 2000). Moran (1999) use these estimates to calculate a critical distance of 214 km at which WTP equals zero. In another study of WTP for preservation of a National Park in England, values were found to diminish with distance very slowly, to the extent that WTP did not fall to zero by the time political and economic jurisdictions coincided (Bateman et al. 2006).

The functional form of the decay function can also be an important consideration. In a study estimating WTP for improvements in low-flow conditions on the river Mimram in England (Hanley 2003), distance enters models in logarithmic form, meaning that WTP will never reach zero, irrespective of the distance from the policy site a household is. To calculate aggregate benefits, the authors specify a 100km radius as the relevant economic jurisdiction. They find that households within 0.5km were WTP approx. £9/year, which dropped down to approx. £1.8/year at 20km. Relatively steep functions include a Scottish study (Hanley et al. 2001) finding WTP for protecting a heather moorland site fell to zero at a 40km radius, and WTP for protection of a rough grassland site at 50km. Georgiou et al. (2000) estimate WTP to clean up the River Tame in Birmingham is equal to zero at a 25km radius. Values for improvements of the River Thame were also examined by Bateman et al. (2006) who find consistent results in estimating an economic jurisdiction at a 20km radius to the improvement site.

Other functional forms of the decay function include quadratic and logarithmic, the former implies that WTP decreases at an increasing rate the further away a household is located, while the former has a more pronounced decrease in WTP initially that flattens out at greater distances. The selection of appropriate form is typically determined by optimising statistical model properties (Schaafsma et al. 2013).

To accommodate distance-decay effects we specify weighting functions for WTP that reduce WTP for management outcomes further from a household’s location. We specify separate weighting functions for each of the three management types: effected homes, public areas, and protected bushland. Where the economic jurisdiction is assumed to be smallest for effected homes, slightly larger for effected public areas, and largest for benefits of management of protected bushland. The original source study did not consider the distance between a household and the nearest managed area. This may reflect the non-specific framing of managed area location, which could occur at any applicable area within the Greater Brisbane jurisdiction. For the purposes of this BT we assume that distance-decay is not present within the Greater Brisbane area. And approximate this area as being one CLIMEX cell. Accordingly, distance-decay equations assume that household’s value benefits outside of a household’s own CLIMEX cell at the same level as the original study, the benefits outside of a household’s own cell are then subject to distance-decay effects and downward adjustment.

To illustrate the adjustment, consider equation 3 describing the relationship for areas of protected bushland:

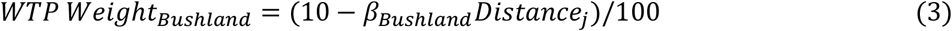

Where:

*β_Bushland_* = 0.5
*Distance_i_* = *Km from Centroid of Cell*_0_

This specifies that managed protected bushland is valued by a household if it occurs within a 200km radius.

Figure 13 depicts this relationship by showing the bands of valuation decay from the households‘ location. Within the households own CLIMEX cell, *Cell_0_*, the distance is zero and therefore no weight is applied. At Cell_1_ the distance is 50km (each CLIMEX cell is a 50km by 50km square, the distance from centroid to centroid for adjacent cells is 50km) generating a weighting of 0.75 (Table 11).

**Figure 13.**
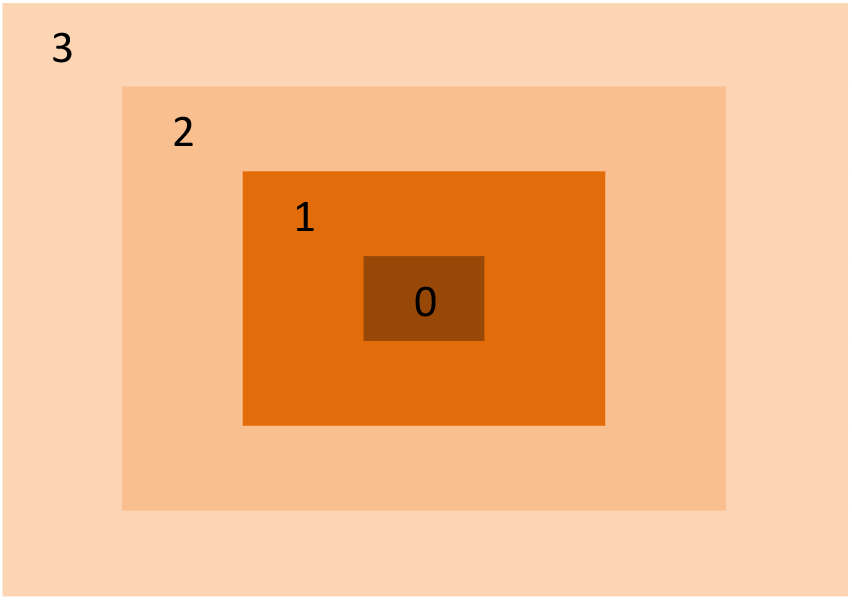
Depiction of distance-decay relationship for valuing protected bushland

The choice of beta in equation 3 dictates the rate of decay and therefore the economic jurisdiction in which management changes are valued by households located in *Cell_o_*. For a beta equal to 0.5 the total area encompasses 49 CLIMEX cells (7×7) which is 122,500km^2^ with a 350km diameter. The main implication of this choice is that management benefits outside of this jurisdiction are not valued by households in *Cell_o._* To illustrate this scheme, we apply the inflation and income adjusted unit value/ha (Table 11), and assume that all contiguous cells contain the average number of hectares of protected bushland per CLIMEX cell (38,000ha), and that management benefits are achieved in all of these cells. Using these figures generates a total WTP for management benefits inside the economic jurisdiction of $734.

**Table 11.** Calculating distance-decay effect for protected bushland valuation

| CLIMEX Cell | Centroid Distance | Weight | Number of Cells in band | WTP with Distance-decay |
| --- | --- | --- | --- | --- |
| 0 | 0 | 1 | 1 | \$35 |
| 1 | 50 | 0.75 | 8 | \$209 |
| 2 | 100 | 0.50 | 16 | \$279 |
| 3 | 150 | 0.25 | 24 | \$209 |
| 4 | 200 | 0 |  |  |

##### 3.3.5 Increasing scale

Failure to account for nonlinearity of marginal WTP with policy scale can be a source of significant error (Schaafsma 2015). The second important adjustment to unit WTP values concerns the effect on WTP of increases in the scale of management outcomes relative that which was conveyed in the original study. Diminishing marginal utility reflects standard economic theory for a normal good, the marginal benefit of the next unit of a good consumed is less than the previous and so demand is downward sloping and prices are lower at higher quantities (Figure 14). Evidence from environmental valuation supports this understanding in finding that per unit values tend to be higher in small local case studies than regional or national ones (Rolfe et al. 2011; Rolfe and Windle 2008; van Bueren and Bennett 2004). And that marginal WTP for an additional hectare of protected area of which supply is already high should be lower than for an additional hectare of a scarce resource (Schaafsma 2015). The practical implication for BT application is that if we naively transferred the source studies marginal WTP, $X in Figure 14, beyond the scale assumed in that study (A) to larger scales we risk generating upwardly exaggerated estimates of value.

**Figure 14.**
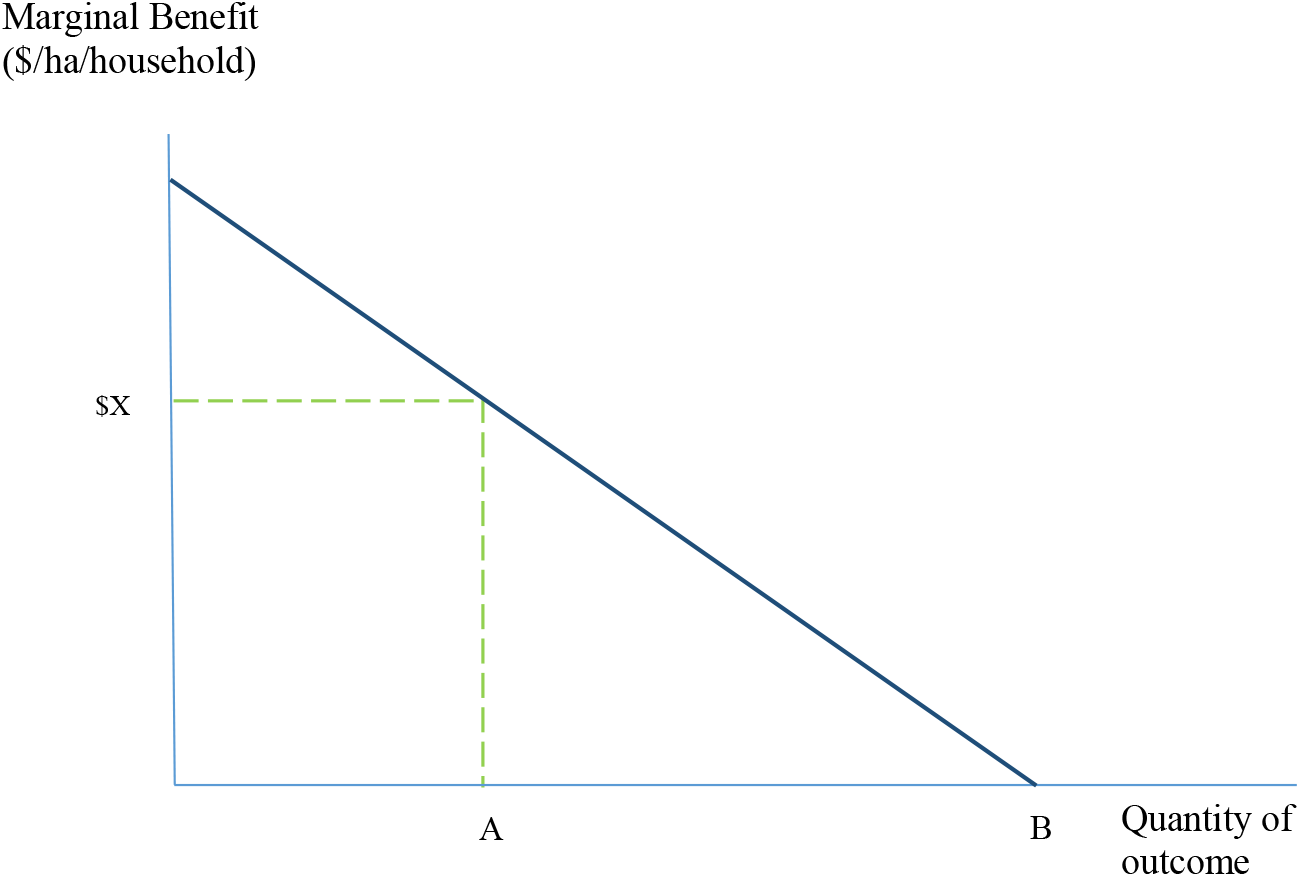
Diminishing marginal benefits with increasing scale

To operationalise an adjustment for unit values incorporating scale effects we apply a method that calibrates the original source study values by the ratio of the original source study scale with that of new policy area (Rolfe et al. 2013). The intuition is that as scale goes above that of the original study, marginal WTP is lowered, conversely if the new policy scale is smaller than that of the original study then marginal WTP will increase (equation 4).

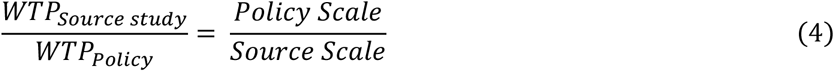

To illustrate we continue the example of valuing protected bushland areas. The scale implied for the original source study is 243,000 ha, the 49 CLIMEX cells within a household’s economic jurisdiction generate a total policy scale of 1,862,000ha. This generates a ratio of 0.13 with which we can calibrate (downward) the WTP estimates in Table 12. We apply this calibrated WTP to benefits generated outside of the households own CLIMEX cell, and assume that benefits realised within a h u h d’ own cell are valued using the inflation and income adjusted source WTP estimates. The practical implication is that areas of bushland outside a household’s own CLIMEX cell are valued at $0.012/100ha ($0.09*0.13). We can see that this reduces value estimates significantly from those omitting a scale adjustment (Table 12).

**Table 12.**
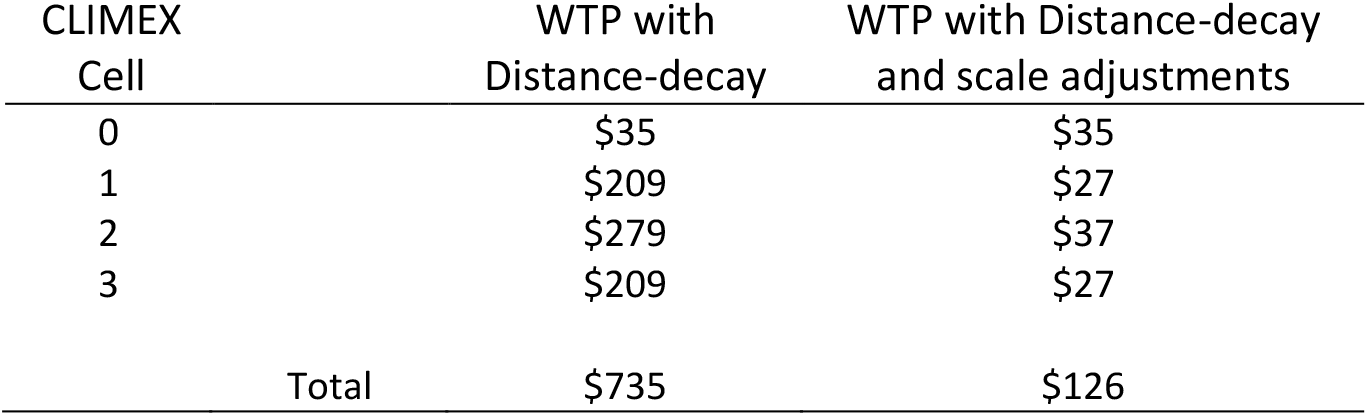
Calibrating WTP for scale effect for protected bushland valuation

| CLIMEX<br>Cell | WTP with<br>Distance-decay | WTP with Distance-decay<br>and scale adjustments |
| --- | --- | --- |
| 0 | \$35 | \$35 |
| 1 | \$209 | \$27 |
| 2 | \$279 | \$37 |
| 3 | \$209 | \$27 |
| Total | \$735 | \$126 |

The method developed thus far for aggregation of marginal WTP values up to the population level considers the spatial jurisdiction of beneficiaries for wasp control outcomes. A further dimension that warrants incorporation into the aggregation process is an estimation of the proportion of households within a jurisdiction that do not have positive WTP for the control outcomes. Relying on simple multiplication of the number of households within a jurisdiction by marginal WTP values while ignoring this issue potentially introduces upward bias into the population level aggregate value estimate (Morrison 2000). To apply this adjustment, response rates to Stated Preference surveys have been used as a measure of the proportion of households with zero WTP (Mitchell and Carson 1989). The assumption is that for most non-respondents the marginal benefits of completing a questionnaire are lower than the marginal costs (Dillman et al. 2014), reflecting that their value of the good of interest is low and therefore their WTP is negligible (Bennett et al. 1997). Complications in applying response rates as proxy measures arise in three important ways. First, the assumption that all survey non-respondents have zero WTP may not be accurate when some factors explaining non-response may imply positive WTP for some non-respondents, including that participants were simply too busy to be able to complete a survey in the given time (Morrison 2000). Secondly, the ability to observe response rates across survey modes varies, and is particularly difficult for internet panel-based approaches. Thirdly, even when rates can be reliably estimated, the applied CE literature demonstrates that rates can vary by survey mode independently of the good being assessed, for example internet modes typically achieve lower rates than interview or mail-and-return modes.

Ideally the survey response rate from the source study (i.e. Rolfe and Windle 2015) would be used here, however as an internet panel survey mode was used the response rate was not directly observable. Reported response rates to CE surveys in Australia for environmental resources include: 44% from a drop-off and pick-up mode study valuing wetland management in New South Wales (Morrison 2000); 38% from a sample drawn from telephone numbers subsequently completed online valuing native biodiversity habitat management in South Australian (MacDonald and Morrison 2010); 85% for the mail-and-return version and between 5% and 44% for internet panel versions of a study valuing protection of the Great Barrier Reef to Australian households (Rolfe and Windle 2012). Considering these response rates overall, and assuming that 30% of non-respondents have WTP equal to the sample mean (Morrison 2000) suggests that for sensitivity analysis a reasonable approximation of the proportional of households with positive WTP is in the range of 50% and 60%.

##### 3.3.6 Discounting future values

The decision analysis model calculates values for each year into the future, those future values must be discounted back to the present day to allow for comparison with current costs and benefits. The following formula can be used to discount future values:

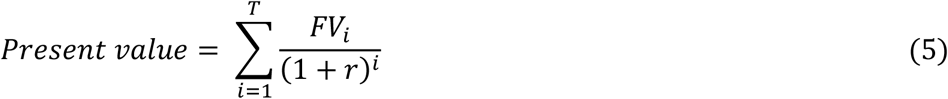

Where: *FV_i_* = value estimate in year *I; r* is the discount rate, and *i*= number years into the future when the value is realised. The Australian Government’ base social discount rate is 8%, recommending sensitivity analysis using 3% and 10% (Harrison 2010)^6^.

## Appendix B: Distribution of impacts and benefit by jurisdiction

**Table B1.**
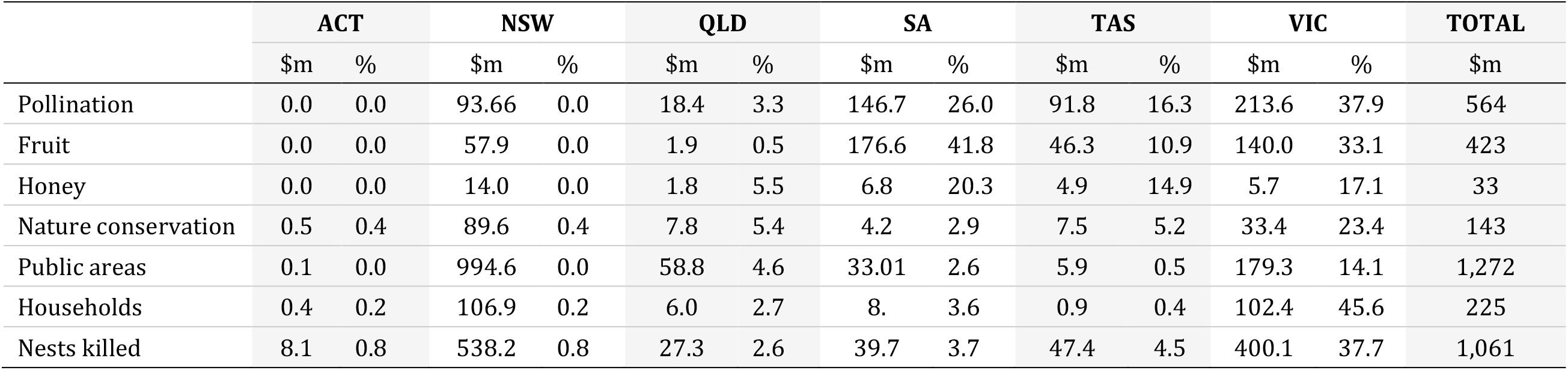
The distribution of wasp damages from the ‘no-control’ scenario.

|  | ACT |  | NSW |  | QLD |  | SA |  | TAS |  | VIC |  | TOTAL |
| --- | --- | --- | --- | --- | --- | --- | --- | --- | --- | --- | --- | --- | --- |
| | \$m | % | \$m | % | \$m | % | \$m | % | \$m | % | \$m | % | \$m |
| Pollination | 0.0 | 0.0 | 93.66 | 0.0 | 18.4 | 3.3 | 146.7 | 26.0 | 91.8 | 16.3 | 213.6 | 37.9 | 564 |
| Fruit | 0.0 | 0.0 | 57.9 | 0.0 | 1.9 | 0.5 | 176.6 | 41.8 | 46.3 | 10.9 | 140.0 | 33.1 | 423 |
| Honey | 0.0 | 0.0 | 14.0 | 0.0 | 1.8 | 5.5 | 6.8 | 20.3 | 4.9 | 14.9 | 5.7 | 17.1 | 33 |
| Nature conservation | 0.5 | 0.4 | 89.6 | 0.4 | 7.8 | 5.4 | 4.2 | 2.9 | 7.5 | 5.2 | 33.4 | 23.4 | 143 |
| Public areas | 0.1 | 0.0 | 994.6 | 0.0 | 58.8 | 4.6 | 33.01 | 2.6 | 5.9 | 0.5 | 179.3 | 14.1 | 1,272 |
| Households | 0.4 | 0.2 | 106.9 | 0.2 | 6.0 | 2.7 | 8. | 3.6 | 0.9 | 0.4 | 102.4 | 45.6 | 225 |
| Nests killed | 8.1 | 0.8 | 538.2 | 0.8 | 27.3 | 2.6 | 39.7 | 3.7 | 47.4 | 4.5 | 400.1 | 37.7 | 1,061 |

**Table B2.**
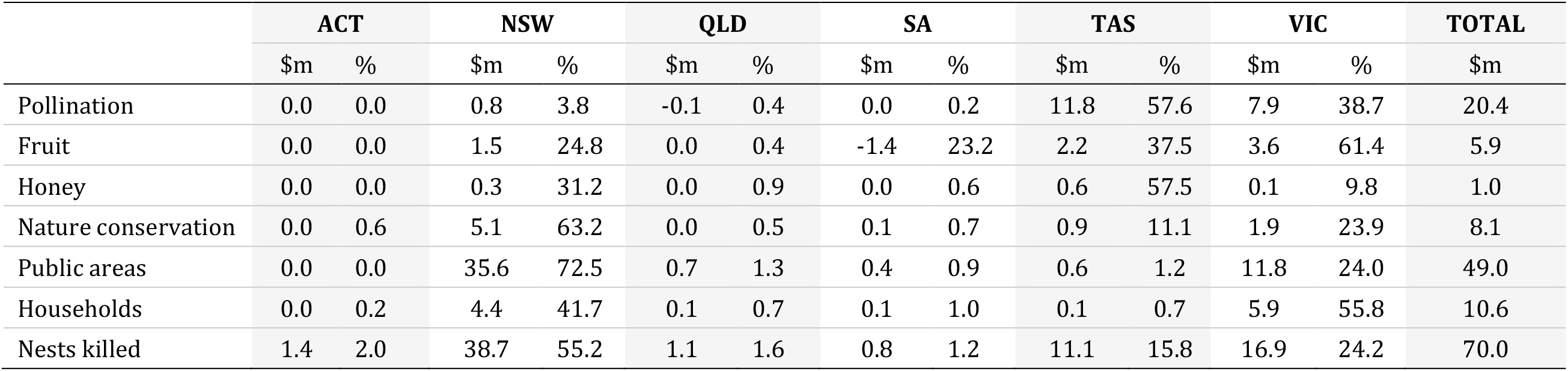
The distribution of benefits from biocontrol under scenario HH.

## Appendix C: Detailed results by state using Scenarios LL and HH

The CDFs presented in this Appendix show the range of values for benefits, including negative values, for each state by industry.

### A.1. Scenario LL

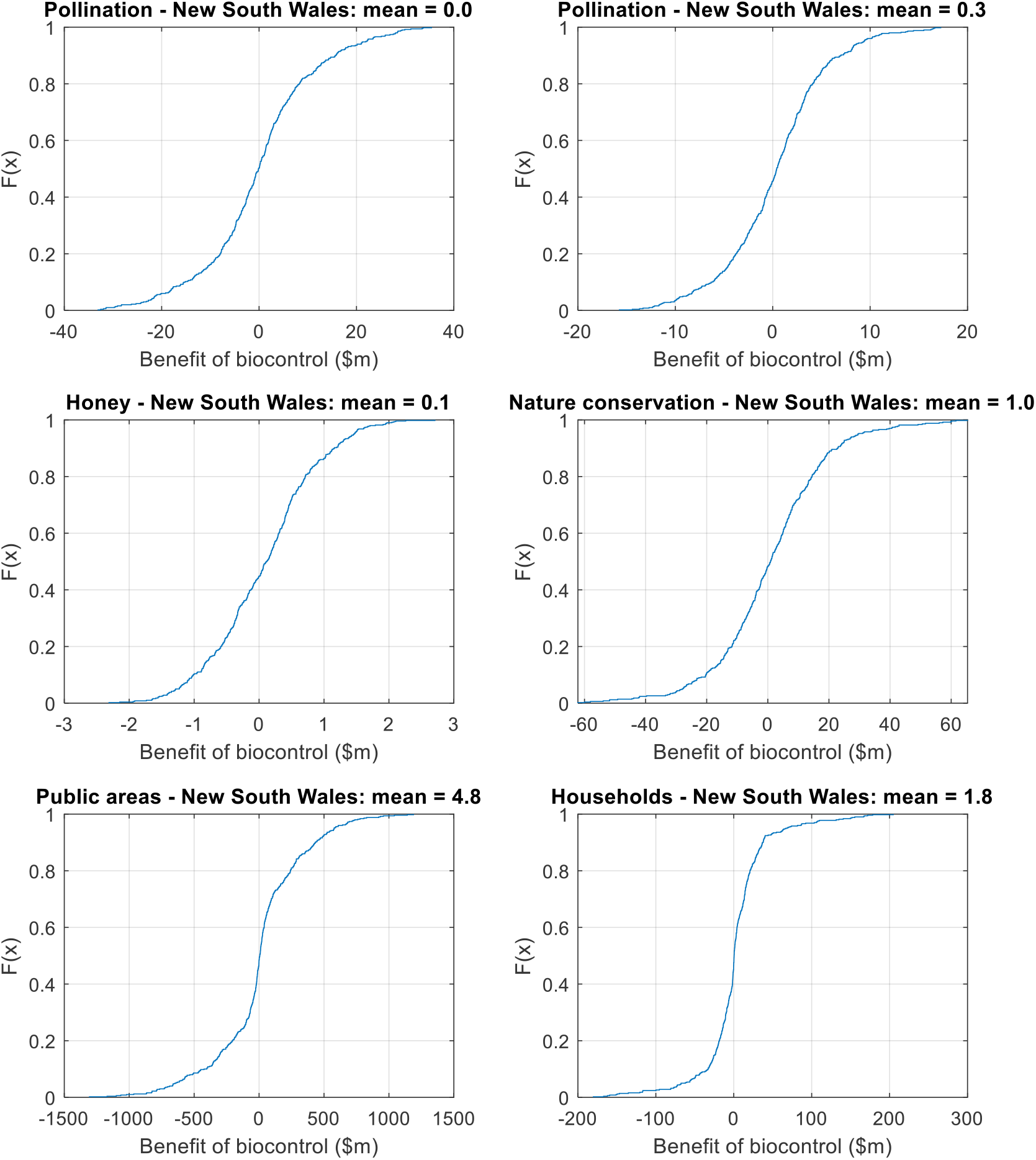
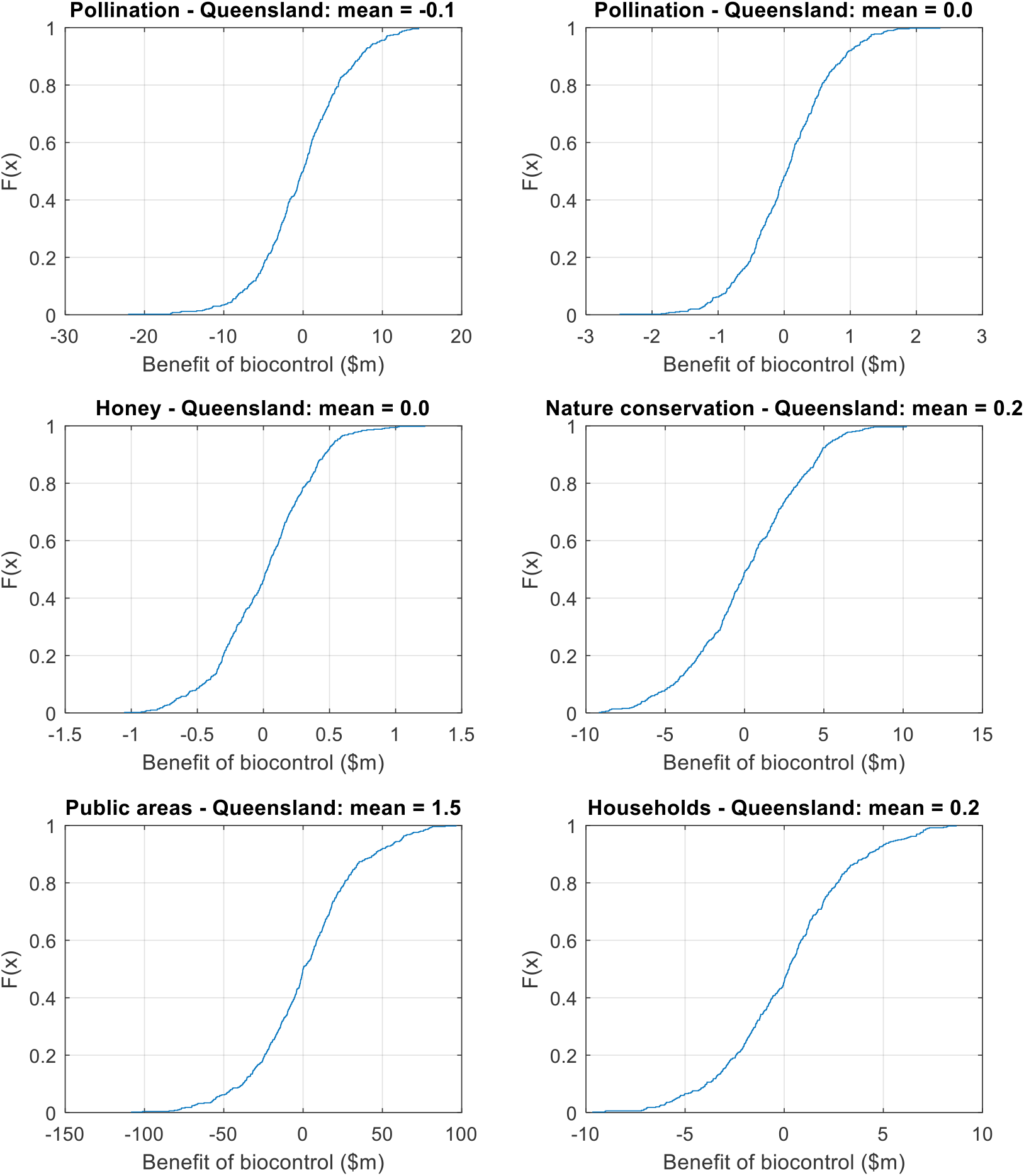
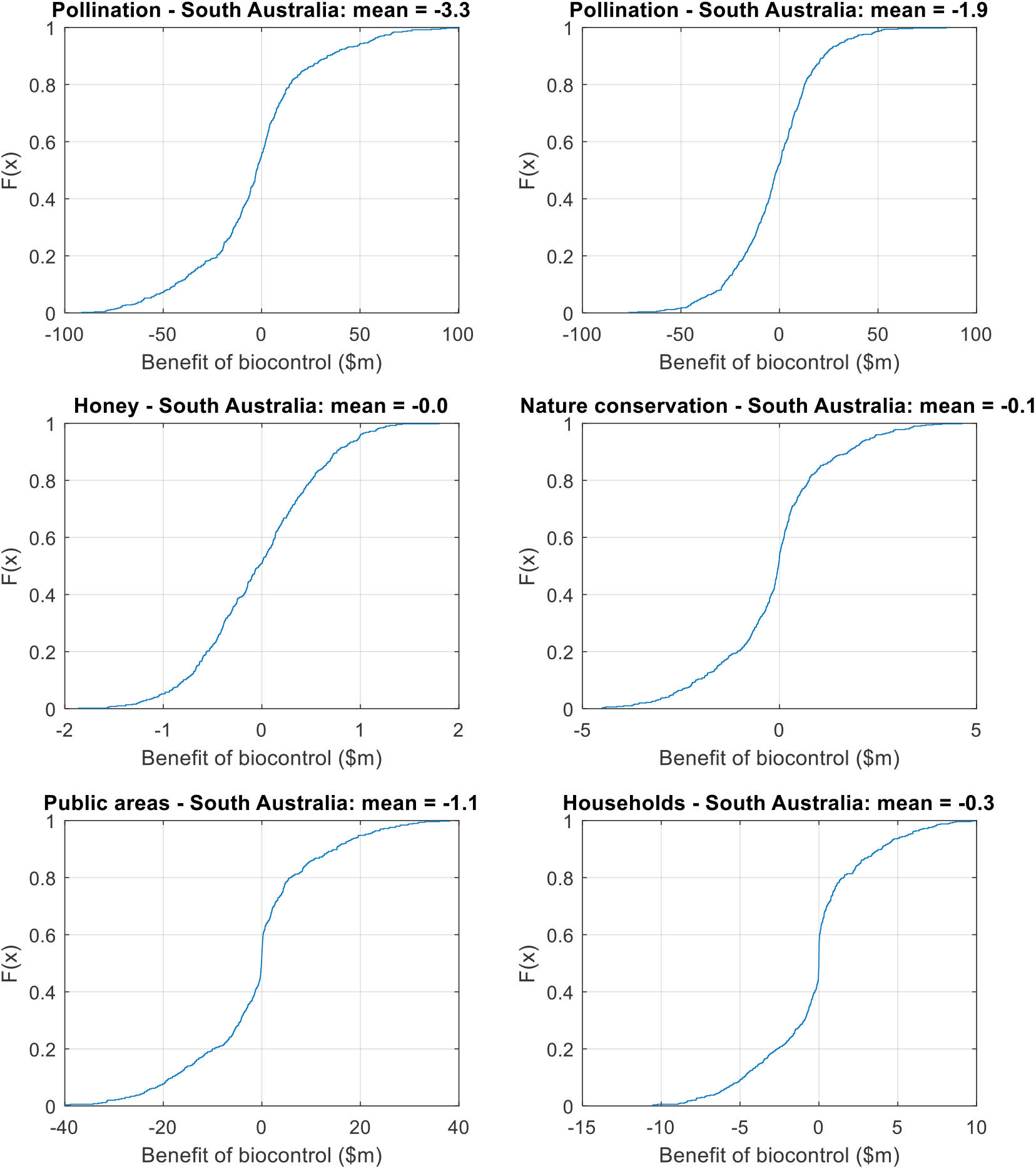
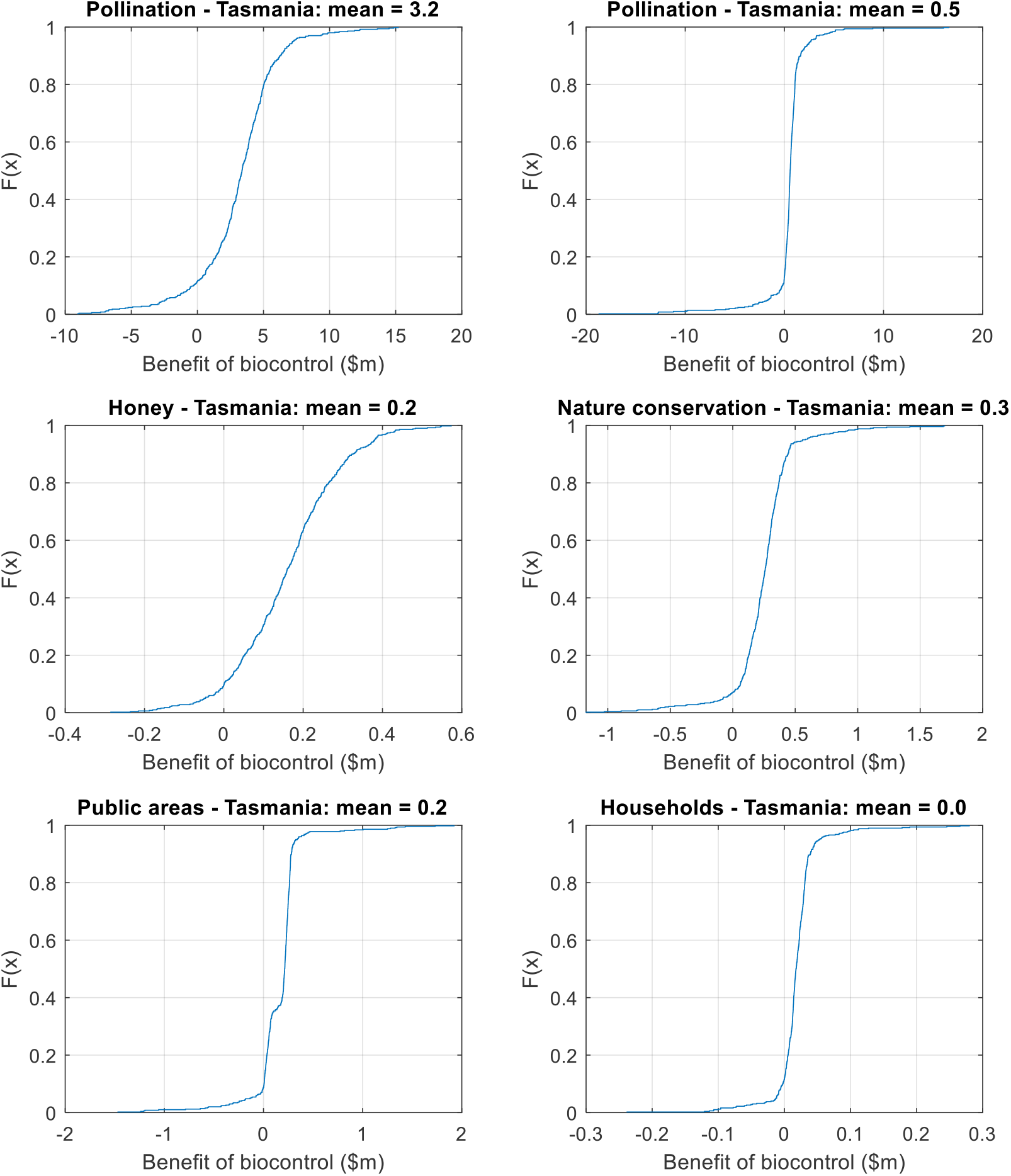
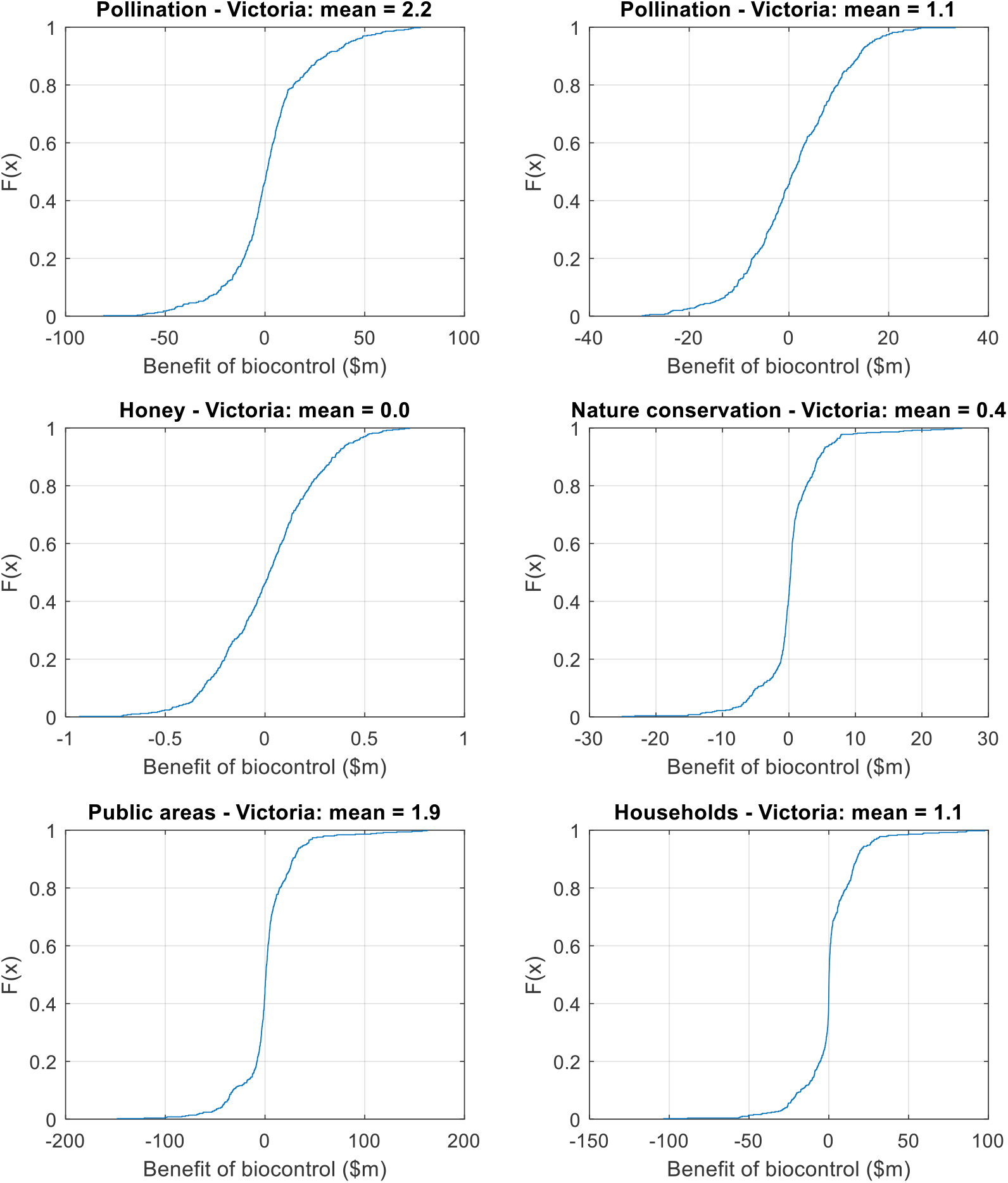

### A.2. Scenario HH

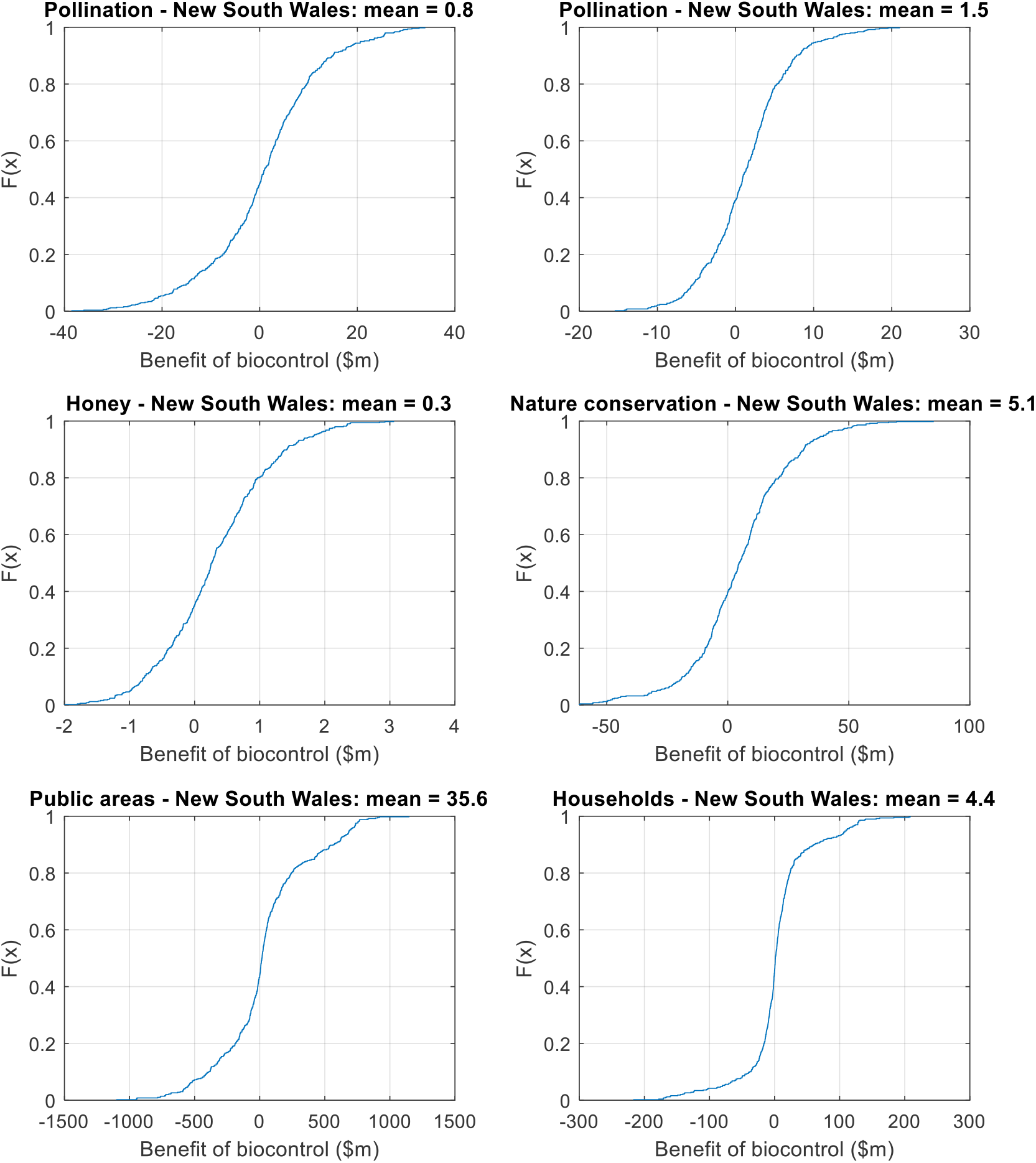
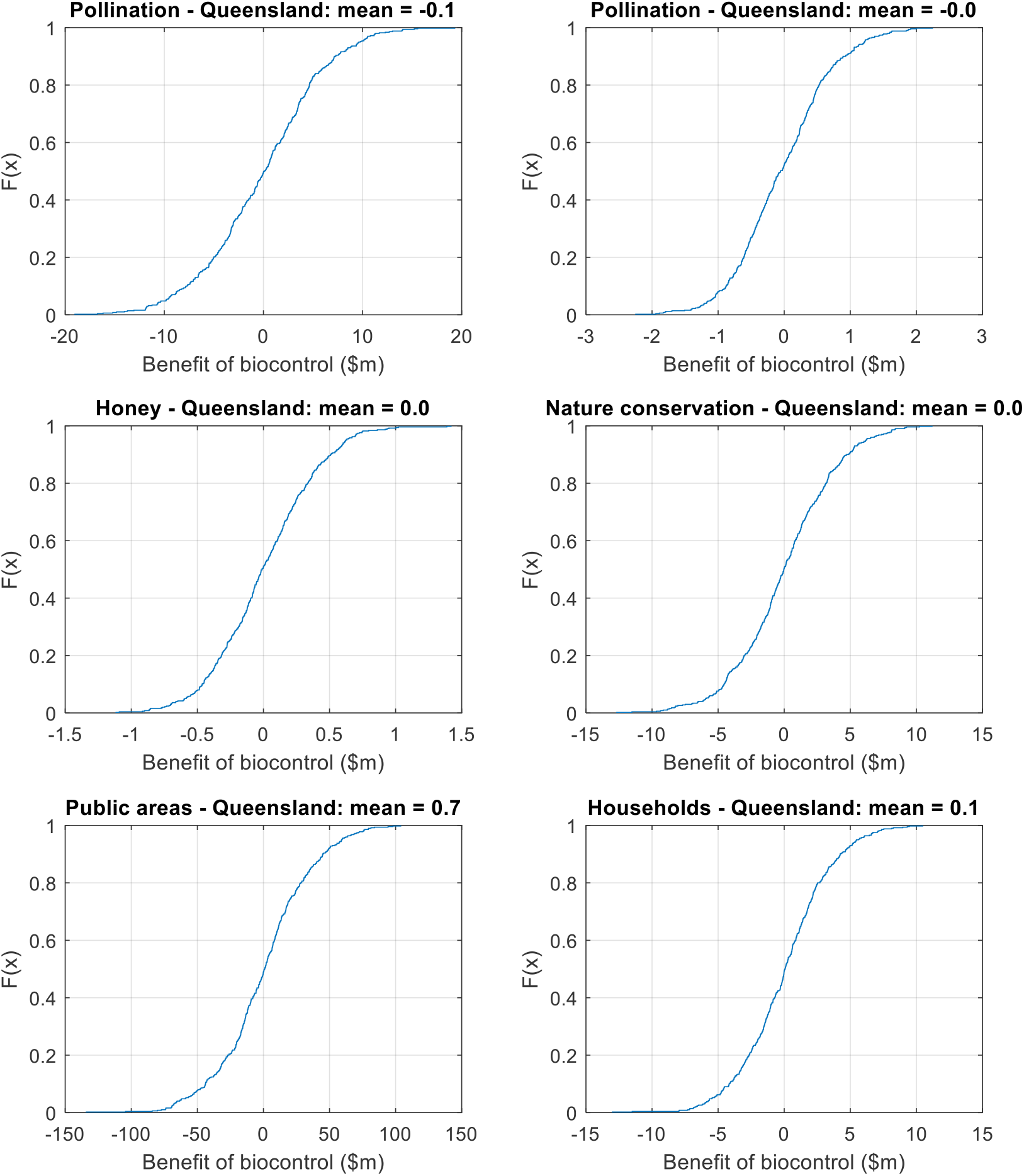
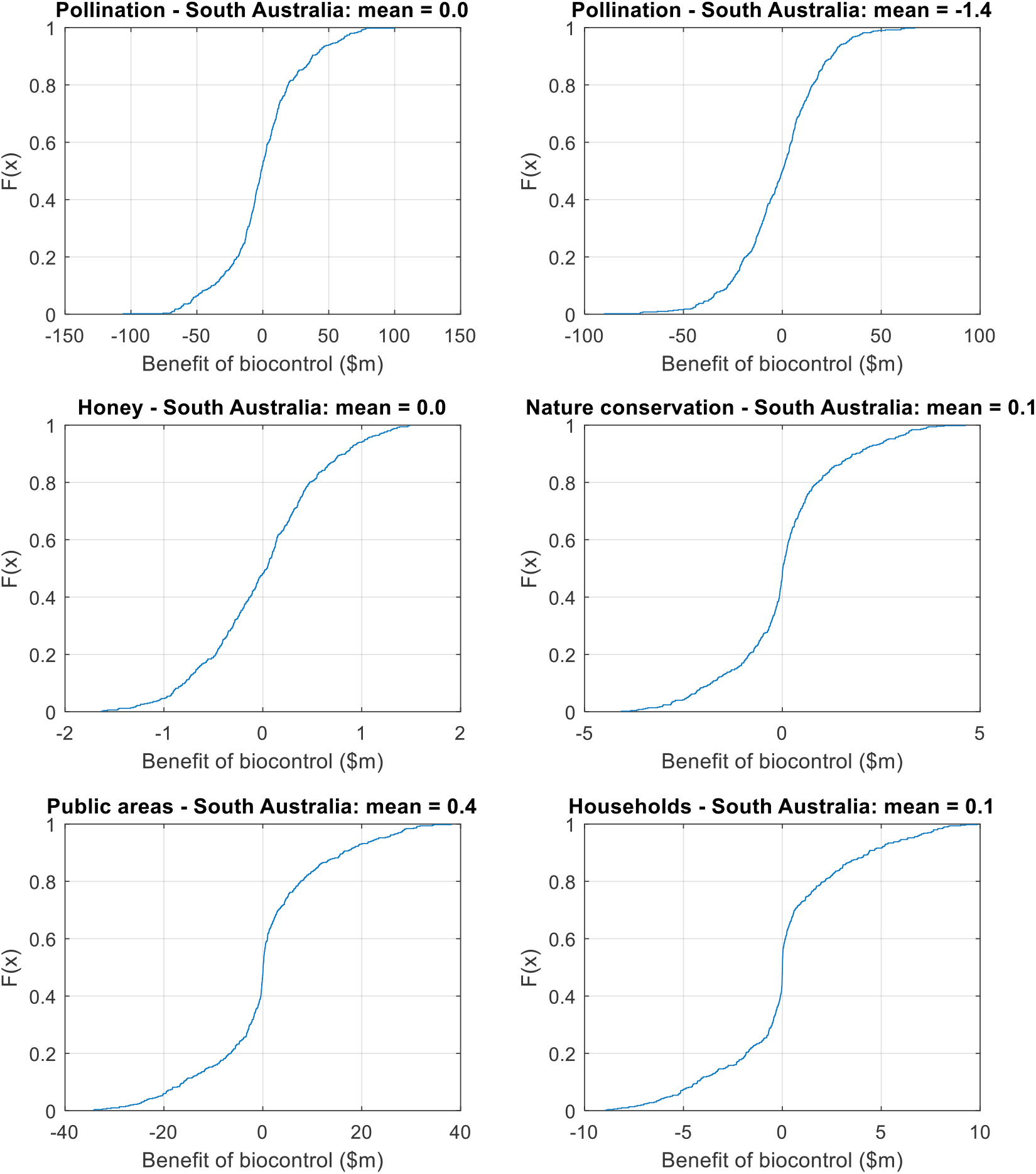
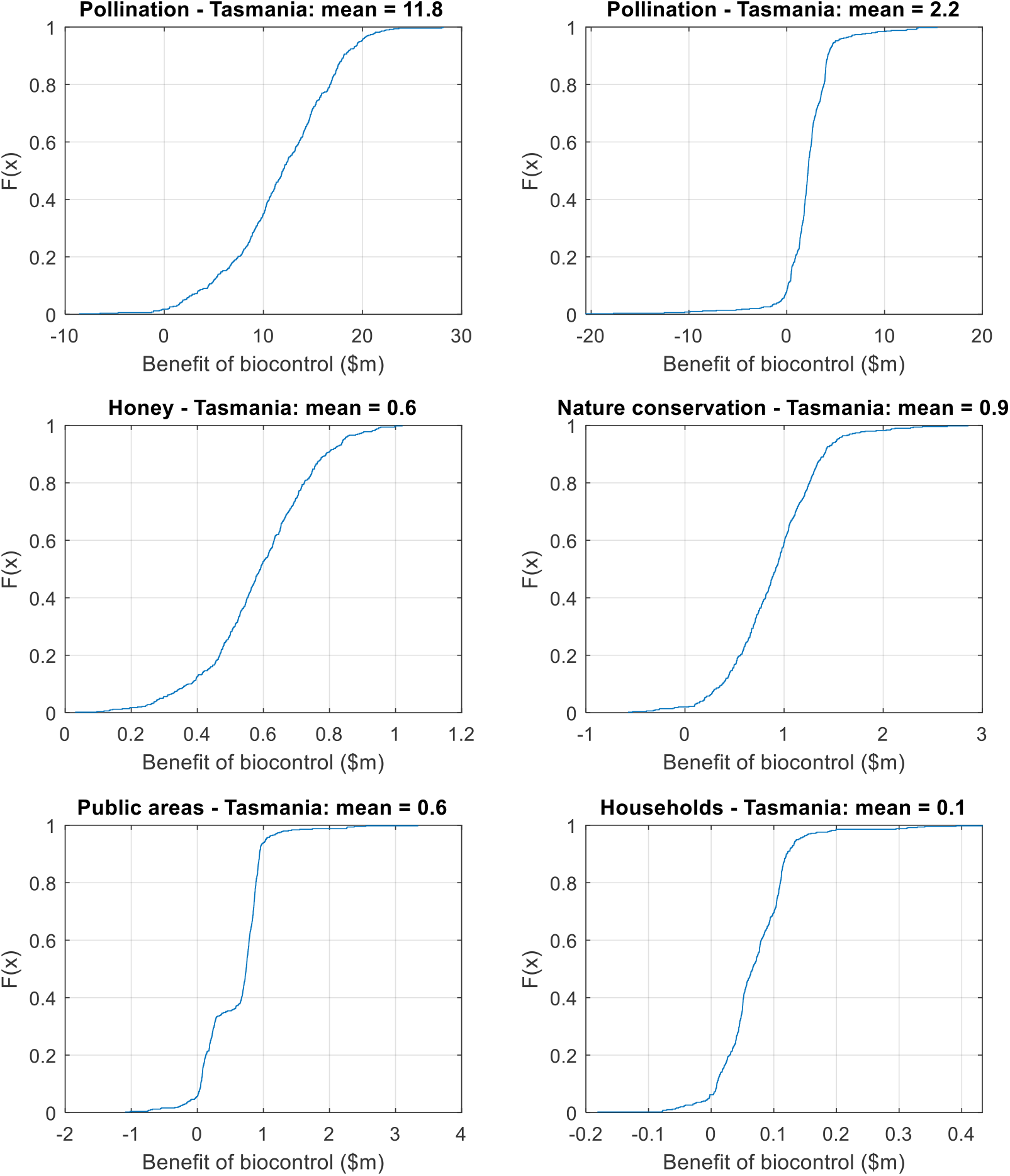
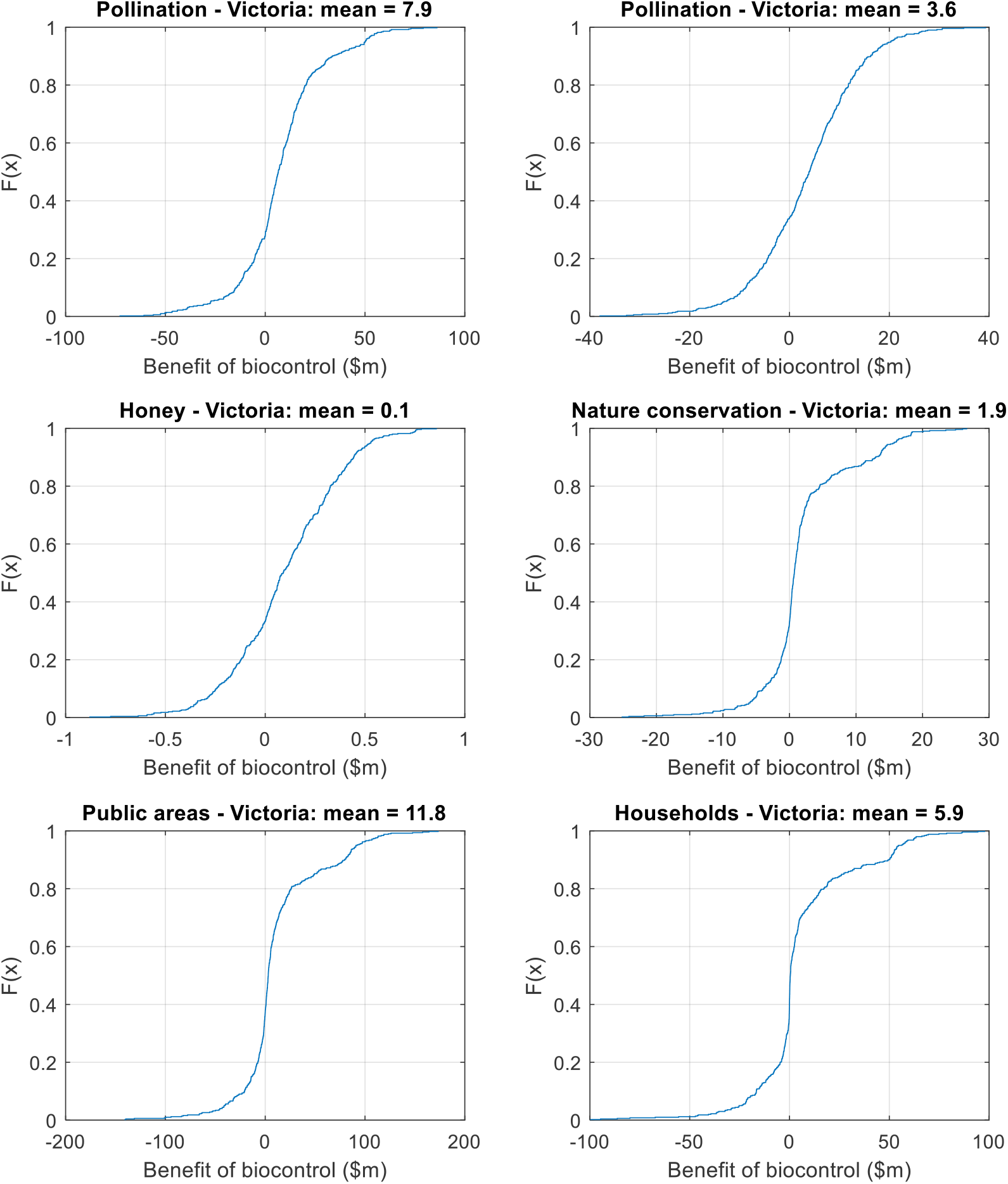

## Footnotes

1 Also known as the German wasp and German yellowjacket

2 Wasps are thought responsible for some traffic accidents in New Zealand, particularly in the summer time when motorists are more likely to be driving their cars with their windows open. The value of accidents attributable to wasps is estimated using the average social cost per vehicle crash.

3 Atlas of Living Australia occurrence download at https://biocache.ala.org.au/occurrences/search?q=qid:1590909475236 accessed on Sun May 31 17:18:47 AEST 2020.

4 ABARES (2016). Land use of Australia 2010–11, Available from https://www.agriculture.gov.au/abares/aclump/land-use/land-use-of-australia-2010-11

5 ABS (2019). 7503.0 - Value of Agricultural Commodities Produced, Australia, 2017-18. Available from https://www.abs.gov.au/AUSSTATS/abs@.nsf/DetailsPage/7503.02017-18?OpenDocument

6 ABS (2019). 7121.0 - Agricultural Commodities, Australia, 2017-18; Available from https://www.abs.gov.au/AUSSTATS/abs@.nsf/DetailsPage/7121.02017-18?OpenDocument

7 ABS (2017) 2016 Census Community Profiles. Available from https://quickstats.censusdata.abs.gov.au/census_services/getproduct/census/2016/communityprofile/036?opendocument

8 Jim Bariesheff, personal communication, June 29, 2020.

1 MNL estimates of Kerr and Sharp (2008) used for both Nelson and Christchurch.

2 WTP estimates from the ‘unlabelled’ CE design in Rolfe and Windle (2014).

3 https://www.rba.gov.au/inflation/measures-cpi.html. We apply equation 1 using the average CPI in 2009 and the first quarter CPI in 2020.

4 https://datapacks.censusdata.abs.gov.au/datapacks/

5 https://www.agriculture.gov.au/abares/aclump/land-use/data-download

6 https://www.pc.gov.au/research/supporting/cost-benefit-discount

